# Convergent evolution of extreme snout elongation in deep-sea fishes reveals constrained trajectories

**DOI:** 10.64898/2026.09.24.754272

**Authors:** Zach Heiple, Katherine E. Bemis, James C. Tyler, Aintzane Santaquiteria, Fernando Meléndez-Vazquez, Casey Lee, Kyoji Fujiwara, Masanori Nakae, Hsuan-Ching Ho, John J. Pogonoski, William T. White, Mark W. Westneat, Ricardo Betancur-R., Dahiana Arcila

## Abstract

Phenotypic extremes are scattered across the Tree of Life, suggesting that evolutionary constraints often limit their emergence. Yet striking departures from standard body plans, such as elongate snouts, have evolved convergently across deep time. We investigate extreme morphologies in two distantly related deep-sea fish clades: spikefishes (pufferfish relatives) and snipefishes (seahorse relatives). Using spikefishes as a study system, including all 24 living and fossil species, we integrated genome-wide phylogenomic data with a fossil and extant morphological character matrix, skull shape and dentition analyses from 64 CT scans, phenotypic and divergence matrices from >200 X-rays, functional manipulations of freshly collected and cleared-and-stained specimens, and formal convergence test across 300 percomorph species. We show that snout elongation evolved independently twice within spikefishes, reaching intermediate and extreme conditions along one predictable, ordered trajectory of additive skeletal rearrangements. Snout bones form a highly integrated, semi-autonomous module that channels evolutionary change along intrinsic lines of least resistance, coinciding with accelerated cranial evolution and expanded morphospaces in a clade with enhanced cranial mobility. Snipefishes converge on the same body plan through largely non-homologous elements and distinct linkage mechanics, except for near-identical vomer convergence. Extreme morphologies can emerge when phenotypic integration aligns with functional innovation, guiding evolution along repeatable trajectories.

## Main

Morphological extremes are rare across the Tree of Life, suggesting that stabilizing selection and evolutionary constraints often limit the emergence or persistence of highly divergent phenotypes^1–3^. Nevertheless, iconic examples, such as the elongated necks of giraffes^4,5^, the bioluminescent lures of deep-sea anglerfishes^6^, and the specialized feeding mechanisms of carnivorous plants^7,8^, highlight the power of evolution to produce dramatic adaptations. These derived phenotypes are especially intriguing because they challenge conventional ideas about trade-offs that govern relationships between form and function^9–12^. Although extreme morphologies are uncommon, they do emerge—and in some cases, repeatedly—across distantly related lineages, suggesting that under specific ecological or developmental conditions, evolution may converge on atypical outcomes^4–7,13–16^. Despite their potential to illuminate key evolutionary principles, the origins and trajectories of morphological extremes remain poorly understood.

Extreme snout elongation is a widespread trait across the Tree of Life, found in diverse taxa, such as elephants, weevils, and fishes ^17–22^. In fishes, elongated snouts enable novel feeding strategies by increasing reach, improving strike speed, and, in some cases, augmenting suction dynamics^22–27^ and likely create functional consequences in traits such as tooth or jaw morphologies. Even so, this repeated evolution underscores the adaptive potential of snout elongation, yet the anatomical structures contributing to elongation vary widely, involving extensions of exoskeletons, soft tissue, or skeletal elements^17,20,28^. This diversity complicates comparisons of elongation mechanisms across deep evolutionary timescales. In vertebrates, elongating the snout often requires significant remodeling of cranial bones. Because the skull is a highly integrated structure, with strong developmental and functional interconnections among its components, modifications in one region can have cascading effects across others. Indeed, many studies have found high levels of evolutionary integration in the skull, reflecting its coordinated architecture^29–34^. Lineages that evolve extreme cranial modifications despite this integration therefore provide powerful systems for investigating how extreme traits arise.

Because traits are developmentally, functionally, and genetically linked, evolutionary change is often biased towards particular directions in phenotypic space^35–41^. Schluter formalized this idea in his prediction that adaptive divergence is biased along intrinsic axes of trait covariance, commonly referred to as genetic “lines of least resistance” (LLRs)^40^. Although genetic covariance is rarely measured in non-model organisms, empirical and theoretical work suggests that phenotypic covariance can provide a useful approximation for identifying these dominant axes of variation^36,40,42^. In this context, extreme phenotypes that evolve along LLRs may not be difficult to generate, but are instead rare because they require a specific alignment between intrinsic trait covariance and ecological selection. Moreover, patterns of integration measured in snout-associated traits or bones (e.g., vomer, metapterygoid, palatine) across species provide a useful framework for understanding how snout bones have coevolved^43^. If integration is concentrated within the snout, this region may behave as a semi-autonomous morphological domain, allowing coordinated structural change with limited correlated effects on the rest of the cranium^43,44^. Repeated snout elongation could then reflect selection acting along an existing axis of intrinsic variation.

Among fishes, snout elongation has evolved numerous times, often through the extension of jaw bones, as seen in butterflyfishes^19^, wrasses^45^, and elephantfishes^46^. In contrast, extreme tubular snout elongation, defined here as >30% standard length via non-jaw elements and forming a bony, tube-like structure, is known only from two distantly related lineages: spikefishes and snipefishes^21,28,47,48^. This threshold distinguishes these taxa from the shorter, snouts of seahorses and pipefishes, that possess similar snout elongation without achieving comparable snout-to-body size ratios^47^. Spikefishes (Tetraodontiformes) are a deep-sea mesopelagic clade that ranges from relatively typical body forms to species with highly elongated, tubular snouts, in some cases comprising nearly 50% of standard length^21,48^. Snipefishes, by comparison, are Syngnathiformes, a group that also includes long-snouted taxa, such as seahorses, pipefishes, and trumpetfishes^49^. Most syngnathiforms have a highly modified feeding mechanism known as pivot feeding, which combines long snouts, head rotation, and buccal expansion to produce rapid strikes and enhanced suction dynamics^22–27,50,51^. Despite their distinct evolutionary histories, these groups are frequently misidentified in museum collections, including a documented case in which a sequence, originally identified and provided as a spikefish, was later shown to correspond to a snipefish^52^, reflecting their remarkable external convergence. Both groups share striking similarities in body shape, dorsal spines, and elongate snouts, and are broadly distributed with overlapping habitat preferences, including underwater seamounts and continental shelves and slopes^53,54^. Yet, the extent and anatomical basis of this convergence have not been rigorously quantified.

Here, we investigate the evolutionary dynamics shaping extreme snout elongation by focusing on the unique, tubular snouts of spikefishes and their striking convergence with snipefishes. We first estimate comprehensive, tip-dated phylogenies for spikefishes by integrating genome-wide data with a newly coded 80-character morphological matrix that includes both known fossil taxa and all extant species. Using this phylogenetic framework, we test two complementary hypotheses regarding snout evolution in spikefishes: (1) that morphological integration channels snout evolution along a predictable, constrained trajectory; and (2) that snout elongation is associated with diversification in tooth morphologies, suggesting functional differentiation in feeding strategies. To test this, we examine patterns of morphological evolution through (i) three-dimensional geometric morphometrics to quantify patterns of evolutionary integration and modularity across the skull, (ii) quantitative linear morphometrics to assess within-species variation and approximate lines of least resistance, and (iii) dentition-related measurements to explore the relationship between tooth morphology and snout elongation. Leveraging the morphological similarities between snipefishes and spikefishes, we further investigate whether convergence in snout form arises via homologous structures, analogous functional adaptations, or distinct anatomical solutions. Together, this study addresses a central question in evolutionary biology: whether extreme phenotypes reflect historically contingent departures from constraint or predictable outcomes shaped by intrinsic developmental and functional architecture.

### Time tree of living and fossil spikefishes reveals multiple origins of snout elongation

To establish the evolutionary context in which extreme snout elongation arose, we inferred phylogenetic relationships and divergence times for all 24 extant and two fossil spikefish species using Bayesian tip-dating. Analyses integrated 1,108 exon markers and a newly compiled 80-character morphological matrix (Figure 1; Tables S1-S6; Datasets 1 & 2, see Materials and Methods). To account for phylogenetic uncertainty in downstream comparative analyses, we applied the Fossilized Birth-Death (FBD) model across four non-overlapping genomic subsets, yielding stable relationships, with minor variation in a clade containing species with short and long snouts (*Tydemania*, *Paratriacanthodes,* and *Bathyphylax*). Maximum-likelihood and coalescent-based methods, using either molecular or combined molecular and morphological data, produced largely congruent relationships, whereas parsimony and morphology-only analyses showed greater topological variation (Appendix 1).

**Fig. 1:**
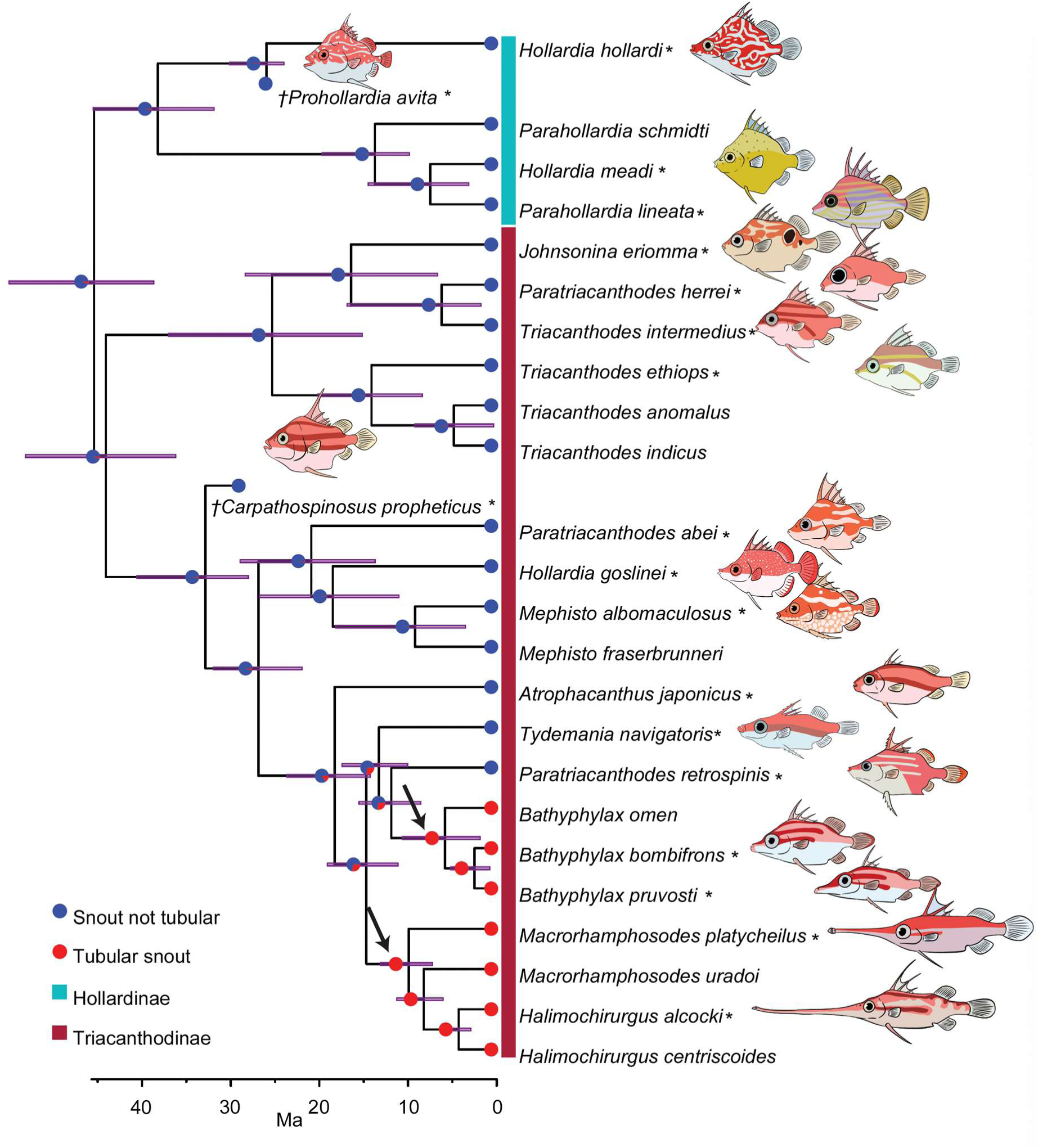
Time-calibrated spikefish phylogeny reveals two independent origins of snout elongation. Topology is derived from Bayesian tip-dating using the Fossilized Birth-Death (FBD) model implemented in MrBayes, integrating genomic data, a newly coded morphological matrix, and fossil taxa (†) as terminal tips. Purple bars indicate 95% highest posterior density (HPD) intervals for node ages. Ancestral state reconstruction of tubular snout evolution, defined by lateral closure space within the snout, is shown as pie charts on tree nodes. Asterisks by species names indicate taxa for which illustrative drawings are shown.

Divergence time estimates were sensitive to the inclusion of spikefishes sister family Triacanthidae (tripodfishes) and clock-model choice, reflecting pronounced rate heterogeneity and long subtending branches separating the two families (Appendix 1). When tripodfishes were included, log-likelihood comparisons favored an uncorrelated IGR clock over an autocorrelated TK02 model, estimating a mean crown age of ∼40 Ma (95% highest posterior density [HPD]: 63–31 Ma) for spikefishes. Excluding tripodfishes reversed this preference, with the TK02 model outperforming IGR and yielding an older crown estimate of ∼45 Ma (95% HPD: 55–39 Ma), consistent with previous studies^55,56^ (Tables S7 and S8). These results highlight known sensitivities of Bayesian dating to taxon sampling and rate heterogeneity^56,57^. To minimize downstream biases, we based comparative analyses on divergence-time estimates excluding tripodfishes.

Our phylogenetic analyses reveal a more complex history of snout elongation in spikefishes than previously inferred^52^, and identify the mid-to late Miocene (15–6 Ma) as a key interval of morphological innovation. Whereas earlier analyses placed the long-snouted genera *Halimochirurgus* and *Macrorhamphosodes* in separate clades^52^, our expanded genomic and total-evidence framework supports a paraphyletic *Macrorhamphosodes* that includes *Halimochirurgus*, consistent with a single origin of extreme snout elongation. Fossil taxa resolve within expected subfamilial placements: †*Prohollardia avita* as sister to *Hollardia* (Hollardinae), and †*Carpathospinosus propheticus* nested within Triacanthodinae^58^ along the stem between two major subclade radiations. Notably, both fossils possess short snouts, suggesting that extreme snout elongation represents a relatively recent innovation within spikefishes (Figure 1).

### Additive morphological changes underly extreme snout elongation

Having established the phylogenetic framework, we next examined the anatomical basis of transitions towards snout elongation. The placement of *Bathyphylax pruvosti*—an intermediate-snout species nested among both short- and extremely long-snouted taxa—suggests either a single origin of snout elongation followed by two independent reversals, or two independent gains of elongation. Model-averaged ancestral state reconstructions of tubular-snout morphology (defined by lateral closure of the snout bones; Figures 2B and 2C) consistently supported two independent origins of snout elongation across equal-rates, symmetric, and asymmetric models (Figure 1 and S1). Moreover, we assessed whether the apparent convergence between *Bathyphylax* and extremely long-snouted spikefishes may reflect hemiplasy, i.e., the possibility that snout length was ancestrally polymorphic and only retained in some lineages. We subsampled the 1,108 gene trees to include only those containing the taxa needed to test one or two independent gains of snout elongation (*Bathyphylax*, *Macrorhamphosodes*/*Halimochirurgus*, *Paratriacanthodes*, and *Tydemania*). Among the 623 trees matching this criterion, only one tree (0.16%) showed a sister relationship of *Bathyphylax* with an extremely long-snouted taxon (Figure S2). Further, using the coalescent-simulation method HeiST^59^, with parameter estimates from our coalescent-based tree and three mutation-rate regimes, we found modest support (52–60%) for two independent origins over a single origin in all scenarios (Figure S3), indicating that hemiplasy is not strongly supported by these data. Finally, these results are consistent with snout-specific morphological characters that distinguish *Bathyphylax* and support an independent, convergently evolved origin of its snout elongation (Table S9).

**Fig. 2:**
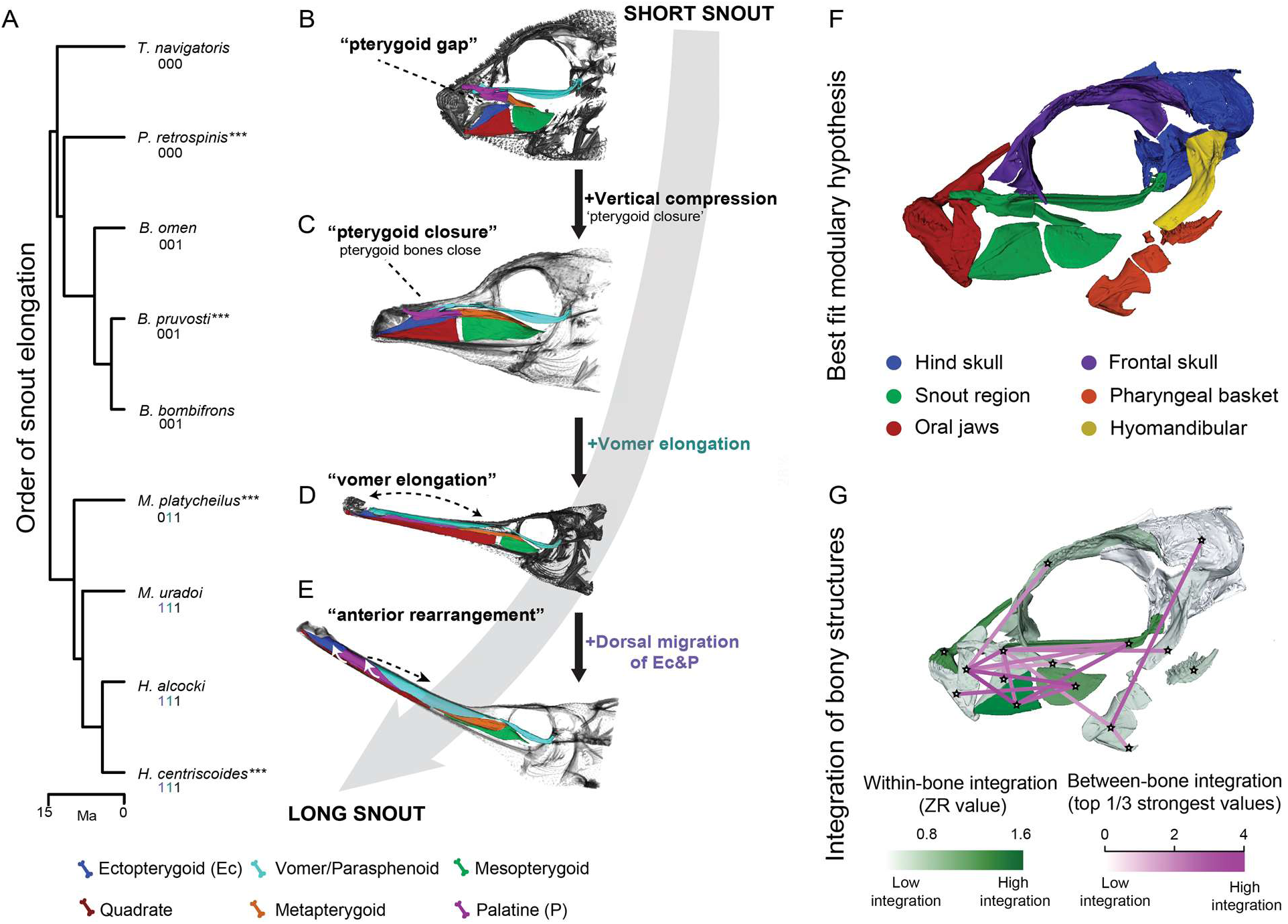
Snout elongation in spikefishes proceeds through stepwise anatomical change. **A**, Phylogenetic ordering of snout elongation, from short- to long-snouted species. Numbers below species names represent the presence (1) or absence (0) of three additive binary traits, and the best-supported model of character accumulation inferred using HyperMk. **B**, Pterygoid gap, the lateral spaces between snout bones in short-snouted spikefishes. **C**, Pterygoid closure, the closing of gaps between the ectopterygoid (blue) and palatine (purple), and between the metapterygoid (orange) and mesopterygoid (green). **D**, Vomer elongation, defined as the elongation of the vomer-parasphenoid complex (light blue). **E**, Anterior rearrangement of the ectopterygoid (blue) and palatine (purple). The quadrate (red) elongates progressively across all steps. Triple asterisks denote species illustrated to the right of the phylogeny. **F**, Best-supported modularity hypothesis (Snout3), based on covariance ratios (CRs) and effect sizes estimated in *geomorph*. Modules correspond to approximate bony regions; specifically, Hind skull = (Supraoccipital, epiotics, pterotic, and basioccipital), Snout region = (palatine, ectopterygoid, quadrate, mesopterygoid, metapterygoid, parasphenoid, and vomer), Oral jaws = (premaxilla, dentary, articular, and maxilla), Frontal Skull = (frontal, lateral and ethmoids), Pharyngeal basket = (ceratohyal, epihyal, urohyal, and pharyngeal jaws), and Hyomandibular = (hyomandibular) **G**, Degree of evolutionary integration for individual cranial bones in spikefish. Nodes of the network (stars) are placed on each bone-level module landmarked. Dark green shading indicates high within-bone integration, and pink lines represent the magnitude of between-bone integration, with darker lines corresponding to stronger correlations.

Next, we examined whether our discrete snout states could be recovered from continuous snout-length data using Ornstein-Uhlenbeck (OU) models under Bayesian and maximum-likelihood approaches. Under the maximum-likelihood framework, which requires *a priori* regime hypotheses, an OU3 model provided the strongest support for distinct adaptive snout regimes consistent with a two-shift hypothesis, in which intermediate-snouted *Bathyphylax* and extremely long-snouted taxa represent separate regimes. This model was favored over BM1, OU1, OU2, and an alternative OU3 hypothesis in which the remaining relatives of *Bathyphylax* were also treated as intermediate. Using 100 stochastic character maps, we further showed that both coding scenarios consistently favored OU3 over OU2 in 98–100% of maps—that is, two shifts over one— providing additional support for two independent gains (Figure S4). We next conducted an OU-shift search using the reversible-jump MCMC (rjMCMC)^60^ method across four shift-count priors, without a priori regime coding. All analyses identified the extremely long-snouted spikefishes with strong support, and the branches leading to *B. pruvosti* and the genus *Bathyphylax* reproducibly carried the next-highest posterior probabilities across all chains and priors (7–25%) (Figure S5), with absolute support scaling with the prior mean but rank order remaining stable. Such modest support is expected for a small-magnitude shift in a species-poor clade, where reversible-jump OU methods have limited power^61,62^.These results converge with our observations based on osteological characters, and we interpret them as support for treating *Bathyphylax* as a distinct snout regime in downstream analyses (Figures S4 and S5).

These results motivated a more detailed test of whether extreme snouts arose through discrete anatomical innovations accumulated in a predictable order. Using our morphological character matrix (Table S6; Dataset 2), we identified three additive binary traits central to snout elongation: (i) pterygoid gap defined by lateral spaces between snout bones in short-snouted spikefishes (Figure 2B); pterygoid closure, in which the pterygoid gap closes to produce a more tubular structure (Figures 2B, 2C); (ii) vomer elongation, in which the vomer bone extends anteriorly, lengthening the snout dorsally and marking an intermediate stage of elongation (Figures 2C, 2D); and (iii) anterior rearrangement, in which forward displacement of the ectopterygoid and palatine fully extends the tube-like snout, achieving extreme projection (Figures 2D, 2E). Evolutionary models for these additive binary traits consistently favored an irreversible pathway (AICw ≈ 1.0), in which traits were gained but not lost (Figures 2A–2E and S1). Moreover, simulations based on our empirical trees and parameter estimates (see Extended Methods and Figure S6) revealed that the model was unlikely to produce well-supported false positives (1.6–4.0% with ΔAIC ≥ 2) and never produced a ΔAIC equivalent to or greater than that observed empirically. Finally, when data were simulated under the known trajectory, the correct pathway was favored in all runs which converged (93.2–97.6% of simulations), and 66.0–74.0% showed significantly strong support (ΔAIC ≥ 2). These results define a well-supported pathway of additive, unidirectional steps from short to long snouts, notably capturing the convergent evolution of pterygoid closure, which also evolved in the intermediate-snouted *Bathyphylax*.

Because discrete traits can oversimplify continuous variation^63^, we further evaluated whether continuous bone-shape change was consistent with this ordered trajectory using two-block partial least squares (2BPLS) analyses across pairs of snout bones (Figure S7; Dataset 3). For each bone pair, we assessed whether species were ordered along regression axes in a manner consistent with our stepwise model. For example, we predicted that genus *Bathyphylax*, which exhibits pterygoid closure and an intermediate snout length, would occupy a midpoint along the short-to-long snout continuum. Using both local and global Procrustes superimposition schemes (LPS and GPS; see below), members of *Bathyphylax* were consistently resolved as intermediate in snout-bone shape, particularly among adjacent elongated elements (Figure S7). All 2BPLS tests showed significant integration between paired snout bones (Table S10). In contrast, *M. platycheilus*, which is inferred as a later step in the discrete model (Figures 2A–2E), was not consistently recovered as intermediate in 2BPLS space. This likely reflects the fact that 2BPLS captures covariation between trait blocks rather than fine-scale evolutionary ordering along a morphological continuum. Nonetheless, multiple independent discrete characters support its inferred position (Tables S6 and S11), and qualitative examination of mapped shape change is consistent with this interpretation (Figure S8). Overall, spikefish snout evolution follows a continuous trajectory of gradual, intermediate shape change, with parallel transformations extending beyond discrete trait transitions.

### Snout elongation evolves along lines of least resistance

Extreme snout elongation may arise either from intrinsic patterns of variation that bias evolutionary change along particular axes or from external ecological pressures that favor longer snouts. Throughout, we treat lines of least resistance as an empirically observable bias in phenotypic variation; therefore, observed alignment is compatible with channeling selective pressures, but does not by itself demonstrate underlying deterministic constraint or causal mechanism. To test whether the direction of snout elongation was aligned with major axes of within-species phenotypic covariance, we quantified the alignment between the major axis of within species phenotypic variance (**p_max_**) and the primary axis of interspecific divergence (**d_max_**). Phenotypic variance-covariance (**P**) matrices estimated from six linear cranial measurements across two short-snouted species of *Parahollardia*, *P. lineata* (n = 82) and *P. schmidti* (n = 37) and two long-snouted *Halimochirurgus* species, *H. alcocki* (n = 41) and *H. centriscoides* (n = 48) (hereafter the “focal species”) were highly similar within genera but markedly different between them (Krzanowski subspace projections over 3–4 PC axes, ∼80% of variance; Figures 3A and S9; Table S12; Dataset 4). This pattern indicates conserved integration at shallow phylogenetic scales and substantial restructuring across deeper divergences. Because **P** matrices were highly similar within genera, we pooled species within each genus to increase sampling robustness. We then estimated a divergence (**D**) matrix for spikefishes following Simon et al.^39^, using phylogenetic independent contrasts to estimate evolutionary rates and trait coevolution.

**Fig. 3:**
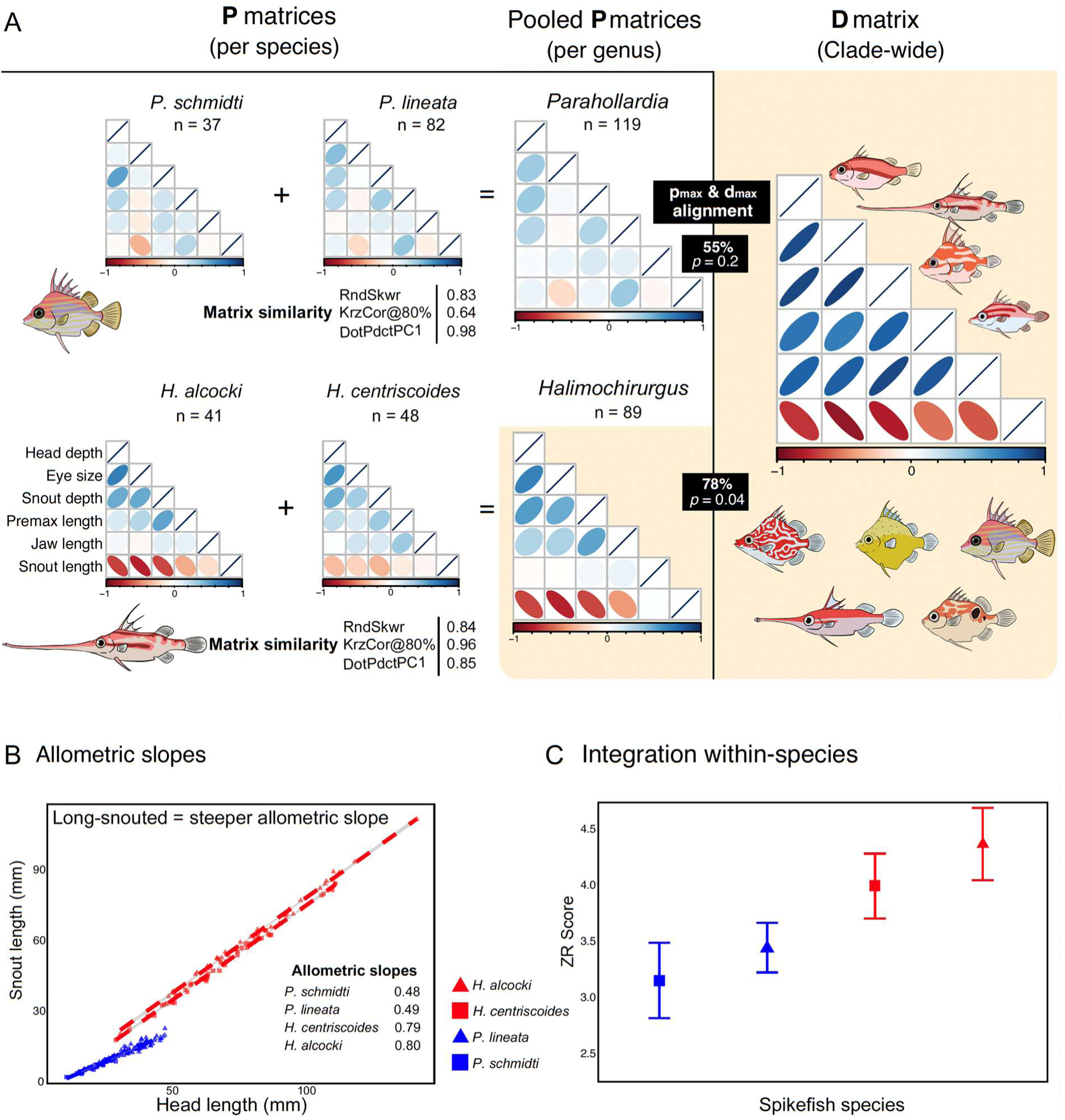
Phenotypic variance predicts macroevolutionary divergence in long-snouted species. Long-snouted spikefishes show close correspondence between within-species phenotypic variation and the dominant directions of evolutionary divergence across the clade. **A**, Within-species phenotypic variance–covariance (**P**) matrices for short-snouted (top) and long-snouted (bottom) taxa, visualized as trait correlation plots. Blue and red ellipses indicate positive and negative correlations, respectively, with shading intensity reflecting correlation strength. Matrix similarity was quantified using Random Skewers, Krzanowski projections (80% cumulative PC variance), and PC1 vector alignment. Krzanowski projections for within-genus comparisons included 3 and 4 PC axes for *Parahollardia* and *Halimochirurgus*, respectively. Species-level **P** matrices were pooled at the genus level when statistically similar. The divergence (**D**) matrix summarizes trait covariation across the clade and closely matches patterns observed in long-snouted taxa. For *Halimochirurgus* and *Parahollardia*, the major axes of within-species variance (***p***ₘₐₓ) and interspecific divergence (**d**ₘₐₓ) are shown. **B**, Linear regressions of head length versus snout length for two long- and two short-snouted species; allometric slopes are reported in the inset (red, long-snouted; blue, short-snouted). **C**, Within-species integration quantified using eigenvalue dispersion (ZR), where ZR = 0 indicates no integration.

To address possible biases associated with snout length, we conducted all alignment tests under size-correction schemes that either included (+SnL) or excluded (-SnL) the snout, and assessed significance against a phylogenetic null generated by simulating trait evolution under multivariate Brownian motion on our phylogeny (following Simon et al.^39^; Extended Methods). Alignment between **p_max_** and **d_max_** was significant for *H. centriscoides* under both size-correction protocols (23.8°, *p* = 0.002; 37.6°, *p* = 0.032) and for pooled *Halimochirurgus* with snout length included (38.8°, *p* = 0.033), whereas *H. alcocki* (44.1°, *p* = 0.067; 53.9°, *p* = 0.155) and pooled *Halimochirurgus* under the more conservative -SnL protocol (49.5°, *p* = 0.099) fell just short of the one-tailed 5% criterion (Figures 3A, S10, and S11; Table S13). Critically, the two genera did not overlap under either protocol: every *Halimochirurgus* comparison fell between 23.8° and 53.9° (*p* = 0.002–0.155), whereas every *Parahollardia* comparison fell between 55.4° and 85.7° (*p* = 0.175–0.863; Table S13). Consistent with this pattern, long-snouted species displayed allometric slopes that were 1.6-fold steeper than short-snouted species (Figure 3B), indicating amplified divergence along intrinsic axes of variation. To further evaluate differences in covariance structure, we compared the strength of trait integration across species using eigen dispersion values (EDV)^64,65^, which assess covariation along leading eigenvectors and are directly comparable across datasets. Across all size-correction schemes, long-snouted species consistently exhibited stronger phenotypic integration than short-snouted species (*p* < 0.0001) (Figure 3C). These results demonstrate that divergence in long-snouted spikefishes is significantly aligned with the dominant axis of within-species phenotypic variance. This pattern is consistent with Schluter’s prediction that evolutionary divergence is biased toward intrinsic axes of trait covariance, rather than occurring independently of them^40^. Importantly, we do not interpret this alignment as evidence that integration drives evolutionary change, but rather it suggests that selection acting on available variation can more readily produce divergence when intrinsic covariance and functional demands are aligned.

### Evolutionary correlations support integration-guided snout evolution

Spikefish skulls show striking diversity in the snout bones (Figure 2), yet overall craniofacial architecture remains largely conserved, suggesting that the snout evolved as a semi-independent unit. To test whether certain cranial regions tend to evolve more synchronously, we quantified evolutionary modularity using covariance ratios (CRs)^66^ using our 3D geometric morphometric dataset (Dataset 3). CRs compare covariation within versus between predefined skull regions, with values <1.0 indicating semi-independent evolution. We tested 11 hypotheses: eight *a priori* and three ‘snout-separate’ models that emphasized modular snout evolution (Figure S12). Because long-snouted species distorted global Procrustes superimpositions (GPS), an artifact known as the “Pinocchio effect”^67^—a consequence of disparate landmarks biasing how remaining landmarks are aligned—while local Procrustes superimpositions (LPS) mitigated this artifact by using a reference shape^68^. CR ratios for both GPS and LPS supported a six-module hypothesis (snout, oral jaws, pharyngeal basket, frontal skull, hyomandibular, and hind skull) and a “fully modular” hypothesis in which individual bones represented unique modules (Figure 2F; Table S14). Given LPS robustness to preservation artifacts and mobile structures, we used LPS coordinates for all downstream analyses.

We further examined how integration was distributed across spikefish skulls using a second round of EDV tests. In this context, evolutionary integration was measured from landmark covariation of clade-wide Procrustes alignments of individual bones or modules (Tables S15 and S16). The snout module exhibited significantly higher integration than the pharyngeals, hind skull, and oral jaws (*p* < 0.05), marginally higher than the hyomandibular (*p* = 0.07), but not significantly different from the face (*p* = 0.46). Within-bone integration of snout elements was, on average, 1.22× higher than that of non-snout bones (Figure 2G), although this difference was not significant (*p* = 0.11); however, the vomer, quadrate, and premaxilla, which are all highly modified in long-snouted spikefishes, showed significantly greater within-bone integration compared with most other cranial elements (*p* < 0.05; Figures 2G, S13, and S14; Table S15). Collectively, these findings suggest that spikefish snout bones co-evolve as a semi-autonomous unit, even in the face of dramatic phenotypic change.

We next assessed between-structure integration using phylogenetic 2BPLS regressions. Of 105 pairwise bone comparisons, 58% showed significant covariation, with 51% remaining robust after FDR correction (Figures 2C and S13; Table S17). Integration was, on average, 1.45× higher in comparisons involving snout bones, although this trend was not statistically significant (*p* = 0.07). Similarly, slightly over one-half (8) of the between-module tests (15) were also significant and robust to FDR correction (Table S18). We also found a moderate correlation of inter-bone integration with per-bone morphological disparity (R² = 0.29, *p* = 0.02), although this association was less clear after FDR correction (*p* = 0.09). Other analyses of within-bone or module-level variation yielded no significant correlations (Figure S14; Tables S19 and S20). Whereas the density of correlations among snout bones may help explain the repeated evolution of extreme morphologies, we caution against interpreting integration as a causal driver of phenotypic divergences, as this has led to confusion in the literature^43^. Instead, these findings— alongside our observations of phenotypic integration, alignment along lines of least resistance, and ordered anatomical transitions of snout evolution—are consistent with the view that fine-scale integration shaped evolutionary trajectories of bone evolution, facilitating repeated transitions toward phenotypic extremes.

### Tooth and skull morphologies diversify with a shift in feeding modality

Because dental traits encode information about prey capture and processing, they provide a functional window into trophic differentiation when direct diet data are limited^69^, as is common for deep-sea fishes^70^. We therefore quantified 22 dental traits as proxies for feeding strategy and examined their relationship with snout elongation and cranial architecture (Table S21; Dataset 5). To minimize biases introduced by extreme snout length, we focused primarily on size-corrected datasets excluding snout length from head-length calculations (-SnL), confirming that overall patterns were robust across correction schemes (Figure S15). Dental morphospace analyses revealed that much of spikefish dental diversity is concentrated within a subclade of nine species characterized by having a cranial elevation coupled buccal expansion mechanism (hereafter, the high-head-mobility clade; Figures S16 and S17; Datasets 5 and 6), a feature common among suction feeding fishes^23,26,71–74^, that is absent in all other spikefishes. Two anatomical traits indicate head mobility: a hinge-like joint that enables dorsally directed cranial rotation and variable positioning of bones in the buccal cavity^23,75,76^, features that are otherwise ossified or absent in the remaining species. The high-head-mobility clade spans the full range of snout lengths, from short to extremely elongated, and includes both independent origins of snout elongation, consistent with the emergence of novel ecological and functional demands. Both -SnL and +SnL showed similar morphospatial patterns, but trait loadings differed across schemes (Figures 4D and S15; Tables S22 and S23). In -SnL morphospace, PC1 (39.7%) captured variation in mouth shape and upper-jaw dentition, isolating long-snouted taxa at the positive extreme. PC2 (25.6%) differentiated long-snouted species based on tooth dimensions and jaw-closing mechanical advantage, with low PC2 values corresponding to wall-like teeth in scale feeding *Macrorhamphosodes* and high values corresponding to tooth-reduced *Halimochirurgus* (Figures 4B, 4D).

**Fig. 4:**
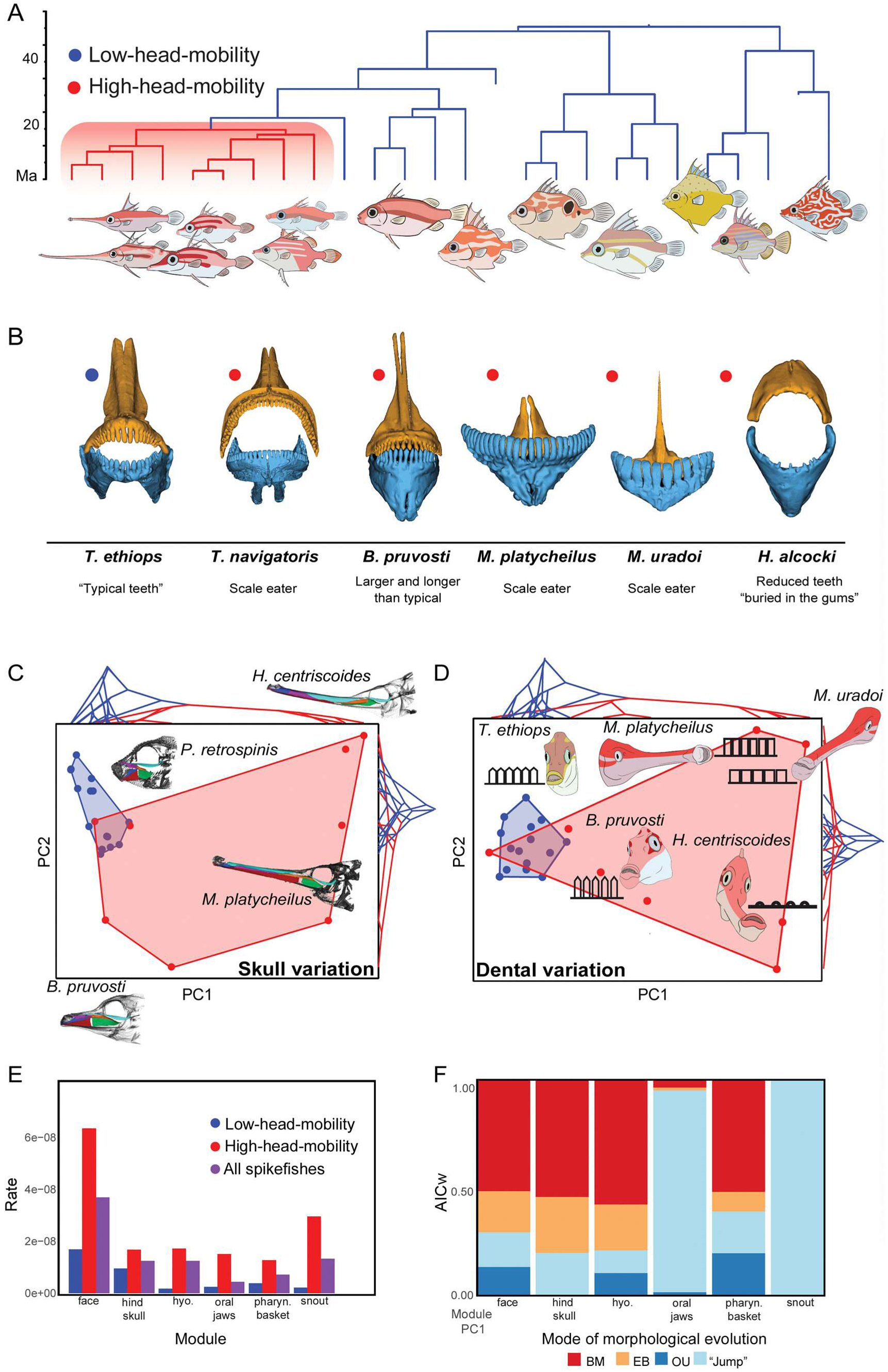
Cranial mobility coincides with expanded morphospaces and accelerated cranial evolution in spikefishes. **A**, Phylogeny of spikefishes with branches colored by the presence (red) or absence (blue) of elevated head mobility. **B**, Representative images of spikefish teeth comparing “typical teeth” of low-head-mobility spikefishes (left) with the range of dental morphologies observed in the high-head-mobility clade. **C**–**D**, Morphospaces of spikefish crania (left; all cranial modules combined) and dental batteries (right). Phylogenetic trees are projected as traitgrams onto PC1 and PC2 axes, with colors corresponding to head mobility states shown in panel A. **E**, Evolutionary rates estimated separately for high-head-mobility species, low-head-mobility species, and the full spikefish dataset, partitioned by cranial module. **F**, Evolutionary model-fitting results for individual cranial modules. Models tested include Brownian Motion (BM), Early Burst (EB), Ornstein-Uhlenbeck (OU), and the best-supported jump (Pulsed) model.

Across size-correction methods, clade-wide regressions revealed a general trend toward tooth reduction with increasing snout length; however, extremely long-snouted spikefishes frequently deviated from this relationship (Figure S18), indicating that extreme elongation is not simply accompanied by uniform dental simplification. Available dietary data support this interpretation: short-snouted spikefishes primarily consume hard benthic prey, whereas members of the high-head-mobility clade exhibit marked departures from this pattern, including two independently evolved scale-feeding lineages (*Tydemania* and *Macrorhamphosodes*)^21,77–79^ and taxa lacking clear evidence of hard-prey consumption *(Halimochirurgus*) (Table S24). The latter is supported by examination of 91 X-rayed *Halimochirurgus* specimens and one stomach-content sample that revealed no evidence of hard prey or scales (Figure S19), reinforcing the inference of a distinct feeding strategy.

Patterns of cranial shape evolution closely mirrored those observed in dentition. Short-snouted spikefishes occupied a restricted region of cranial morphospace, reflecting strong phenotypic conservatism, whereas the high-head-mobility clade exhibited a pronounced expansion into previously unoccupied regions regardless of +SnL or -SnL Procrustes schemes (Figures 4C and S20). PC1 (54.1% of variation) captured vomer elongation and pterygoid closure (Figures 2B and 2C), with long-snouted species exhibiting high PC1 scores. PC2 (6.98%) further differentiated long-snouted species based on variation in vomer elongation, maxillary configuration, and ethmoid shape, consistent with the ordered trajectory of snout evolution inferred from discrete and continuous analyses (Figures 2A–2E). Evolutionary rate analyses revealed a strong acceleration of cranial evolution within the high-head-mobility clade, with cranial traits evolving 1.5–8.1× faster than in otherwise morphologically conserved spikefishes (Figure 4E). All in all, these results indicate that diversification of skull and tooth morphologies coincides with the appearance of cranial elevation, consistent with elevated head mobility reshaping the evolutionary and functional landscape of spikefish feeding.

### Snout evolution shows mixed signals of gradual and pulsed change

Spikefish evolution is characterized by sequential morphological innovations that bear directly on debates over gradualism versus punctuated equilibrium^80,81^. We tested the tempo and mode of cranial evolution by fitting evolutionary models ranging from Brownian Motion to jump (pulsed) processes^82^ to linear functional traits (Dataset 5) and module-level morphospace coordinates derived from geometric morphometrics (Dataset 3). Most linear traits favored gradual Brownian evolution, but snout traits, upper-jaw tooth metrics, and, in some cases, oral-jaw modules showed stronger support for jump models (Figures S21–S23), indicating that some anatomical transitions occurred as relatively discrete shifts. Next, we assessed the reliability of these inferences by simulating data on our phylogenies under the fitted pulsed models. Power to recover a pulsed model varied among traits and trees (24–91%): lower-jaw length and upper-jaw tooth region yielded consistently recoverable pulsed signatures, whereas most other traits varied across trees. Although a pulsed model was preferred for snout length in all empirical analyses, the fitted snout parameters produced recoverable pulsed models in only 24–50% of simulations (9–21% as the exact generating model), reflecting limited power at our sample size; we therefore regard pulsed snout evolution as suggestive rather than firmly established (Figure S24 and Table S25). We further quantified heterogeneity in evolutionary rates across cranial modules, noting that local Procrustes superimposition may conservatively estimate rates in the snout as bone position is lost when mapping landmarks to a reference^68^. Although the frontal skull module showed the fastest evolutionary rates, driven primarily by the orbital and ethmoid bones (Figure 4E,F), it was best supported by a Brownian model, indicating rapid but directionless divergence (an unconstrained random walk with no attraction toward a shared optimum) rather than change confined to a small region of morphospace. In contrast, the snout module retained high rates while showing stronger support for punctuated shifts into novel regions of morphospace (Figures 4E,F). Evidence of intermediate morphologies accompanied by large leaps in morphospace (Figures 4E,F, S7 and Tables S6, S10, and S11) suggests that macroevolutionary patterns that appear pulsed may in some cases arise from rapid but fundamentally gradual change, motivating alternative but non-exclusive fitness-landscape models. Support for this interpretation comes from ontogenetic variation in snout length observed in long-snouted spikefishes (Figure 3B; see also ref.^21^) and from the retention of typical (percoid) configurations of the snout and suspensorium in juvenile tetraodontiforms^83^, indicating that developmental trajectories may preserve access to intermediate phenotypes and provide a mechanistic basis for the evolution of extreme snout elongation.

### Convergence in functional trophic morphologies through deep time

Leveraging the striking similarities between long-snouted spikefishes (Tetraodontiformes) and snipefishes (Syngnathiformes), lineages that both inhabit seamounts and continental shelves yet diverged more than 100 million years ago^84^, we investigated whether their anatomical similarities reflect shared homology or non-homologous many-to-one mapping^85^ of distinct skeletal elements (Figure 5; Dataset 7). Comparisons of homologous snout bones revealed that, in both snipefishes and spikefishes (*Halimochirurgus* and *Macrorhamphosodes*), the vomer has independently evolved into an extremely elongated structure forming the dorsal roof of the snout, representing an exceptional case of structural and functional convergence. In contrast, the remaining cranial elements differ substantially between the two lineages, indicating that overall snout similarity cannot be explained by direct homology among corresponding snout elements. Instead, several bones occupy analogous positions and exhibit similar shapes, consistent with functional equivalence despite distinct anatomical identities (Figures 5, S25, and S26). Collectively, these results suggest that long snouts in spikefishes and snipefishes primarily arose via reshaped non-homologous structures, while also revealing an exceptional case of near-identical convergence in the vomer (Figure 5).

**Fig. 5:**
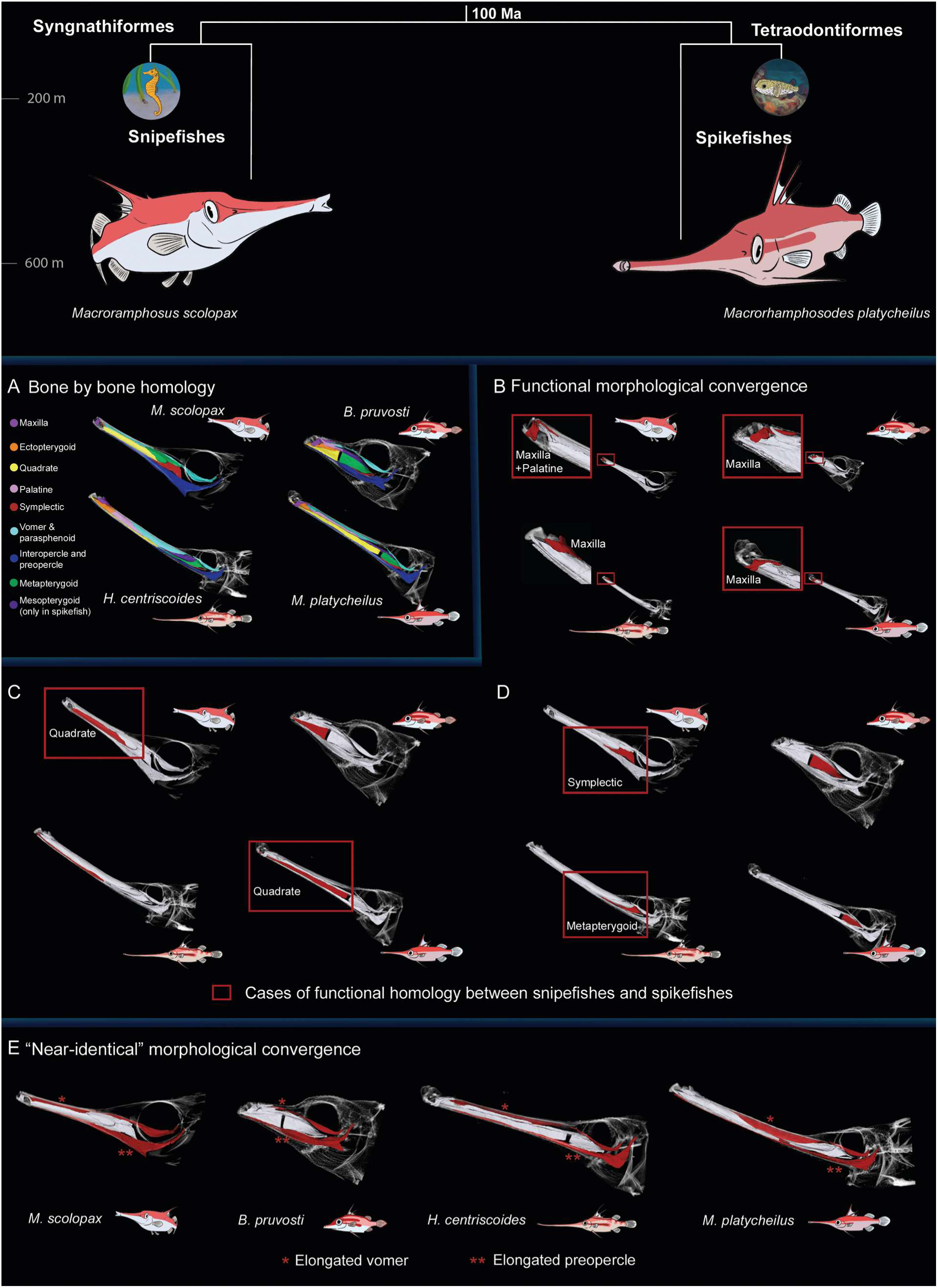
Extraordinary convergence between spikefishes and snipefishes despite 100 Ma divergence. Comparisons of snout morphology between the snipefish (*Macroramphosus scolopax*) and long-snouted spikefishes species, including *Bathyphylax pruvosti*, *Halimochirurgus centriscoides*, and *Macrorhamphosodes platycheilus*. Simplified phylogeny illustrating convergence in body plans, snouts, and habitat despite 100 Ma divergence (Top). **A**, Snout bone in snipefishes and spikefishes colored according to homologous structures. **B*–*D**, Convergence between snipefishes and spikefishes involving bones that are similarly shaped and positioned (functionally homologous). Red boxes indicate similarity between comparisons. **E**, Homologous bones that occupy similar shapes and functional roles in both spikefishes and snipefishes.

We formally tested for morphological convergence in a phylogenetic framework. We curated an expanded FishShapes^86^ dataset, capturing eight well-established body-shape measurements for 300 percomorph fishes (Dataset 9), and estimated Ct metrics^87,88^ under three workflows differing in size correction and morphospace construction (Tables S26 and S27; Extended Methods). Ct1 quantifies the proportion of the maximum ancestral morphospace distance between focal lineages that has been closed by subsequent evolution. Long-snouted spikefishes and snipefishes showed significant convergence under the two standard workflows (Ct1 = 0.72, *p* = 0.006; Ct1 = 0.74, *p* = 0.013; 1,000 simulations); under a third, doubly phylogenetically corrected workflow expected to be conservative, Ct1 did not reach significance (Ct1 = 0.41, *p* = 0.07; Ct3 = 0.13, *p* = 0.06), although Ct2 and Ct4 remained significant (*p* = 0.01 and *p* = 0.022). Because FishShapes does not include a snout-length measurement, these tests recover convergence from overall body form alone, independent of the trait that motivated the comparison, indicating that convergence between these groups extends beyond the snout itself. These pronounced anatomical similarities described raise the question whether spikefishes might have converged on a feeding mode analogous to the pivot feeding characteristics of syngnathiform fishes. Testing this hypothesis directly requires access to live specimens, which have been available for snipefishes, but largely inaccessible for deep-dwelling spikefishes. Syngnathiformes pivot feeding represents one of the most mechanically specialized feeding modes among fishes, relying on elongated snouts and four-bar linkage systems with bistable, torque-reversible properties that enable rapid prey capture^22–27,50,51^.

During the 2025 Coral Sea Frontier Expedition aboard the RV *Investigator*, we tested whether the head elevation and torque-reversal mechanisms described for snipefishes^27^ could be reproduced via manipulation of freshly collected spikefishes (Dataset 8). Consistent with our morphological observations of cranial elevation (Figure S16), manual manipulation revealed a four-bar linkage mechanism (*sensu* Muller et al.^23^) that couples head elevation with hyoid motion in all species examined, except *Triacanthodes ethiops*. Spikefishes among the high-head-mobility clade achieved head-elevation magnitudes comparable with or exceeding those reported for snipefishes^23,27^ (≥30°, Figure S27; Supplementary Videos S1–S3). Supplementing these observations with cleared and stained specimens, we expanded the phylogenetic context of four-bar linkage assessment to include seven of the nine remaining tetraodontiform families (Figure S28; Dataset 10). Of the seven non-spikefish families examined, none demonstrated a clear capacity to dorsally rotate the head, indicating that, while head elevation may be common among other fish lineages, the spikefish subclade is unique among tetraodontiforms in this regard and provides an additional layer of functional convergence with snipefishes (Figure S28). In snipefishes, torque-reversible mechanisms are supported by a limited range of head elevation when the suspensoria are constrained. Repeating this exercise with the freshly collected *Halimochirurgus centriscoides,* head elevation appeared restricted, with skin tightening around the suspensoria, and rotation seemingly occurring from the spine bending (as opposed to the cranial joint) whereas this strain was not observed when the suspensoria were unconstrained (Supplementary Videos S3 and S4). Additional observations during head elevation in *H. centriscoides* and adduction of the suspensoria combined with light posterior pressure along the epaxials in a freshly collected *Macrorhamphosodes uradoi* showed the hyoid transitioning abruptly from a horizontal to a vertical configuration (Supplementary videos S5 and S6), comparable to the abrupt shifts of hyoids in seahorses (previously described as ‘popping’ motion^89^). Attempts to reverse this configuration resulted in a brief delay followed by a quick snap back to the vertical state. These observations support that spikefishes may possess torque-reversible four-bar dynamics, although we caution that this interpretation is based on fresh specimen manipulations (and could not be recreated within day-old or cleared and stained specimens) rather than *in vivo* kinematics. Furthermore, spikefishes possess a smaller, restricted gill opening compared to “typical” teleost fishes, which has unknown impacts on the generation of negative pressure during buccal expansion.

Despite these strong mechanical similarities, hyoid anatomy in spikefishes differs substantially from that of snipefishes (Figures S17, S25, and S26). To further assess biomechanical similarity between the four-bar linkages of snipefishes and spikefishes relative to other taxa, we quantified kinematic transmission—a mechanical sensitivity metric describing hyoid rotation relative to input head rotation–using cleared and stained specimens (Dataset 10). Among the 23 species examined (spanning numerous teleost radiations), only spikefishes and snipefishes demonstrated a capacity to rotate the hyoid to a vertical configuration (Figures S17, S26, and S28), but nevertheless displayed markedly different KT values (1.4–1.9 in long-snouted spikefishes versus 5.2 in the snipefish; Figure S28 and Table S28), reinforcing the conclusion that similar feeding mechanics may be achieved through distinct anatomical configurations. Finally, specimens of *Macrorhamphosodes uradoi*, which exhibit either increasingly clockwise or counterclockwise rotation of the oral region with age, were able to restore a typical mouth posture when lateral displacement was applied to the hyoid (Supplementary Video S7), further suggesting a role of hyoid adaptations in spikefish evolution. In summary, while these similarities suggest spikefish-snipefish mechanical similarities, functional anatomies are morphologically distinct. Nevertheless, the findings reported herein positions spikefishes as a compelling system for future biomechanical research.

## Conclusions

Extreme morphologies are often interpreted as evolutionary outliers that arise through departures from constraint. Using a newly generated, time-calibrated total-evidence phylogeny that includes all known fossil and extant spikefish species, we show that intermediate and extreme snout morphologies evolved independently along a shared, ordered evolutionary trajectory. This shared trajectory is consistent with evolutionary contingency^90,91^and Schluter’s^40^ prediction that divergence is biased along intrinsic axes of phenotypic variation. Long-snouted species exhibit significantly stronger morphological integration and closer alignment between intrinsic covariance and phenotypic divergence than short-snouted taxa, suggesting that extreme phenotypes arise by following pre-existing axes of variation. Of course, our conclusions rest on **P** matrices estimated for the focal long-and short-snouted genera. Extending **P**-matrix estimation across the clade, which would require dense within-species sampling that is currently unavailable for most of these rare, deep-sea taxa, presents a natural way to refine the timing of this shift, for instance by pinpointing when the axis of least resistance rotated in intermediate forms such as *Bathyphylax.* Nevertheless, our macroevolutionary approach is consistent with this interpretation, evolutionary modularity support snout bones coevolving as a semi-autonomous unit, enabling coordinated and concentrated morphological change while largely preserving the broader cranium architecture. While intrinsic covariances help explain the emergence of extreme snout elongation, morphological integration itself is not an evolutionary force, but rather a means through which evolutionary change is facilitated. Identifying the genomic and developmental basis of snout elongation, for example, loci underlying craniofacial scaling, remains an important next step as population-level resources for these deep-sea taxa become available.

The channeling of these intrinsic axes into phenotypic diversity appears to depend on selection acting on available variation. In spikefishes, we find strong evidence for marked diversification in feeding strategies at both functional and morphological levels. Short-snouted spikefishes retain conserved biting-associated morphologies alongside limited cranial mobility, whereas a distinct clade (the ‘high-head-mobility clade’) exhibits cranial mobility coupled with buccal expansion. Cranial mobility coincides with the appearance of both intermediate and extreme snout elongation, increased dietary breadth, and pronounced diversification of dental morphologies. Collectively, these adaptations support the view that snout elongation in spikefishes is not merely a structural extreme, but, instead, represents a broader functional shift toward enhanced feeding versatility. In this light, snout elongation in spikefishes is consistent with a “spontaneous coincidence” scenario^7^, in which the alignment of cranial mobility, trait integration, and ecological opportunity enables selection on a novel functional configuration.

Of particular functional interest are the three scale-feeding spikefishes—all equipped with large, spongy lips and spatulate, wall-like teeth—suggesting either two independent gains or two losses of lepidophagy. Despite these oral-jaw similarities, *Tydemania* is short-snouted and lacks mouth “handedness” (i.e., twisting to the left or right side)^92,93^, like that seen in *Macrorhamphosodes*, which we note convergently evolved with respect to “lefty” and “righty” cichlids. Interestingly, Nakae and Sasaki^78^ determined that *M. uradoi* primarily feeds on scales from the caudal region of other fishes, hypothesizing that long snouts may position slow-swimming spikefishes in the blind spot of other fishes, optimize strike distance, and be consistent with cichlid scale feeders, which are rarely known to attack from the front. A comparable trophic novelty arose on a much faster timescale in the scale-eating pupfish *Cyprinodon desquamator* of San Salvador Island, which evolved enlarged oral jaws and modified strike kinematics within the past ∼10,000 years^94^; together with cichlids and spikefishes, these radiations suggest that lepidophagy, among the rarest of vertebrate trophic strategies, is repeatedly assembled through lineage-specific reconfigurations of the oral jaws. We expand this hypothesis with our observations that *Macrorhamphosodes* may induce mouth rotation with hyoid control, allowing for extreme scale-eating precision (Video S7) although *in vivo* kinematics are lacking to confirm this hypothesis. Nevertheless, the long-snouted *Halimochirurgus* appears to abandon biting strategies with highly reduced teeth, consequently converging upon the toothless seahorse and snipefish phenotype.

Morphological convergence through deep time highlights the power of selection in shaping evolutionary outcomes despite contingency. Comparisons with snipefishes reveal that similar cranial phenotypes can arise through distinct anatomical routes. Despite striking external similarity and overlapping ecological distributions, spikefishes and snipefishes evolve long snouts using primarily non-homologous skeletal elements, with the vomer representing a rare case of near-identical structural convergence. Moreover, our manipulations of fresh spikefish specimens suggest there are biomechanical similarities to the extreme pivot-feeding behavior of syngnathiforms, although live specimens are needed to confirm this hypothesis. These findings suggest functional demands can repeatedly shape comparable phenotypes even when developmental and anatomical starting points differ. In summary, our results support the view that extreme phenotypes emerge not by escaping constraint, but by evolving along it. When strong phenotypic integration aligns with ecological opportunity, evolution can generate exaggerated morphologies through coordinated, stepwise change. This framework helps reconcile the rarity of extreme traits with their repeated emergence across deep evolutionary time and provides a general model for understanding how constraint and innovation jointly shape the limits of organismal form.

## Methods

For additional details, including materials examined, analytical procedures, and bioinformatic workflows, see the SI Appendix, Extended Materials and Methods.

### Study system and dataset

We assembled ten complementary datasets, six focused on spikefishes (Tetraodontiformes: Triacanthodidae) and four used for comparative analyses with snipefishes. Spikefish datasets included Dataset 1, genome-wide data for phylogenetic reconstruction (49 specimens, 15 species plus four outgroups; Tables S1-S4); Dataset 2, a discrete morphological character matrix for total-evidence divergence-time estimation (24 extant and two fossil taxa, 80 characters; Tables S2, S5, and S6); Dataset 3, three-dimensional geometric morphometric data capturing macroevolutionary cranial shape variation from CT scans (23 specimens, 23 species; Tables S29 and S30), using a modified landmark scheme optimized for spikefish snout morphology (247 landmarks: 65 fixed, 182 semi-sliding; Figure S29); Dataset 4, linear cranial measurements from X-ray images used to quantify within-species phenotypic variance (six traits, 228 specimens, 24 species; Tables S31 and S32), with sampling emphasizing long-snouted (*Halimochirurgus alcocki*, *H. centriscoides*) and short-snouted (*Parahollardia lineata*, *P. schmidti*) taxa; Dataset 5, dental morphology data used as functional proxies for feeding strategy (40 specimens, 23 species; Tables S21 and S29); and Dataset 6, X-ray data used to identify species exhibiting cranial elevation and to distinguish high- and low-head-mobility clades (144 specimens, 24 species; Table S33). Datasets incorporating snipefishes (Syngnathiformes: Centriscidae) included: Dataset 7, CT-based cranial data used for qualitative and landmark-based comparisons with long-snouted snipefishes (24 spikefish species and one snipefish; Table S34), and Dataset 8, qualitative functional manipulation data from freshly collected spikefish specimens used to assess cranial and hyoid mobility. Sampling emphasized taxonomic completeness and integrated museum specimens examined using multiple imaging approaches alongside fresh material when available. Dataset 9 (Table S26) included linear morphometric-based body-shape data spanning Percomorpha for broad-scale snipe- and spikefish comparisons (278 species with pre-existing data and 46 spikefish individuals representing 22 spikefish species). Dataset 10 (Table S28) included 23 cleared and stained specimens for recreating head-elevation mechanics across a diversity of percomorph fishes, with an emphasis on tetraodontiform and triacanthoid species.

### Genomic data generation, phylogenetic inference, and divergence-time estimation

We generated genomic data for 49 individuals representing 15 spikefish species and four outgroups (Dataset 1) using a combination of targeted exon capture and whole-genome sequencing (Tables S1, S3, and S4). Exon capture libraries were enriched using the FishLife Eupercaria probe set^84^ and sequenced on Illumina platforms, and whole-genome libraries were sequenced to approximately 20-40x coverage. Sequence cleaning, assembly, exon mining, alignment, and quality control followed established FishLife pipelines^84^, including exclusion of poorly aligned loci and taxa with insufficient gene recovery (Tables S3 and S4). After filtering, the final dataset comprised 1,108 exon alignments across 17 species, which were concatenated for downstream phylogenetic analyses.

We inferred phylogenetic relationships using morphology-only (Dataset 2), molecular-only (Dataset 1), and total-evidence datasets (combined Datasets 1 and 2) under parsimony, maximum likelihood, Bayesian inference, and multispecies coalescent frameworks. Morphology-only analyses were based on a newly generated matrix of 80 discrete characters scored for all 24 extant spikefish species, both described fossil taxa, and two outgroups (Table S6). Maximum parsimony trees were inferred using TNT^95^. Molecular phylogenies were estimated in IQ-TREE II^96^ using both gene-specific and concatenation-based approaches, and resulting gene trees were used as input for coalescent-based species-tree inference in ASTRAL-III to account for incomplete lineage sorting^97^. One taxon (*Hollardia meadi*) exhibited an exceptionally long molecular branch and was excluded from downstream Bayesian divergence-time analyses.

We estimated divergence times using Bayesian tip dating under the Fossilized Birth-Death (FBD) model^98^ implemented in MrBayes^99^. To accommodate heterogeneity among loci, the exon dataset was partitioned into four non-overlapping subsets of 277 genes each, chosen to contain comparable numbers of parsimony-informative sites^100^. Two fossil spikefish taxa (†*Prohollardia avita* and †*Carpathospinosus propheticus*) were included as terminal tips with stratigraphic age priors, and alternative root-age priors were applied based on previous tetraodontiform phylogenomic studies^55,56^ We evaluated alternative molecular clock models under different taxon-sampling schemes using stepping-stone^101^ marginal likelihood estimation and selected best-fitting clock models accordingly. Molecular and morphological partitions were modeled under standard substitution and character-evolution models, and speciation, extinction, and fossilization parameters were informed by estimates from previous phylogenetic analyses^102^. Topological constraints enforcing subfamilial monophyly were applied based on prior evidence^21,48,58^ and revised morphological data presented here for *Hollardia goslinei*.

### Phenotypic integration, lines of least resistance, and evolutionary modularity

We estimated lines of least resistance as the first principal axis (PC1; **p_max_**) of phenotypic variance-covariance matrix (**P**), which can approximate the major axis of genetic variance from the genetic variance covariance matrix (**G**) when controlled breeding designs are infeasible, particularly in non-model taxa such as deep-sea fishes. Empirical and theoretical work supports the use of **P** as a proxy for **G**^36,40,42^, and here we treat lines of least resistance as an empirically observable bias in phenotypic variation rather than a deterministic constraint or causal mechanism. Within-species **P** matrices were estimated from the residuals of size-corrected cranial measurements in Dataset 4, comprising six traits measured from X-ray images (head depth, snout length, snout depth, eye size, lower jaw length, and the ascending process of the premaxilla; Table S32) using the EvolQG package (https://github.com/lem-usp/evolqg/wiki). Snout curvature was accounted for by tracing the vomer–parasphenoid bone, and modified head length was used as a size proxy for allometric correction (Figure S30).

For each focal species, we estimated phenotypic variance–covariance (**P**) matrices from size-corrected trait values derived from Dataset 4 and evaluated robustness under two size-correction schemes, including and excluding snout length during correction (see Extended Methods for justification). To assess conservation of phenotypic integration within genera (using 3–4 PC axes or 80% of variance), **P** matrices were compared using Random Skewers, Krzanowski subspace projections, and correlations among PC1 vectors. Because results were consistent across size-correction schemes (+SnL and -SnL), matrices were pooled to the genus level for subsequent analyses. We estimated phenotypic divergence matrices (**D**) under both Brownian Motion and Ornstein–Uhlenbeck models following Simon et al.^39^ and retained the simpler Brownian model given near-identical matrix similarity metrics. Alignment between **p_max_** and **d_max_** was evaluated using vector correlations and assessed against a phylogenetic null following Simon et al.^39^: we simulated 10,000 six-trait datasets under multivariate Brownian motion on our phylogeny using the empirically estimated evolutionary rate matrix, recomputed **d_max_** for each simulated dataset, and tested whether observed **p_max_**–**d_max_** angles fell within the smallest 5% of simulated angles (Figure S11). Uncertainty in overall matrix subspace similarity was quantified via bootstrap resampling (Figure S11). Overall phenotypic integration within each focal taxon was summarized using eigenvalue dispersion.

We evaluated evolutionary modularity and integration across the skull using the three-dimensional cranial shape dataset (Dataset 3) by testing alternative modularity hypotheses and fine-scale bone-level partitions. Modularity was quantified using covariance ratios and associated effect sizes implemented in the geomorph package^103^. All modularity tests were performed under both local and global Procrustes superimposition to mitigate Pinocchio effects^67^ associated with extreme snout elongation. Integration strength within and between cranial structures was assessed using eigenvalue dispersion and two-block partial least squares analyses, respectively, with statistical comparisons among structures based on standardized effect sizes and permutation-based *p*-values. To test whether snout bones follow predictable evolutionary trajectories, we visualized two-block partial least squares relationships for all pairwise combinations of snout elements and evaluated species ordering along the resulting trendlines. Spatial patterns of cranial integration were further visualized using network graphs, with significance assessed after false discovery rate correction^104^. Full results are provided in the SI Appendix (Figure S13).

### Morphological diversity and evolutionary dynamics

We quantified dental morphology as feeding-related proxies using the dental morphology dataset (Dataset 5), based on CT scans of 40 individuals representing 23 spikefish species (including short- and long-snouted taxa), averaging measurements to species means when multiple individuals were available. Using a modified scheme from prior work^69,105^, we measured 22 functional traits capturing tooth morphology, spacing, counts, and jaw mechanics for both upper and lower jaws, with metrics generally based on the five tallest teeth per jaw (or four when only four teeth were present). In *Macrorhamphosodes* (n = 6), upper jaws lack teeth, so tooth-count and toothed-region variables for the upper jaw were adjusted by small constants to accommodate log transformations. Allometric corrections were applied using head length as a size proxy under two alternative size-correction schemes (+SnL and -SnL), and relationships between tooth traits and snout elongation were evaluated under both schemes (Figures S18 and S31).

To summarize morphological diversity and evolutionary dynamics, we integrated results from Datasets 3, 5, and 6 by generating phylogenetically informed morphospaces for cranial shape and dentition and contrasting low- and high-head-mobility clades. Phylogenetic trees were pruned to match each dataset, and morphospaces were constructed using principal component analyses of cranial shape (Dataset 3) and dental traits (Dataset 5); leading axes were used to visualize phenotypic distributions and generate traitgrams (Figure 4C). Using the cranial shape dataset (Dataset 3), we quantified evolutionary rates and morphological disparity at two phenotypic scales (per bone and per module) in geomorph, including explicit contrasts between high- and low-head-mobility spikefishes. We further tested relationships between phenotypic integration and both evolutionary rate and disparity using FDR-corrected regressions. Finally, we evaluated evolutionary modes of cranial module diversification by fitting gradual, constrained, and punctuated-process models in PulsR^82^, following Brennan et al.^106^. Because PulsR does not accommodate multiple PC axes simultaneously, we used PC1 and PC2 scores for each cranial module as proxies for multivariate shape, alongside linear cranial traits from Dataset 4. To assess the robustness of PulsR given our small phylogenetic tree, we evaluated the proportion of times PulsR could correctly identify a jump model using simulations. For each strongly supported occurrence of a jump model from our empirical tests (n = 30), we extracted the parameters estimated from the underlying model and simulated 34 datasets, totaling 1,020 tests (Table S25).

### Direction and ordering of snout evolution

Direction and ordering of snout evolution. To assess whether snout elongation evolved once or convergently within spikefishes, and whether associated anatomical changes accumulated in an ordered manner, we combined ancestral state reconstruction with tests of ordered character evolution using the discrete morphological dataset (Dataset 2). We first reconstructed the evolutionary history of snout elongation using ancestral state estimation implemented in *fitMk* in phytools^107^, scoring species as elongated based on characters 28 and 33 of the discrete matrix, which capture pterygoid closure (see also Figures 2B and 2C). We compared three alternative transition models: equal rates of gain and loss (symmetric), unequal rates (asymmetric), and an irreversible model allowing gains but not losses of snout elongation. Model fits were compared using likelihood-based criteria, and ancestral states were estimated by model-averaging results across equal-rates, all-rates-different, and irreversible models. All analyses were repeated across four alternative phylogenetic trees to account for topological uncertainty.

To address concerns about whether hemiplasy—which may appear as convergence if snout elongation was ancestrally polymorphic and retained in only some lineages—could be obscuring our snout-origin inference, we used two approaches: (a) a gene tree screen and a (b) simulation-based approach with parameter estimates drawn from our coalescent-based species-tree. First, we examined the proportion of gene trees that support a single snout origin. Specifically, we screened the 1,108 generated gene trees used for ASTRAL. We restricted the scan to gene trees containing at least one representative of each of the four focal groups necessary to show two independent snout-elongation events: c1 = *Bathyphylax*; c2 = *Macrorhamphosodes* or *Halimochirurgus*; c3 = *Paratriacanthodes*; and c4 = *Tydemania navigatoris*. We then examined the proportion of gene trees that grouped c1 + c2—that is, *Bathyphylax* as sister to any extremely elongated-snouted taxon, which would support a single origin of snout elongation—to the exclusion of both c3 and c4 (Figure S2).

Next, we simulated the likelihood of hemiplasy using Heist^59^. HeiST takes the ASTRAL species tree, including its branch lengths in coalescent units, and uses the user-defined distribution of a trait across taxa as input. It then randomly simulates many possible gene trees under that species-tree model and simulates trait-associated mutations along the branches of those simulated gene trees based on user-defined mutation rates. Only a fraction of these simulated histories reproduce the pre-coded trait pattern observed in the sampled taxa. HeiST retains those matching histories and asks whether the trait pattern is better explained by homoplasy (defined by two or more independent mutations) or hemiplasy (define by a single mutation inherited along a gene tree that differs from the species tree).

Accordingly, we subsampled our ASTRAL topology to include only the focal clade including the snout-elongated taxa in question and performed the analyses across three mutation rates spanning three orders of magnitude (0.05, 0.005, 0.0005)—a protocol mirrored in cichlid fishes in ref^108^. Because lower mutation rate parameters (i.e., HeiST s-values) make our observed data more unlikely to be simulated, we employed the following simulation scheme: s0.05 = 1,000,000 simulations, s0.005 = 4,000,000 simulations, and s0.0005 = 40,000,000 simulations— which recovered 491, 232, and 236 focal scenarios for comparing support for either homoplasy (two origins) or hemiplasy (single origin).

We further assessed the robustness of our *a priori* snout-coding regimes by fitting various OU models with either one, two, or three adaptive peaks using continuous snout-length data in the package OUwie^108^. Relative snout length for these continuous analyses was measured from CT scans as the linear distance from the anterior tip of the premaxilla to the anteriormost point of the orbit, standardized as a proportion of head length. Models ranged from OU1 (a single adaptive peak) to OU3, which described intermediate snout states as a second transition (see Extended Methods). Further, we performed an exploratory search for adaptive peaks using a Bayesian search using the package bayou. We supplied broad half-Cauchy priors for alpha and sigma² values, which covered a 95% confidence interval with upper and lower bounds defined by our OUwie parameter estimations. Bayesian analyses were performed in the package bayou^60^ under four shift-count priors (L = 1–4) to account for bias in prior choice, each run as two independent MCMC chains of 500,000 generations (eight runs total).

To test whether complex snout morphologies arose through additive, ordered anatomical changes rather than single-step transitions, we implemented HyperMk, a hypercubic-transition framework that models multicharacter evolution while restricting transitions to single-character changes^109^. Under this framework, the ancestral state (000) represents the absence of snout elongation, and gains of individual traits represent discrete anatomical rearrangements. We evaluated support for an ordered evolutionary sequence involving (i) articulation, compression, and elongation of the metapterygoid-mesopterygoid and palatine-ectopterygoid (pterygoid closure; characters 28 and 33), followed by (ii) elongation of the vomer-parasphenoid complex (character 11), and finally (iii) anterior rearrangement of the ectopterygoid and palatine (characters 33 and 34). Analyses were conducted under both reversible and irreversible transition matrices and repeated across all four phylogenetic trees, with model support assessed using Akaike weights (AICw). Given the small sample sizes of our datasets, we tested the robustness of HyperMk using two complementary simulation analyses: false-positive rate and statistical power. First, we fitted two alternative HyperMk models to the empirical data across the four trees to estimate transition-rate parameters for the subsequent simulations. These models were: (a) an unconstrained independent-accumulation model, in which off-path transitions were permitted, and (b) a constrained ordered-accumulation model (see Extended Methods). We then used the estimated parameters to construct transition matrices, which served as inputs for data simulation using the *simulate_mk_model* in castor^110^. For each tree, we generated 250 replicate datasets under each generating model.

### Spikefish-snipefish convergence

To evaluate morphological and functional convergence between spikefishes and snipefishes, we integrated cranial shape (Dataset 3), dental morphology and feeding proxies (Dataset 5), and X-ray–based head-mobility data (Dataset 6) with an expanded comparative cranial dataset incorporating published and museum CT scans of long-snouted snipefishes (Dataset 7), yielding a total of 25 species (Table S34). This combined dataset captures a broad gradient of snout elongation and cranial mobility across spikefishes and snipefishes. To focus comparisons explicitly on snout elongation, we defined long-snouted taxa as those in which a tubular snout is formed primarily by elongation of the ethmoid region and adjacent neurocranial elements (vomer, parasphenoid, ethmoid, and quadrate), with only minor contributions from jaw elements. In spikefishes, this condition occurs in *Halimochirurgus*, *Macrorhamphosodes*, and *Bathyphylax*. We further defined extreme snout elongation as snout length exceeding 30% of standard length, a threshold that includes snipefishes and the most elongate spikefishes but excludes *Bathyphylax*, which represents an intermediate condition. Functional convergence in cranial elevation and feeding mechanics was assessed using hyoid configuration as a proxy for head mobility, with mobility coded based on the presence of both horizontal and vertical hyoid orientations among conspecific individuals (Figure S26; Table S33; Dataset 6). Anatomical substrates of convergence were evaluated by identifying candidate functional analogs between spikefishes and snipefishes and quantifying their similarity by applying the spikefish cranial landmarking scheme (Dataset 3) to snipefish specimens. Joint local Procrustes superimpositions and PCA-based comparisons were then used to assess bone-shape similarity across clades (Table S34). Tooth traits were additionally used to compare jaw opening and closing mechanical advantages as functional correlations in feeding convergence (Figure S25).

Finally, we expanded our convergence analyses to include a broader phylogenetic diversity spanning 300 percomorph fishes (Table S26). Phylogenetic hypotheses for these analyses stem from the tree estimated in this study and the grafting of four phylogenies (this study and refs. ^49,84,102^). We complemented publicly available FishShapes^86^ data with an additional 46 spikefish specimens representing 22 species. In brief, FishShapes spans eight measurements of body shape that have been shown to be major axes of variation across fishes—however, snout length is not a measurement included in these datasets. Using the Ct metrics from the convevol package^87,88^., we estimated statistical significance of spike- and snipefish convergence across three conceptually different workflows, which varied in size-correction strategy and the use of ordinary or phylogenetically informed morphospace coordinates, which were used as input to Ct tests (see Extended Methods). Focal taxa for the convergence test comprised the four extremely long-snouted spikefishes (*Halimochirurgus alcocki*, *H. centriscoides*, *Macrorhamphosodes uradoi*, and *M. platycheilus*) and the *Macroramphosus* snipefishes; all other percomorphs were treated as background taxa.

### Functional validation

To qualitatively assess the mechanical plausibility of feeding-related cranial mobility inferred from comparative analyses, we conducted functional manipulations of fresh spikefish specimens (Dataset 8) collected during the 2025 Coral Sea Frontier expedition aboard the RV Investigator (CSIRO, Australia). Six specimens representing five species spanning a gradient of snout lengths were examined: *Triacanthodes ethiops* and *Tydemania navigatoris* (short snout), *Bathyphylax bombifrons* (short to intermediate snout), and *Macrorhamphosodes uradoi* and *Halimochirurgus centriscoides* (extremely long snouts). Specimens were captured using Scampi or beam trawls at depths of 350–750 m and, although deceased upon retrieval, were placed in a seawater-filled container lined with a mesh lifting net and dosed with fish anesthetic (Aqui-S) to minimize tissue stiffness during manipulation. For each specimen, we manually manipulated the head and hyoid apparatus to assess coupling between head elevation and hyoid depression, recording the approximate degree of head elevation achieved. Previous work has shown that fresh snipefish material exhibits mechanically reversible coupling between hyoid depression and head elevation (Longo et al.^27^) and abrupt transitions between horizontal and vertical hyoid configurations during cranial elevation (Wainwright et al.^89^). Accordingly, we evaluated whether similar patterns could be recreated in spikefishes, including whether head elevation differed when the hyoid was depressed prior to cranial elevation versus when the suspensoria were adducted. When feasible, manipulations were conducted independently by two investigators following a standardized protocol, with short-snouted species serving as functional controls. Manipulations were video-recorded when possible; however, not all tests could be performed on every specimen due to variable specimen condition. These observations are therefore interpreted as qualitative assessments of mechanical potential rather than direct measurements of *in vivo* feeding kinematics.

To further examine the gain of head elevation in spikefishes within a broader phylogenetic context, we recreated head-elevation mechanics using cleared and stained specimens (Dataset 10) to calculate head and hyoid rotational capacities and subsequent kinematic transmission (KT) values. In brief, KT values demonstrate mechanical sensitivity by evaluating the ratio of angle changes in the input and output links (the head and hyoid, respectively) and can be compared across diverse taxa (see Extended methods). Our sampling included eleven non-tetraodontiform suction feeding fishes (including three Syngnathiformes), seven non-spikefish tetraodontiforms spanning seven families, and five spikefishes spanning both subfamilies and snout phenotypes.

## Data availability

All materials are available in the main text and the supplementary data on Zenodo Digital Repository at: https://doi.org/10.5281/zenodo.22033939. Raw data and sequence alignments are available on NCBI under BioProject number (SUB15379368). Any additional information required to reanalyze the data reported in this paper is available from the lead contact (Dahiana Arcila) upon request.

## Code availability

All code and scripts are available on Zenodo Digital Repository: https://doi.org/10.5281/zenodo.22033939

## Acknowledgements

We thank Matt Girard whose guidance on the morphological matrix proved invaluable. We also thank Liam Ward, Bill Ludt, Todd Clardy, Emily Troyer, Kory Evans, Leo MacLeod and Matt Girard for generating additional CT scans. We thank Shing-Lai Ng for helping share additional spikefish samples for sequencing. We are grateful to Daniel Moen and Monique Simon for insightful discussions on phenotypic and divergence matrices. We are indebted to all museum and collections staff who helped provide materials for this project and to Paweł Socha and Krzysztof Stefaniak for providing photographs of fossil specimens. Additionally, to those who helped directly or indirectly in X-ray generation including Jessica Zhong, Ben Frable, Amanda Hay, Mark Sabaj, Sarah Huber, Kevin Conway, Vinicius Correa Espindola Glen Moore, Jenelle Ritchie, Worth Pugh, Abigail Reft, Diane Pitassy, Susan Mochel, Caleb McMahan, and Jon Fong. Additionally, we are thankful to Griffin McRory, Matthew Burnett, Jay Champlin, and Zishuoguo Li for helping generate photos for cleared and stained specimen manipulation. We are grateful to the University of Oklahoma Supercomputing Center for Education & Research (OSCER) for providing bioinformatics support and guidance. We thank Lynne Parenti and the Smithsonian Institution Division of Fishes for facilitating ZH’s visit through the Schultz Fund. We additionally thank Floriaan Devloo-Delva and Younis Menkara for assistance with video recording during functional manipulations, as well as all members of the 2025 Coral Sea Frontier expedition for their help during specimen handling.

## Additional information

Correspondence and requests of materials should be addressed to D.A.

## Ethics declarations

## Contributions

This study was conceived and designed by Z.H. and D.A. The morphological matrix was curated by Z.H., K.E.B., and J.C.T. Z.H. and F.M.V. conducted DNA extractions. Phylogenomic and total evidence trees, as well as the divergence-time estimation scheme were conducted by Z.H. and supervised by A.S., R.B.R., and D.A. CT-scan data were collected by Z.H., K.B., K.F., N.M and museum colleagues. X-ray generation was conducted by Z.H., J.J.P., H.-C.H., N.M. K.E.B, and museum colleagues. Body-shape data for the convergence analyses, were collected by Z.H. Functional manipulations of fresh specimens were conducted by D.A., J.P., and W.W. Cleared- and-stained head-elevation experiments and four-bar linkage characterization were conducted by Z.H. with guidance from M.W., who also advised on the biomechanical analyses and their interpretation. Fish illustrations and figures were prepared by Z.H., C.L., A.S., and D.A. Tissue samples and additional specimens were collected by J.P., W.W., H.-C.H., N.M, and by museum colleagues. Phylogenetic comparative analyses were conducted by Z.H. Data analyses were conducted by Z.H. The manuscript was written by Z.H. and D.A., with input from all authors.

