## Supplementary materials for "Convergent evolution of extreme snout elongation in deep-sea fishes reveals constrained trajectories"

**This file includes a detailed explanation of all the supplemental information:**

- Extended Methods
- Figures S1 to S31
- Tables S1 to S34 deposited in Zenodo:
    - Table S1. Specimens and metadata for phylogenomic analysis.
    - Table S2. Species with only morphological data in Bayesian tree estimation.
    - Table S3: Alignments quality control.
    - Table S4: Alignments used in downstream analyses.
    - Table S5: Materials examined, morphological matrix.
    - Table S6: Scored morphological matrix.
    - Table S7: Divergence times per scheme per subset.
    - Table S8: The results from clock fitting using the Stepping-stone approach implemented in MrBayes.
    - Table S9: Morphological characters consistent with independent snout elongation in *Bathyphylax*.
    - Table S10: The results of two-block partial least squares tests for all bones in the snout module.
    - Table S11: *Macrorhamphosodes platycheilus* intermediate status.
    - Table S12: Within and between genera **P** matrix comparisons.
    - Table S13: Alignment of species- and genus-level PC1 with divergence PC1.
    - Table S14: Modularity fitting results.
    - Table S15: The results of integration compare.ZVrel for bones.
    - Table S16: The results of integration compare.ZVrel for modules.
    - Table S17: Integration network values used to create network graph for bones.
    - Table S18: Integration network values used to create network graph for modules.
    - Table S19: Dataset for testing relationship of integration, rates of evolution and morphological disparity for each bone.

- Table S20: Dataset for testing relationship of integration, rates of evolution and morphological disparity for each module.
- Table S21: Description of 22 linear measurements for tooth dataset.
- Table S22: Tooth morphometric dataset PC1&2 loadings, size corrected by head length.
- Table S23: Tooth morphometric dataset PC1&2 loadings, size corrected by snout-subtracted head length.
- Table S24: Summary diet table.
- Table S25: PulsR simulations.
- Table S26: Raw FishShapes data for Ct analyses.
- Table S27: Ct results across size-correction methods.
- Table S28: Head mobility estimates from cleared and stained specimens.
- Table S29: Materials examined, CT scans.
- Table S30: Landmark metadata, spikefish only.
- Table S31: Materials examined for estimation of **P** and **D** matrices.
- Table S32: Description of linear measurements for cranial dataset.
- Table S33: Materials examined, head rotation.
- Table S34: Landmark metadata, spikefish and snipefish.

### Extended Materials and Methods

**Study system and datasets.** We assembled ten complementary datasets, six to investigate the evolution of snout elongation in spikefishes (Tetraodontiformes: Triacanthodidae) and four additional datasets for assessing morphological convergence with snipefishes (Syngnathiformes: Centriscidae). Spikefish datasets included: Dataset 1, genome-wide data for phylogenetic reconstruction (Tables S1–S4); Dataset 2, a discrete morphological character matrix for total-evidence divergence time estimation (Tables S2, S5, and S6); Dataset 3, three-dimensional geometric morphometric data for skull shape analysis (Tables S29 and S30); Dataset 4, linear cranial measurements for within-species phenotypic variance (Tables S31 and S32); Dataset 5, dental morphology data used as functional proxies for feeding strategy (Tables S21 and S29); and Dataset 6, an X-ray dataset of spikefishes for identifying species with cranial elevation (Table S33). Snipefish convergence datasets included: DatGaset 7, CT-based cranial data used for qualitative and landmark-based comparisons with long-snouted snipefishes (24 spikefish species and one snipefish; Table S34); and Dataset 8, qualitative functional manipulation data from freshly collected spikefish specimens used to assess cranial and hyoid mobility. Sampling emphasized taxonomic completeness and integrated museum specimens examined using multiple imaging approaches alongside fresh material when available. Dataset 9 (Table S26) included linear morphometric-based body-shape data spanning Percomorpha for broad-scale snipe- and spikefish comparisons (278 species with pre-existing data and 46 spikefish individuals representing 22 spikefish species). Dataset 10 (Table S28) included cleared and stained material for recreating head-elevation mechanics across a diversity of percomorph fishes, with an emphasis on tetraodontiform and triacanthoid species.

**Genomic data generation and processing.** We extracted genomic DNA from 49 individuals, representing 15 spikefish species and four outgroups using the DNeasy Blood and Tissue Kit (Qiagen) following the manufacturer's protocol. We combined targeted exon capture and whole-genome sequencing to maximize locus recovery across specimens (Tables S3 and S4). Exon capture libraries were enriched using the FishLife Eupercaria probe set[1] and sequenced on Illumina NovaSeq 6000 platforms (150-bp paired-end reads) at Arbor Biosciences (Ann Arbor, MI). Whole-genome libraries were prepared and sequenced on an Illumina NovaSeq 6000 platform (150-bp paired-end reads), targeting 20–40X coverage per sample, at the Oklahoma Medical Research Foundation (OMRF).

Sequence cleaning, assembly, exon mining, and alignment followed established FishLife pipelines[1]. First, low-quality reads and adapters were removed using Trimmomatic v.039[2]. Subsequent trimmed reads were mapped to exonic reference sequences with BWA v.0.7.17[3] followed by removal of PCR duplicates in SAMtools v.1.9[4]. Processed reads were subjected to two rounds of assembly, initially using Velvet v.1.2.10, and the longest contig was identified using the getLongest.py script[1,5]. The second assembly extended the longest contig for each exon by using aTRAM v.2.2 with the Trinity v.2.2 assembler[6,7]. If contigs were present multiple times, they were identified and excluded using CD-HIT-EST v.4.8.1[8]. Reading frames were identified using Exonerate v.2.4.0[9]. Exons were aligned with MACSE v.2.05[10], cleaned using the AlignmentCleanerCodons.py script, and visually inspected in Geneious Prime v.2050.0.3 (https://www.geneious.com). Species with poor gene alignments (affecting seven genes) or fewer than 100 genes post-alignment were excluded. Furthermore, genes with fewer than ten taxa were also excluded (Tables S3 and S4). Post quality control, 1,108 gene alignments remained across 17 species (Dataset 1). Gene alignments were concatenated with the AMAS.py[11] Python script for downstream phylogenetic analyses. Full commands, scripts, and files are provided in the Supplementary Information.

**Discrete morphological characters.** We scored 80 discrete morphological characters across 239 specimens (Dataset 2), including ethanol-preserved (n = 90), cleared and stained (n = 85), and CT-scanned individuals (n = 64), with resolutions ranging from 16 to 54 μm, capturing all extant spikefishes, two fossil species (†*Prohollardia avita* and †*Carpathospinosus propheticus*), and two outgroups from the established sister family. Characters were designed for morphology-only and total-evidence phylogenetic inference and for explicit tests of ordered anatomical transitions associated with snout elongation. Additional details including descriptions of each morphological character are provided below.

**Phylogenetic inference.** Phylogenetic relationships were inferred using morphology-only (Dataset 2, see below), molecules-only (Dataset 1), and total-evidence (combined Dataset 1 and 2) under Parsimony, maximum likelihood (ML), Bayesian inference, and multi-species coalescence (MSC) frameworks. This multi-method approach allowed us to assess topological robustness to data type and inference strategy. For the morphology-only dataset, we analyzed our newly generated matrix of 80 discrete characters scored for all extant and fossil spikefishes and outgroups (Dataset 2) using Parsimony, ML, and Bayesian frameworks (Table S6). Parsimony analyses were conducted in TNT v1.6[12] using multistate characters (nstates 32) with gaps treated as missing data (nstates NOGAPS). Tree searches retained up to 1,000 trees (hold 1000) and employed 1,000 ratchet iterations (mult=ratchet 1000) to obtain the most parsimonious trees. When multiple equally parsimonious trees were retrieved, a strict consensus tree was estimated (nelsen*). Node support was assessed using 1,000 bootstrap replicates. Maximum likelihood analyses were performed in IQTREE v1.6.12[13] using the Mk model with correction for ascertainment bias (-m mKv). Polymorphic character states are not supported in IQTREE-II and therefore were recoded as missing data ("?") in 20 cells (~0.9% of the matrix). Node support was assessed with 1,000 ultrafast bootstrap replicates[14]. Bayesian inference was conducted in MrBayes v.3.6.7[15] under the Mk model, defining rates as drawn from a gamma distribution and coding set to "variable" to account for ascertainment bias. The Markov chain Monte Carlo analyses were run for 20 million generations using four chains and a temperature of 0.1, with sampling and print frequencies of 1,000, across eight independent runs. Convergence was assessed in Tracer v1.7.2[16] with effective sample sizes exceeding 200 for all parameters.

Molecular phylogenies were inferred from the concatenated 1,108-exon dataset using both concatenation and coalescent-based approaches. Concatenation-based analyses used parsimony and ML. Parsimony inference followed the parameters outlined for the morphology-only dataset, adjusted for nucleotide data (nstates DNA). ML analyses were performed in IQTREE under a codon-based scheme (which internally selected the GTR model for each partition) and a mixture model approach (-m MIX{JC, K2P, HKY, GTR, TPM2})[17]. Node support was assessed with 1,000 ultrafast bootstrap replicates. Subsequently, gene trees were estimated in IQTREE using the same partitioning and bootstrapping workflow. To account for potential gene-tree discordance arising from incomplete lineage sorting, we additionally inferred a species tree using ASTRAL-III[18]. Support values were derived from quartet frequencies (-t 10)[18]. One taxon (*Hollardia meadi*) exhibited an exceptionally long molecular branch and was excluded from downstream Bayesian dating analyses to avoid distortion of branch-length estimation. We integrated molecular and morphological data under parsimony, ML, and Bayesian frameworks to infer total-evidence phylogenies. Bayesian total-evidence analyses were conducted in MrBayes under the Fossilized Birth-Death (FBD) process[19,20], allowing fossil taxa to be treated as terminal tips rather than node calibrations. This approach jointly estimates topology, divergence times, and rates of lineage diversification while explicitly modeling fossil sampling; specific settings are outlined below.

**Divergence-time estimation.** To estimate divergence times while accounting for heterogeneity in phylogenetic signal across loci, we partitioned the 1,108-exon dataset into four non-overlapping subsets with comparable numbers of parsimony-informative sites (PIS) using a custom R script[21]. This strategy reduces stochastic variation among subsets and allows independent estimation of divergence times under replicated data partitions. Next, divergence times were estimated using Bayesian tip dating under the Fossilized Birth-Death (FBD) model implemented in MrBayes. Two fossil spikefish taxa (†*Prohollardia avita* and †*Carpathospinosus propheticus*) were included as terminal tips with stratigraphic age priors reflecting their known temporal ranges. Root-age priors followed previous phylogenomic studies of tetraodontiform fishes and were implemented under two alternative schemes: one including the sister family Triacanthidae (95% CI: 61.1–90.4 Ma)[22] and one excluding it (95% CI: 35.2–44.7 Ma)[23].

Bayesian inference effectively integrates observed and prior information but can be sensitive to the priors supplied, such as tree age and clock models[24]. Because deep divergences between Triacanthodidae and Triacanthidae introduce substantial rate heterogeneity, we explicitly evaluated clock-model performance under both taxon-sampling schemes. Marginal likelihoods were estimated using stepping-stone sampling[25] to compare an uncorrelated independent gamma rates (IGR) clock with an autocorrelated Thorne-Kishino (TK02) clock. Model comparison revealed that the preferred clock model depended on outgroup inclusion: the IGR clock was favored when Triacanthidae were included, whereas the TK02 clock fitted better when analyses were restricted to Triacanthodidae. This sensitivity is consistent with known challenges of clock-model selection in datasets spanning large rate shifts and long subtending branches.

Unless otherwise stated, the same priors and evolutionary models were applied across all dating analyses. Molecular data were modeled under a GTR+I+G substitution model with codon-based partitions, allowing substitution rates, invariable sites, base frequencies, and gamma distributions to vary independently across partitions. Morphological characters were treated as a single partition under the Mk model with ascertainment bias correction. Speciation priors were calculated using the *fit.bd* function in the R package phytools[26], based on a previously published time-calibrated phylogeny for tetraodontiform fishes[22] pruned to include only Triacanthidae and Triacanthodidae. Extinction and fossilization were assigned a beta distribution (0–1), and branch lengths were modeled using the 'fossilization' prior appropriate for the FBD framework. Because all described extant species were included, the sampling fraction was set to 1. Based on previous analyses supporting the monophyly of spikefish subfamilies[22,23,27–30], we imposed two topological constraints corresponding to subfamilial designations[29]. An exception was made for *Hollardia goslinei*, which was placed within Triacanthodinae based on revised morphological evidence. All constraint schemes and associated input files are provided in Appendix 1.

### Continuous morphological trait datasets

**Three-dimensional skull shape.** Cranial shape was quantified using three-dimensional geometric morphometrics from CT scans (Dataset 3). One representative specimen per species was selected based on scan quality to capture macroevolutionary differences, excluding *Bathyphylax omen* due to poor scan quality. Within-species variation was addressed separately using linear measurements (see below). We adapted the landmarking scheme of Evans et al.[31], originally comprising 178 landmarks, by removing 21 landmarks inapplicable to spikefishes and adding 90 new landmarks to densely sample the snout region (Figure S29; Table S30). The final dataset comprised 247 landmarks, including 65 fixed (51 from Evans et al. and 14 newly defined) and 182 semi-sliding landmarks (106 from Evans et al. and 76 newly defined).

Landmark coordinates were aligned using Procrustes superimposition with the *gpagen* function in the R package geomorph[32], which rescales, translates, and rotates homologous landmarks to minimize inter-landmark distances. To address artifacts associated with extreme snout elongation (Pinocchio effect)[33], whereby disproportionately displaced landmarks can distort global alignments (Figure S20), we compared two superimposition strategies: i) a global Procrustes superimposition (GPS), in which landmarks are aligned simultaneously, and ii) a local, bone-by-bone, Procrustes superimposition (LPS) following Rhoda et al.[34]. For LPS, we used the mean spikefish shape (calculated using *mshape* in geomorph) as a reference configuration to avoid biases associated with selecting either long- or short-snouted taxa as templates. Because LPS better accounts for preservation artifacts[34], all downstream shape analyses were conducted using LPS coordinates unless otherwise stated.

**Linear cranial measurements and phenotypic covariance.** To quantify within-species trait covariation, we measured six linear traits from X-rays of 228 spikefish skulls, representing all 24 described species (Tables S31 and S32; Dataset 4). Sampling emphasized four 'focal taxa': the two long-snouted species of *Halimochirurgus* (*H. alcocki* and *H. centriscoides*), two short-snouted species of *Parahollardia* (*P. lineata* and *P. schmidti*), and one individual for each of the remaining 20 species. Because extreme snout elongation can inflate covariation when head length is used as a size proxy[35], we applied two size-correction schemes: head length including snout length (+SnL) and head length excluding snout length (-SnL). Analyses were conducted under both schemes, and inferences were considered robust only when consistent across corrections.

### Phenotypic integration and lines of least resistance

**Estimation of lines of least resistance and phenotypic integration.** Lines of least resistance were operationally defined as the first axis of phenotypic variance (PC1), which can approximate the major axis of genetic variance (**G**) when controlled breeding designs are infeasible, particularly in non-model taxa[36–38]. Because such designs are rarely possible for deep-sea fishes, multiple empirical and theoretical studies have shown that PC1 of **G** is often well approximated by PC1 of the phenotypic variance-covariance matrix (**P**)[36–38]. For justification and empirical validation of approximation, see Simon et al.[39]. Throughout, we treated lines of least resistance as an empirically observable bias in phenotypic variation rather than as a deterministic constraint or causal mechanism. To quantify within-species trait covariation, we collected six linear cranial traits using ImageJ: head depth, snout length, snout depth, eye size, lower jaw length, and the ascending process of the premaxilla (Table S32). Traits were chosen to minimize sensitivity to skull orientation. Thirty-one *Halimochirurgus* were not identified to the species level or possessed improper identification; subsequent identification was evaluated from X-ray images and preliminary morphospatial observations. To account for snout curvature caused by fixation artifacts or specimen damage, head length and snout length were measured along the curved vomer-parasphenoid axis using the line-segment tool. We verified that this approach provides a reliable proxy for linear snout length (Figure S30). These modified measurements prevented systematic underestimation of snout length and reduced bias in variance estimates. Modified head length was used both as a proxy for overall body size and for size correction.

For each focal species, we estimated **P** matrices using size-corrected trait values. Specifically, from each species we extracted residuals from linear models of each trait on head length and then calculated **P** matrices using the *CalculateMatrix* function in the EvolQG package (https://github.com/lem-usp/evolqg/wiki). Because allometric size can inflate estimates of trait covariation[35], and because extremely long snouts may similarly bias covariances, we repeated analyses with +SnL and -SnL.To assess whether trait integration was conserved within genera, we compared **P** matrices using the following three similarity indices: (1) Random Skewers (RS)[40], which calculates the average correlation of response vectors generated by multiplying each matrix by a set of random vectors (*RandomSkewers* function in EvolQG); (2) Krzanowski projections (KrzCor)[41], which estimates the similarity of matrices by comparing the eigenvector subspaces defined by a specified number of PC axes (*KrzCor* function in EvolQG); and (3) PC1 vector correlations, which are estimated using the absolute value of the dot product of two normalized PC axes. All matrix indices were scaled between 0 to 1, with 1 indicating perfect similarity. Because within-genus matrix indices showed overall similarity regardless of size correction method, we pooled matrices to the genus level. Pooled **P**-matrix calculation was repeated as above, with species as a fixed factor during residualization.

We estimated divergence matrices (**D**) following Simon et al.[39] under Brownian Motion (BM) and Ornstein-Uhlenbeck (OU) models of evolution. This approach uses phylogenetic independent contrasts to estimate evolutionary rates and trait coevolution. Similarity indices showed divergence matrices were highly similar (RS = 0.97; KrzCor = 0.99), so we used BM for downstream analyses as it was the simpler model. We estimated vector correlations for PC1 and KrzCor similarities for PC1–PC4, as this range frequently covered the majority of explained variance. We assessed significance of vector correlations against a phylogenetic null following Simon et al.[39]: we simulated 10,000 six-trait datasets under multivariate Brownian motion on our phylogeny using the empirically estimated evolutionary rate matrix, recomputed dmax for each simulated dataset, and tested whether observed pmax–dmax angles fell within the smallest 5% of simulated angles (Figure S11). We further quantified uncertainty in KrzCor similarities of **D** to the pooled **P** of either genus by using 1,000 bootstrapping replicates with resampling (Figure S11).

Although our size-correction procedures explicitly minimize allometric size as a contributor of lines of least resistance, we emphasized three considerations motivating this approach. First, the rarity of spikefishes and the absence of developmental series prevented reliable discrimination among ontogenetic stages. Second, **P** matrices are sensitive to small sample size; despite this, our within-species datasets included 37–82 specimens per species, a large proportion (30–70%) of all specimens reported globally per iDigBio (https://www.idigbio.org/). Third, spikefishes represent one of the most extreme cases of snout-elongation among ray-finned fishes, with snouts comprising up to 50% of total body length in long-snouted taxa, rendering conventional size proxies difficult to interpret. To further address these issues, we also calculated allometric slopes of snout length across focal species, given that snout length is the primary trait of interest. Lastly, we quantified overall phenotypic integration using eigenvalue dispersion (EDV), which measures how variance is distributed across the leading eigenvectors of a covariance matrix, calculated for each focal species using the *integration.Vrel* function in geomorph.

**Evolutionary modularity and integration.** To evaluate patterns of covariation across the spikefish crania, we tested 11 alternative modularity hypotheses using the *phylo.modularity* function in geomorph, each assessed with 1000 iterations (Figure S12). Under this framework, strong modularity is inferred when user-defined modules exhibit high internal integration and semi-autonomous evolutionary trajectories from other modules. Modularity strength was quantified using covariance ratios[42] (CR) and associated effect sizes (Z-scores) calculated in the *compare.CR* function of geomorph, with lower CR values indicating stronger modular signal. In addition to evaluating predefined modular hypotheses, we tested integration under a fine-grained framework, in which each cranial bone was treated as its own module. For clarity, we refer collectively to bones and multi-bone modules as structures. Unless otherwise specified, all modularity and integration tests were performed twice: once treating each bone as an independent structure and once using multi-bone modules defined by the best-supported modular hypothesis. To characterize how integration was distributed within and between structures, we quantified within-structure integration using eigenvalue dispersion calculated with the *integration.Vrel* function in geomorph. This metric measures how variance is distributed across the leading eigenvectors of a structure's variance-covariance matrix and is comparable across datasets[35,43]. Effect sizes were calculated following Conway et al.[36], and statistical differences among structures were evaluated using *compare.ZVrel*. For interpretability, Z-scores were converted to ZR ratios, with lower ZR values indicating lower integration.

To assess whether snout bones followed a predictable evolutionary trajectory, we conducted two-block partial least squares (2BPLS) plots (*two.b.pls* in geomorph) for all pairwise combinations of snout bones. Although conceptually similar to PCA space, 2BPLS maximizes the *covariation* between two partitions of shape, rather than the *variance* within a single dataset. A predictable pattern of evolution would be supported if *Bathyphylax*, the convergently evolved intermediate snout-length species, also had intermediate placement in 2BPLS space compared with short snout and fully snout-elongated species. These analyses were conducted twice using both GPS and LPS Procrustes coordinates to assess robustness to superimposition strategy. Broader patterns of skull integration were assessed using phylogenetically informed 2BPLS tests with *phylo.integration* in geomorph. This method estimates integration and statistical significance for all pairwise comparisons of bones, but it does not provide trait space visualization. Because this approach estimates a large number of tests (105 pairwise comparisons), we performed *p*-values for false discovery rates (FDR)[44] using the *p.adjust* function in the stats package.

To visualize the spatial distribution of integration across the cranium, we plotted network graphs using the *plot.Network* function in qgraph[45], with structures represented as nodes and significant pairwise integration strengths (Z-scores) represented as branches[46]. Following FDR correction, non-significant pairwise comparisons were excluded, and for visual purposes, only the strongest one-third comparisons were plotted in Figure 2. A complete network, including all pairwise comparisons, is provided in Figure S13.

### Morphological diversity and evolutionary dynamics

**Dental morphology and feeding proxies.** To assess whether snout elongation is associated with shifts in feeding-related morphology, we quantified dental traits in spikefishes using CT-based measurements (Dataset 5). The dental dataset comprised 40 individuals, representing 23 spikefish species, including both short- and long-snouted taxa. When multiple individuals were available, measurements were averaged to generate species-level means. CT scans were imported into 3DSlicer (v.5.2.1)[47] and SlicerMorph (v.9f375b5)[48] using the 'ImageStacks' module, and linear measurements were collected using the line segment tool in the Markups[48] module. Following a modified scheme from Heiple et al.[49] and Mihalitsis et al.[50], we quantified traits capturing tooth morphology, spacing, and jaw mechanics. Measurements focused on the five tallest teeth per jaw, when present, and were collected for both upper and lower jaws (U/L). Traits included average tooth length, width, and spacing; tooth aspect ratio; shape of the tooth base; horizontal gape; snout length; angular; distance from jaw joint to the tip of the first tooth; coronoid process length of the articular; in-lever and out-lever ratios; premaxillary protrusion; dentary length; tooth count (U/L); jaw length (U/L); extent of the toothed region (U/L); and the proportion of each jaw bearing teeth.

In long-snouted species with only four lower-jaw teeth, average spacing, width, and length were calculated using the four teeth rather than five. When multiple individuals were available, species means were calculated prior to comparative analyses. In all species of the genus *Macrorhamphosodes* (n = 6), the upper jaws lack teeth (values = 0), which complicates downstream log transformations. To accommodate these cases, tooth count and extent of the toothed region for the upper jaw were adjusted by adding small constants (+1.0 and +0.01, respectively). Prior to calculating species means, we performed allometric corrections using head length as a size proxy. Because snout elongation contributes disproportionately to head length in spikefishes, particularly in long-snouted taxa, we implemented two alternative size-correction schemes to evaluate robustness. In the first scheme (+SnL), snout length was retained as a variable and all traits were corrected using total head length. In the second scheme (-SnL), snout length was excluded as a variable and head length was corrected as *-SnL = head length - snout length*.

Allometric corrections were applied separately under each scheme, after which size-corrected values were transformed into species means. Although the protocol outlined by Mihalitsis et al.[50] recommends using residuals when allometric regressions are significant ($p < 0.05$), we did not adopt this step. Preliminary analyses indicated that extreme snout elongation strongly distorted regression slopes and residual distributions, producing biologically implausible patterns (Figure S31). Finally, we used linear regressions to examine how tooth length, spacing, width, and tooth counts covaried with snout elongation, conducting analyses under both +SnL and -SnL schemes where applicable (Figure S18).

**Quantification of skull and tooth diversity.** To visualize and quantify morphological diversity in cranial shape and dentition, we generated phylogenetically informed morphospaces for both datasets. Phylogenetic trees were pruned to match morphological datasets using the *drop.tip* function in ape[51]. Morphospaces were constructed using principal component analyses of dental traits (*prcomp*) and cranial shape data (*gm.prcomp*) in geomorph. Lastly, the first two principal components from each morphospace were used to visualize phenotypic distributions and to generate traitgrams using the phenograms function in phytools (Figure 4C).

**Rates of morphological evolution and punctuated evolution.** To characterize the tempo of cranial evolution in spikefishes, we quantified evolutionary rates and morphological disparity across two phenotypic scales, namely per bone, and per module. We calculated evolutionary rates for each structure using *compare.multi.evol.rates* in geomorph (iter = 1000), which provides a net rate of change for each shape partition. Initially, this was done across all species, and subsequently by comparing rates of high-head-mobility to low-head-mobility spikefishes.

To test whether phenotypic integration influenced evolutionary dynamics, we tested for relationships between integration and both evolutionary rate and morphological disparity using linear regressions implemented in the *lm* function in the stats package. We quantified integration using two complementary metrics: within-structure integration (eigenvalue dispersion estimated with *integration.Vrel*) and between-structure integration (mean pairwise Z-scores derived from *phylo.integration*). Morphological disparity was calculated at the structure level using Procrustes variance. All regression results were corrected for multiple testing using FDR procedures.

**Evolutionary model fitting.** In order to infer the evolutionary mode influencing cranial module diversification in spikefishes, we fitted a suite of evolutionary models using *fit_reml_levy* in the PulsR[52] package (*levy_pruning_optim.r*). We evaluated seven models, including gradual changes (Brownian Motion; BM), gradual evolution around an optimum (Ornstein Uhlenbeck, OU), an early burst of diversification (Early Burst, EB), explicitly punctuated processes (Variance Gamma, VG; Jump Normal, JN), and mixed models with both punctuated and gradual processes (Brownian Motion + Variance Gamma, BMVG; Brownian Motion + Jump Normal, BMJN). Because PulsR cannot accommodate multivariate trait blocks, we used PC1 and PC2 scores from principal component analyses of each cranial module as proxies for module-level variation. Although inclusion of all principal components would have been ideal, this approximation was necessary given current methodological constraints. Model fitting followed the protocol by Brennan et al.[53] with two modifications. First, we replaced the two 'Normal Inverse Gaussian' models that failed to converge for our dataset with two VG models. Second, we increased the number of optimization attempts (num_attempts = 10) to reduce sensitivity to local optima inherent to heuristic fitting procedures.

We further assessed the robustness of PulsR in correctly identifying pulsed models, given our small 24-tip phylogenies. In a post hoc analysis, we extracted the jump-model parameters from all instances in which they were strongly supported by our empirical data (n = 30). Using the representative trees and parameter estimates from each of these instances, we simulated 34 unique datasets with different seeds using the *levy_prurning_sim.r* script, totaling 1,020 simulations (1,017 analyzed after excluding three runs that failed to converge). Finally, we examined the proportion of runs in which PulsR recovered either any jump model or the specific jump model on which the data were simulated.

### Direction and ordering of snout evolution

**Quantifying snout-elongation events.** To assess whether snout elongation evolved once or convergently within spikefishes, we conducted ancestral state estimation with the *fitMk* function in phytools. Species were scored as elongated based on characters 28 and 33 in our discrete morphological matrix (Dataset 2), which capture the presence of pterygoid closure (also see Figures 2B and 2C). We tested three alternative transition models: (i) equal rates of snout elongation gain and loss (symmetrical); (ii) unequal rates (asymmetrical); and (iii) a custom irreversible model, which allows for snout elongation to be gained but not lost (irrev). Subsequently, fitMk objects were processed using the *anova* function and passed to the function *ancr* to estimate model averages. These analyses were conducted across four phylogenetic trees to account for topological uncertainty.

To address concerns about whether hemiplasy—which may appear as convergence if snout elongation was ancestrally polymorphic and retained in only some lineages—could be obscuring our snout-origin inference, we used two approaches: (a) a gene tree screen and a (b) simulation-based approach with parameter estimates drawn from our coalescent-based species-tree. First, we examined the proportion of gene trees that support a single snout origins. Specifically, we screened the 1,108 generated gene trees used for ASTRAL. We restricted the scan to gene trees containing at least one representative of each of the four focal groups necessary to show two independent snout-elongation events: c1 = *Bathyphylax*; c2 = *Macrorhamphosodes* or *Halimochirurgus*; c3 = *Paratriacanthodes*; and c4 = *Tydemania navigatoris*. We then examined the proportion of gene trees that grouped c1 + c2—that is, Bathyphylax as sister to any extremely elongated-snouted taxon, which would support a single origin of snout elongation—to the exclusion of both c3 and c4 (Figure S2).

Next, we simulated the likelihood of hemiplasy using HeiST[54]. HeiST takes the ASTRAL species tree, including its branch lengths in coalescent units, and uses the user-defined distribution of a trait across taxa as input. It then randomly simulates many possible gene trees under that species-tree model and simulates trait-associated mutations along the branches of those simulated gene trees based on user-defined mutation rates. Only a fraction of these simulated histories reproduce the pre-coded trait pattern observed in the sampled taxa. HeiST retains those matching histories and asks whether the trait pattern is better explained by homoplasy (defined by two or more independent mutations) or hemiplasy (defined by a single mutation inherited along a gene tree that differs from the species tree).

Accordingly, we subsampled our ASTRAL topology to include only the focal clade including the snout-elongated taxa in question and performed the analyses across three mutation rates spanning three orders of magnitude (0.05, 0.005, 0.0005)—a protocol mirrored in cichlid fishes in[55]. Because lower mutation rate parameters (i.e., HeiST s-values) make our observed data more unlikely to be simulated, we employed the following simulation scheme: $s_{0.05}$ = 1,000,000 simulations, $s_{0.005}$ = 4,000,000 simulations, and $s_{0.0005}$ = 40,000,000 simulations—which recovered 491, 232, and 236 focal scenarios for comparing support for either homoplasy (two origins) or hemiplasy (single origin).

We further tested whether our interpretation of snout length from discrete characters could be accurately recovered under a continuous snout-length framework using either a predefined ML-based approach[56] or an exploratory Bayesian framework[57]. To test whether snout elongation in spikefishes is best explained by a single adaptive optimum, one evolutionary shift, or two independent shifts, we conducted maximum-likelihood model fitting in OUwie[56] using continuous snout length and *a priori* defined snout-elongation regimes. We compared the following snout-elongation hypotheses: (a) OU1, the "No-shift hypothesis," in which spikefishes evolve under a single adaptive optimum for snout length; (b) OU2, the "One-shift hypothesis," in which extremely long-snouted spikefishes evolve under a distinct adaptive optimum relative to all other taxa; (c) OU3, the "Two-shift *Bathyphylax* hypothesis," in which extremely long-snouted species represent one shift and the genus *Bathyphylax* alone, herein the *Bathyphylax*-only hypothesis, represents a second, intermediate snout-length shift; and (d) OU3, the "Two-shift intermediate-clade hypothesis," in which extremely long-snouted species represent one shift and the broader clade including *Bathyphylax* + *Tydemania* + *Paratriacanthodes* represents a second, intermediate snout-length shift. We then evaluated these hypotheses using AIC weights. Lastly, because preliminary results demonstrated that the OU3 hypotheses were sensitive to the stochastic character map used as input, we performed a sensitivity test with 100 unique SIMMAP iterations per coding regime, refit the models, and evaluated the proportion of SIMMAPs that supported BM, OU1, OU2, or OU3.

Finally, we examined whether an exploratory Bayesian search of the continuous snout data could also identify the two shifts along branches consistent with our *a priori* groups using the package bayou[57]. Prior expectations for the OU parameters, alpha and $sigma^2$, were specified such that the minimum and maximum OUwie parameter estimates defined the boundaries of a 95% confidence interval under a half-Cauchy prior Because bayou requires a prior on the expected number of shifts, we ran analyses under four alternative shift-count priors (L = 1–4, where L is the prior expected number of shifts). For each prior setting, we ran two independent MCMC chains for 500,000 generations (eight runs total). Detailed results for L = 1 and L = 2 are shown in Figure S4A–H, and branch-level support across all four priors is summarized in Figure S5.

**Ordered accumulation of snout-associated traits.** Although ancestral state estimation informs the number and direction of snout elongation events, it does not address how complex snout morphologies arise. To test whether snout elongation evolved via additive, ordered anatomical changes, we implemented HyperMk[58], a custom R script that models trait evolution using hypercubic transition matrices while accounting for phylogenetic relatedness. HyperMk allowed all transitions with the criteria that only one character could change at a time (e.g., forbidding 000 → 011). In our hypothesis, the state '000' represents no snout elongation, and gains of '1' represent individual rearrangements associated with snout elongation. Support for our hypothesis required the presence/absence traits to accumulate in the following sequence: (i) the articulation and elongation of the metapterygoid-mesopterygoid and palatine-ectopterygoid (pterygoid closure; based on matrix characters 28 & 33); (ii) elongation and splitting of the vomer-parasphenoid complex (matrix character 11); and (iii) anterior rearrangement of the ectopterygoid and palatine (matrix characters 33 & 34). This analysis was repeated using both reversible and irreversible transition matrices and was run across all four phylogenetic trees. Model fits were next evaluated using AICw.

Given our small sample, size we evaluated whether HyperMk could perform adequately given our small dataset. Accordingly, we performed two complementary simulation tests: false-positive rate and statistical power. First, we fitted two alternative HyperMk models to the empirical data across the four trees to estimate transition-rate parameters for downstream simulations. These models included an (a) unconstrained independent-accumulation model, in which off-path transitions were allowed, and (b) a constrained ordered-accumulation model. The unconstrained model was specified with first.order = TRUE and reversible = FALSE, allowing all irreversible single-character gains. Under this model, one rate was estimated for the gain of each binary character and shared across all transitions involving acquisition of that character, including transitions not directly observed in the empirical tip data. Alternatively, the constrained model permitted only the empirically recovered pathway, 000 → 001 → 011 → 111, with all other transitions fixed to zero. We then used these estimated parameters to define unconstrained and constrained transition-rate matrices which served as inputs for data simulation using the function *simulate_mk_model* in castor[59], generating 250 replicate datasets under each generating model for every tree.

For the false-positive analysis, datasets were generated under the unconstrained model. These simulations produced stochastic, unconstrained patterns in which different lineages could accumulate characters through different allowable pathways. Because the unconstrained model generated the data, it was expected to fit well, whereas forcing the same data to fit the constrained pathway should generally produce a poorer fit. Therefore, if the constrained model nevertheless received the lower AIC, the simulated dataset was interpreted as randomly converging on a similar (or identical) ordered model and was counted as a false-positive model-selection result.

For the power analysis, datasets were generated under the correct constrained pathway, and HyperMk was then run again specifying the constrained and unconstrained transition matrix specifications. Because the constrained model generated the data, it was expected to be favored, whereas support for the unconstrained model represented a failure to detect the true pathway and was interpreted as a failed run (Figure S6).

**Spikefishes-snipefishes convergence**

**Convergence dataset and functional proxies.** To evaluate morphological and functional convergence between spikefishes and snipefishes, we assembled a comparative cranial dataset focused on elongated tubular snouts and the osteological organization of snout elements (Dataset 7). We defined long-snouted species as those in which the tubular snout is formed primarily by elongation of the ethmoid region and adjacent neurocranial elements (e.g., vomer, parasphenoid, ethmoid, and quadrate), with only minor contributions from jaw elements. In spikefishes, this condition occurs in *Halimochirurgus*, *Macrorhamphosodes*, and *Bathyphylax*. We further classified 'extreme' snout elongation as snout length >30% of standard length; this threshold includes snipefishes and the most elongate spikefishes, but excludes *Bathyphylax*, which represents an intermediate condition. This condition contrasts with 'longirostris' morphologies in which elongation is driven primarily by jaw elements (e.g., premaxilla/maxilla in butterflyfishes, swordfishes, and halfbeaks). Within spikefishes, snout length was categorized as short (up to ~15% of body length), intermediate (up to 28% of body length), or elongate (up to 48% of body length)[27,60].

This comparative dataset supplemented the 64 spikefish CT scans with five additional scans capturing head angle variation: three (*Halimochirurgus centriscoides,* n = 2; *H. alcocki*, n = 1; scanned at LACM), and two *Macroramphosus scolopax* (one from MorphoSource and one previously published[61]). Combined with the long-snouted spikefish scans described above, this dataset represents 25 species spanning a gradient of snout elongations and reducing biases associated with interspecific variation.

To assess convergence in feeding-related cranial mechanics, we compared spikefish cranial structures (Dataset 6) to the pivot-feeding apparatus of syngnathiform fishes, which couples rapid dorsoventral head rotation with suction and relies on coordinated hyoid depression and a mobile pharyngeal basket[61–64]. Because *in vivo* feeding recordings were unavailable for spikefish species, we used hyoid configuration as a proxy for cranial elevation and structural mobility. Mobility was coded based on the presence of both horizontal and vertical hyoid configurations among conspecific individuals (Figure S16). This coding was supported by new X-rays of three *Halimochirurgus centriscoides* (Faxitron MX-20), Smithsonian Institution X-rays and photographs (68 individuals; 12 spikefish species), ASIZP X-rays (2 individuals; 2 species), and published X-ray data[60], including *Bathyphylax pruvosti*, totaling 144 observations (Table S33).

**Comparative anatomy of snipefishes and spikefishes.** To identify anatomical similarities underlying snout convergence, we first conducted a qualitative comparative assessment of snout bones, focusing on bones that appeared to function as analogs, defined herein as structures with similar shape and position that may serve comparable functional roles, regardless of homology (Dataset 7). Based on this visual assessment, we identified candidate functional analogs between the two groups. We then quantified these similarities by applying the spikefish cranial landmarking scheme to snipefish specimens at each identified functional analog (Table S34). Landmark coordinates were subjected to local Procrustes superimpositions which included spikefishes and snipefishes, allowing direct comparisons of bone shape independent of size, orientation and position. Results were summarized using PCAs, and similarity between groups was evaluated based on the clustering of data points in morphospaces. To further assess functional convergence in feeding mechanics, we quantified jaw opening and closing mechanical advantages for both groups following Heiple et al[49]. Measurements were corrected for jaw width to allow for comparisons to be made among taxa with different cranial proportions and for those whose values were derived from standardized photographs.

To test whether convergence with the snipefishes was statistically significant under a broad phylogenetic framework, we expanded our taxonomic sampling to 300 species spanning Percomorpha. Phylogenetic hypotheses were grafted to include (a) the spikefish phylogeny estimated in this study, (b) a phylogeny of Syngnatharia, (c) a phylogeny of Tetraodontiformes, and (d) a percomorph-wide phylogeny. We subsampled this phylogeny to include overlapping species within the existing FishShapes dataset[65], representing 278 taxa, which we supplemented with 46 spikefish individuals representing 22 species (collectively, Dataset 9). In brief, FishShapes includes eight traits, each of which has been demonstrated to capture major axes of fish body shape, yet notably excludes snout length.

We then used the package *convevol* to estimate Ct metrics with the *calcConvCt* function and *convratsig* to estimate statistical significance[66,67]. We focused our post hoc interpretations on Ct1 metrics, which estimate the greatest historical distance between focal taxa in morphospace relative to their extant positions. We tested convergence using three unique inputs from conceptually distinct workflows that differed in their size-correction schemes and whether phylogenetically corrected morphospace coordinates were used as input to Ct, which itself incorporates phylogenetic relatedness. The workflows used to generate these inputs were (a) residuals of a phylogenetically informed linear model using trait ~ standard length, a non-phylogenetically informed morphospace, and then Ct; (b) volume-based size correction, non-phylogenetically informed morphospace, and then Ct; and (c) volume-based size correction, a phylogenetically informed morphospace, and then Ct.

### Functional validation

**Functional manipulation of fresh spikefish specimens.** Six fresh spikefish specimens were collected during the 2025 Coral Sea Frontier expedition aboard the RV *Investigator,* (CSIRO, Australia) (Dataset 8). We examined five species spanning a gradient of snout lengths: *Triacanthodes ethiops* (short snout), *Tydemania navigatoris* (short snout), *Bathyphylax bombifrons* (short to intermediate snout), *Macrorhamphosodes uradoi* (extremely long snout), and *Halimochirurgus centriscoides* (extremely long snout). All specimens were captured using Scampi or beam trawls at depths characteristic of spikefish occurrence (350–750 meters). Although specimens were deceased upon retrieval to the surface, prior to manipulations, individuals were placed in a large seawater filled megabin lined with a mesh lifting net and dosed with fish anesthetic (Aqui-S). This protocol minimized tissue stiffness and mechanical resistance during manipulation and enabled controlled assessment of cranial and hyoid mobility.

For each specimen, we manually manipulated the head and hyoid apparatus to evaluate the coupling between head elevation and hyoid depression, and recorded the approximate degree of head elevation. When possible, we compared spikefish four-bar linkage mechanics to pivot-feeding patterns reported for snipefishes[61] and seahorses[68]. Specifically, we assessed whether specimens (i) could elevate the head with the hyoid partially depressed, (ii) could raise the head while the suspensoria were adducted, and (iii) exhibited abrupt transitions of the hyoid between horizontal and vertical configurations during head elevation. To assess repeatability and minimize potential observer bias, manipulations were conducted independently by two investigators on the same specimens when conditions permitted, following a standardized protocol. Short-snouted species served as functional controls, allowing evaluation of whether observed mobility patterns were uniquely associated with elongate snouts rather than a shared character of spikefish cranial anatomy. Manipulations were video-recorded when feasible; however, not all individuals were recorded, and not all tests could be performed on every specimen due to variation in specimen condition and availability. Accordingly, these observations are interpreted as qualitative assessments of mechanical potential and constraint, rather than direct measurements of *in vivo* feeding kinematics.

Given the limited taxonomic sampling of fresh specimens and the challenges of sampling in high-pressure, time-sensitive environments, we repeated this experiment using cleared and stained specimens from a broad diversity of species (Dataset 10). Our sampling included additional teleost fishes with an emphasis on well-studied suction-feeding taxa (e.g., *Morone* and *Lepomis*) and syngnathiforms (including the snipefish *Macroramphosus*), seven non-spikefish tetraodontiform families (Tetraodontidae, Diodontidae, Molidae, Balistidae, Monacanthidae, Ostraciidae, and Triacanthidae), and five spikefish species spanning both subfamilies and all snout phenotypes, including the extremely long-snouted *Macrorhamphosodes* and *Halimochirurgus* (23 specimens representing 23 species; Table S28).

Head elevation was recreated as follows: specimens were laid horizontally, with the body and pectoral girdle held in a static position using probes. We then applied dorsal pressure to the upper jaw. Head elevation was considered present if the following conditions were met: (a) the head rotated along the posttemporal-neurocranial joint; (b) the hyoid bar was lowered; and (c) minimal bending occurred in the spine, as spinal bending could be confused with head elevation. Photographs were collected in neutral and head-elevated postures. To compare long-snouted spikefishes with snipefishes, we quantified change in head angle (Δhead), change in hyoid angle (Δhyoid), and kinematic transmission values (KT; Δhyoid/Δhead). In brief, KT represents the mechanical efficiency of four-bar linkages and is estimated as the ratio of observed rotation in the output link (hyoid depression) to observed rotation in the input link (head elevation), relative to the fixed link (i.e., the link between the cleithrum and head joint). We mapped the four-bar linkage following[69]. The points of the four-bar were defined as the posttemporal-neurocranial joint, the interhyal, the symphysis of the ceratohyals, and the ventralmost point of the cleithrum. Changes in head posture were calculated in Adobe Illustrator using the measure tool. Specifically, we projected lines through the parasphenoid, a straight bony segment that rotates with the head, and the hyoid bar, i.e., the output link, through the fixed link and extracted angles before and after manipulation.

### Specimens examined and description of morphological characters

**Specimen preparation.** External osteological characters were examined using cleared and stained (C&S), CT scanned (CT), and ethanol preserved (EtOH) specimens. Skeletal drawings of spikefish individuals from Tyler (1968)[27] and (1980)[28] were frequently referenced, with citations and page numbers provided. For fossil species, *†Carpathospinosus propheticus* and *†Prohollardia avita*, we coded the characters using photographs and drawings of the holotypes, as well as descriptions of the type (holotypes and paratypes) material from Tyler (1993)[29].

**List of specimens examined.** A total of 28 species were included in the morphological phylogenetic analyses, comprising all 24 valid extant spikefishes, two fossil spikefishes, and two outgroups from their sister group, Triacanthidae. Specimens used were provided by many institutions as follows: the Academy of Natural Sciences, Philadelphia (ANSP); National Museum of Natural History, Smithsonian Institution, Washington (USNM); Muséum National d'Histoire Naturelle, Paris (MNHN); Commonwealth Scientific and Industrial Research Organisation (CSIRO); Scripps Institution of Oceanography (SIO); National Science Museum, Department of Zoology, Tokyo, Japan (NSMT); South African Institute for Aquatic Biodiversity (SAIAB). Metadata regarding each specimen is formatted as follows: museum acronym and specimen voucher number, [number of individuals within lot], (preparation type). Voucher numbers may be represented multiple times per species, reflecting some combination of overlap across materials. For example, a CT scan and an EtOH specimen or a drawing and a cleared and stained individual. The following specimens are the basis for the morphological component of this study.

***Atrophacanthus japonicus***: ANSP 102139 [5] (C&S); ANSP 109016 [2] (CT); MNHN 2014-2112 [1] (EtOH); MNHN 2002-2990 [1] (EtOH); USNM 440413 [1] (EtOH); USNM 440419 [1] (EtOH); ANSP 103654 [1] (EtOH); ANSP 101239 [1] (Drawing Tyler 1968; pg. 357).

***Bathyphylax bombifrons***: CSIRO H 1181-10 [1] (CT); CSIRO H 6141-06 [1] (CT); MNHN 2004-103 [1] (CT); MNHN 1998-1095 [1] (EtOH); MNHN 2014-1288 [1] (EtOH); MNHN 2014-1256 [1] (EtOH); MNHN 2004-0105 [1] (EtOH); MNHN 2004-0103 [3] (EtOH); MNHN 2014-2097 [2] (EtOH); MNHN 2004-0104 [6] (EtOH).

***Bathyphylax omen***: ANSP 103313 [1] (CT).

***Bathyphylax pruvosti***: MNHN 2000-1179 [1] (C&S); MNHN 2000-4526 [1] (CT); MNHN 2000-4526 [1] (EtOH); MNHN 2005-0779 [3] (EtOH); MNHN 2003-1006 [5] (EtOH); MNHN 2001-0025 [2] (EtOH); MNHN 2000-4519 [3] (EtOH); MNHN 2003-0950 [3] (EtOH); MNHN 2000-4576 [4] (EtOH); Species description [N/A] (Santini 2006).

***Carpathospinosus propheticus*†**: ZPALWr. A/3000 [1] (Drawing Tyler 1993; pgs. 13-17).

***Halimochirurgus alcocki***: USNM 093474 [1] (C&S); USNM 306612 [1] (C&S); CSIRO H 6368-19 [1] (CT); CSIRO H 4074-08 [1] (CT); CSIRO H 1162-11 [1] (CT); USNM 306612 [4] (EtOH); USNM 93474 [1] (Drawing Tyler 1968; pgs. 225, 230).

***Halimochirurgus centriscoides***: ANSP 100864 [2] (C&S); CSIRO H 4070-27 [1] (CT); CSIRO H 4070-28 [1] (CT); ANSP 100828 [1] (CT); ANSP 100828 [1] (Drawing Tyler 1968; pgs. 225, 360).

***Hollardia goslinei***: BPBM 9655 [1] (C&S); ANSP 109175 [1] (CT); BPBM 14198 (EtOH).

***Hollardia hollardi***: ANSP 9775 [2] (C&S); ANSP 103303 [1] (C&S); ANSP 103299 [1] (C&S); ANSP 100867 [2] (C&S); USNM 280328 [2] (C&S); ANSP 98178 (CT); UF 88561 [1] (CT); FMNH 64394 [1] (CT); USNM 187810 [5] (EtOH); ANSP 100867 (Drawing Tyler 1968; pg. 353).

***Hollardia meadi***: ANSP 103246 [1] (CT); USNM 442062 [1] (EtOH).

***Johnsonina eriomma***: ANSP 103149 [2] (C&S); ANSP 103300 [2] (C&S); ANSP 97750 [2] (C&S); UF 88573 [2] (CT); ANSP 98177 [1] (CT); USNM 352028 [1] (EtOH); SIO 90- 23 [1] (EtOH); ANSP 103300 [1] (Drawing Tyler 1968; pg. 356).

***Macrorhamphosodes platycheilus***: USNM 306621 [1] (C&S); ANSP 100865 [2] (C&S); ANSP 100862 [1] (C&S); ANSP 100862 [2] (CT); CSIRO H 4070-31 [2] (CT); USNM 305526 [2] (EtOH); USNM 306627 [1] [EtOH]; ANSP 100865 [1] (Drawing Tyler 1968; pgs. 204, 215).

***Macrorhamphosodes uradoi***: USNM 101254 [1] (C&S); ANSP 109817 [2] (CT); CSIRO H 601-28 [1] (CT); CSIRO H 617-16 [1] (CT); MNHN 2004-0102 [1] (EtOH); MNHN 2014-0743 [1] (EtOH); ANSP 101254 [1] (Drawing Tyler 1986; pgs 204, 359).

***Mephisto albomaculosus***: NSMT 132271 [1] (CT) (Images Matsuura 2019; Bemis et al. 2020).

***Mephisto frasserbruneri***: USNM 350153 [1] (C&S); ZHM5629 [1] (CT); USNM 306629 [1] (EtOH); Review of genus [N/A] (Bemis et al. 2020).

***Parahollardia lineata***: ANSP 93375 [1] (C&S); ANSP 102144 [1] (C&S); ANSP 102145 [1] (C&S); ANSP 100473 [1] (C&S); ANSP 97637 [3] (C&S); USNM 287252 [4] (C&S); USNM 280326 [5] (C&S); ANSP 97637 [2] (CT); ANSP 93375 [1] (CT); ANSP 100473 [1] (CT); UF 87584 [2] (CT); FMNH 46686 [1] (CT); USNM 101319 [1] (EtOH); USNM 117283 [2] (EtOH); USNM 352029 [2] (EtOH); (Drawing Tyler 1980; pgs. 57-61).

***Parahollarida schmidti***: ANSP 100511 [2] (C&S); ANSP 100128 [4] (C&S); ANSP 103302 [2] (C&S); ANSP 100866 [5] (C&S); ANSP 100511 [2] (CT); UF 88579 [2] (CT); USNM 407215 [1] (EtOH); USNM 407220 [1] (EtOH); ANSP 103302 [1] (Drawing Tyler 1968: pg 352).

***Paratriacanthodes abei***: USNM 342571 [1] (CT); USNM 342571 [1] (EtOH).

***Paratriacanthodes herrei***: USNM 93302 [1] (C&S); USNM 93293 [1] (CT); USNM 93293 [1] (EtOH) USNM 93302 [1] (EtOH).

***Paratriacanthodes retrospinis***: ANSP 103285 [1] (C&S); SAIAB 82065 [1] (CT); SAIAB 82363 [1] (CT); CSIRO H 6574-08 [1] (CT); USNM 351336 [1] (EtOH).

***Prohollardia avita*†**: PALWr.A/2096 [1] (Drawing Tyler 1993; pgs 5-10). PALWr.A/2096 [1] (Photograph curtesy of Wroclaw University).

***Pseudotriacanthus strigilifer***: ANSP 100472 [1] (CT); SU 41732 [1] (Drawing Tyler 1968; pg. 363).

***Triacanthodes anomalus***: ANSP 101256 [1] (C&S); ANSP 101253 [1] (C&S); ANSP 101257 [1] (C&S); ANSP 166874 [4] (C&S); ANSP 101257 [2] (CT); FMNH 121112 [1] (CT); USNM 59693 [3] (EtOH); ANSP 101256 [1] (Drawing Tyler 1968; 354).

***Triacanthodes ethiops***: ANSP 103286 [1] (C&S); USNM 93487 [3] (C&S) ANSP 103286 [2] (CT); CSIRO H 6412-10 [1] (CT); USNM 93489 [1] (EtOH); USNM 93494 [1] (EtOH).

***Triacanthodes indicus***: NSMT 19726 [1] (CT); Species description [N/A] (Matsuura 1982).

***Triacanthodes intermedius***: NSMT 22373 [1] (CT); Species description [N/A] (Matsuura 1984).

***Tripodichthys blochii***: CSIRO H 7360-06 [1] (CT).

***Tydemania navigatoris***: ANSP 100861 [6] (C&S); ANSP 102137 [1] (C&S); ANSP 100863 [2] (C&S); CSIRO 6578-19 [2] (CT); CSIRO H 3156-34 [1] (CT); USNM 307542 [1] (EtOH); USNM 93473 [1] (EtOH); USNM 93470 [1] (EtOH); ANSP 100861 [1] (Drawing Tyler 1968; pg. 358).

**Description of characters:**

The primary goal of this character list is to assess the phylogenetic placement of both extant and fossil triacanthodid species. Characters were categorized by anatomical regions and further subcategorized into subregions following Tyler (1980). The cranial (skull) region included characters from the occipital, otic, orbital, ethmoid, jaw, palato-pterygoid, and opercular regions. The paired-fin girdle encompassed the pectoral and pelvic fin regions, while the dorsal, anal and caudal fin regions were also examined. Characters not assignable to these regions were categorized as miscellaneous.

### Cranial region

#### *Occipital*

1. **Supraoccipital shape:** (character based on Tyler 1968: 73, Arcila & Tyler 2017)
   (0) Dorsally rounded, growing to a blunted point posterodorsally, and without a laterally compressed triangular crest
   (1) Dorsally flattened, not growing to a blunted point posterodorsally, and with a laterally compressed triangular crest

2. **Supraoccipital crest: shape when supraoccipital is flattened**
   (0) Crest high, height of crest greater than or equal to the length at base of crest
   (1) Crest low, height of crest less than 0.5x the length at base of crest
   (-) Not applicable, supraoccipital is dome-like and lacks a crest

#### *Otic*

3. **Epiotic: articulation on the dorsalmost portion of the skull** (character based on Tyler 1980: 56)
   (0) Absent, not articulating on the dorsalmost part of skull
   (1) Present, articulating on the dorsalmost part of skull

4. **Epiotic: articulating with the frontal** (character based on Tyler 1968: 62)
   (0) Present, sphenotic not hindering articulation.
   (1) Absent, sphenotic hindering articulation

5. **Epiotic: projection on dorsoposterior edge**
   (0) Expanded posterodorsally, exhibiting rounded to triangular shapes and curved distally
   (1) Not expanded posterodorsally, exhibiting small and sharp points which are not curved distally

6. **Epiotic: shape of projection on dorsoposterior edge of epiotics when present**
   (0) Projection height comparable or greater than width posterior to cranium
   (1) Projection height less than to width posterior to cranium
   (-) Not applicable, projection are small sharp points

***Orbital***

7. **Eye: size compared to head length** (data recorded from Tyler 1986: 75, 87, 95, 100, 109, 117, 125, 135, 143, 146, 154, 167, 177, 181, 186, 198, 211, 218, 228; Matsuura 1982: 389; Matsuura 1984: 32 Matsuura 2018: 32; Santini 2006: 197)
   (0) Relatively small, less than 50% of head length.
   (1) Large, greater than 50% of head length

8. **Space anterior to orbit: shape between interorbital and mouth**
   (0) Straight to concave
   (1) Convex

***Ethmoid***

9. **Lateral ethmoid in relation to the ethmoid** (character based on Tyler 1980; 46, 78, 81)
   (0) Fused
   (1) Not fused, each bone a unique element.

10. **Vomer: bearing posterolateral extensions which suture to the ethmoid** (character based on Tyler 1980; 78)
    (0) Present
    (1) Absent

11. **Vomer: forming dorsal portion of snout**
    (0) Absent, situated laterally between palatines, not forming dorsal portion of snout
    (1) Present, situated dorsal to posterior portion of palatines, forming dorsal portion of snout

12. **Vomer: position in relation to ascending process of premaxillae**
    (0) Anterior edge ventral to posterior margin of ascending process of the premaxilla
    (1) Anterior edge distinctly posterior to ascending process of the premaxilla

13. **Vomer: position in relation to distal ends of maxillae**
    (0) Anterior edge ventral to posterior margin of the maxillae
    (1) Anterior edge distinctly posterior to margin of the maxillae

*Jaws*

**14. Teeth: shape**
(0) Square, heavy and block-like
(1) Rectangular, compressed and spatulate
(2) Conical, tapering to sharpened points

**15. Second row of teeth situated behind the dentary and premaxillary teeth** (character based on Tyler 1968; 73)
(0) Present
(1) Absent

**16. Teeth: dorsal surface when spatulate**
(0) Dorsal surface of teeth exclusively flat, not varying throughout dentition
(1) Dorsal surface of teeth flat or pointed, varying throughout dentition
(-) Not applicable, teeth are not spatulated

**17. Teeth: spacing when conical**
(0) Spacing narrow, the gaps between many times smaller than the height of the teeth (Alternatively, teeth larger and not buried greatly within the gums*)*
(1) Spacing great, the gaps between teeth as wide as teeth in height or greater (Alternatively, teeth small and buried greatly within the gums)
(-) Not applicable, teeth are not conical.

**18. Maxillae: dorsal margin in relation to symphysis of ascending process of premaxillae**
(0) Dorsal maxillae are not close, and not located ventrally to symphysis of the ascending process of premaxillae
(1) Dorsal maxillae are close, and located ventral to the symphysis of ascending process of premaxillae.

**19. Maxilla: anteroventral expansion**
(0) Absent
(1) Present

**20. Maxilla dorsal condyle: width-to-depth ratio**
(0) Width is approximately equal to depth
(1) Width is approximately 1.5 to 2.5x the depth
(2) Width is approximately 3.3x to 3.5x the depth
(3) Width is 4x or greater the depth

**21. Maxilla dorsal condyle: shape**
(0) Laterally compressed and plate-like

(1) Rounded and rod like

**22. Mouth: orientation**
(0) Terminal to slightly supraterminal, mouth opening forward
(1) Distinctly supraterminal, mouth opening dorsally (includes *Macrorhamphosodes* even when mouth is turned laterally)

**23. Mouth orientation: when distinctly dorsal** (observed only in species with ontogenetic series)
(0) Not twisting to left or right with ontogeny
(1) Twisting to left or right with ontogeny

**24. Quadrate: angle formed by posteriormost points and anteriormost point**
(0) Quadrate angle greater than 30 degrees and less than 40 degrees
(1) Quadrate angle less than 30 and greater than 10 degrees
(2) Quadrate angle less than 10 degrees
(3) Quadrate angle greater than 40 degrees

***Palato-pterygoid***

**25. Mesopterygoid** (character based on Tyler 1980; 46)
(0) Absent
(1) Present

**26. Mesopterygoid: medial ridge**
(0) Present, bearing a distinct medial ridge
(1) Absent, lacking a distinct medial ridge
(-) Not applicable, mesopterygoid absent

**27. Mesopterygoid: articulating with palatine anteriorly**
(0) Present, posterior palatine in contact with anterior mesopterygoid
(1) Absent, posterior palatine not in contact with anterior mesopterygoid
(-) Not applicable, mesopterygoid absent

**28. Metapterygoid: in contact with mesoptyergoid across entire dorsal edge.**
(0) Absent, mesopterygoid not articulating with entire dorsal edge
(1) Present, mesopterygoid articulating with entire dorsal edge
(-) Not applicable, mesoptyergoid absent

**29. Ectopterygoid: obliqueness to horizontal axis**
(0) More oblique, the ventral edge approximately 30 degrees or more above horizontal axis
(1) Not oblique, the ventral edge less than 30 degrees above horizontal axis

**30. Ectopterygoid: triangularly shaped**

(0) Present, the ectopterygoid with three strong points
(1) Absent, ectopterygoid with four strong points

**31. Ectopterygoid: dorsal edge when triangularly shaped**

(0) Lacking a recurved posterior point
(1) Bearing a recurved posterior point
(-) Not applicable, ectopterygoid is not triangular

**32. Ectopterygoid: height to length when triangular**

(0) Width less than 2.4x the height
(1) Width 2.8 to 3.6x the height
(2) Width 3.9 to 4.7x the height.
(3) Width at least 5x greater than height.
(-) Not applicable, ectopterygoid is not triangular.

**33. Palatine: articulating with the ectopterygoid**

(0) Absent, the palatines and ectopterygoid separated
(1) Present, the palatines and ectopterygoid articulating dorsally or anteriorly

**34. Palatine: when articulating with ectopterygoid**

(0) Articulating ventrally with the ectopterygoids
(1) Articulating anteriorly with the ectopterygoids

**35. Palatine: relation to the anteriormost point of lateral ethmoid**

(0) Posteriormost end situated posterior to lateral ethmoid
(1) Posteriormost end situated approximately in line with or anterior to

**36. Palatines: articulating dorsally in the anterior portion of snout** (character based on Tyler 1968; 204, 225)

(0) Absent, situated laterally and not expanded dorsally for articulation
(1) Present, situated laterally and expanded dorsally allowing for articulation

**37. Palatine: shape**

(0) Sigmoidal
(1) Rod-like
(2) Round
(3) Square

**38. Palatine: bearing a posterior point on the dorsalmost edge when sigmoidal**

(0) With a sharp posterior point on the dorsalmost surface
(1) Without a sharp posterior point on the dorsalmost surface
(-) Not applicable, palatine shape not sigmoidal

*Opercular region*

**39. Gill opening: depth compared to base of pectoral fin**
(0) Equal or similar in size (0.5-1.5x greater) than pectoral fin base
(1) Larger (+2x) the pectoral fin base height

**Paired-fin girdles**

***Pectoral fin***

**40. Cleithrum: widening posteriorly at level of scapula**
(0) Present, cleithrum narrowing dorsally and then broadening in the midsection
(1) Absent, cleithrum narrowing but not broadening in the midsection

**41. Cleithrum: shape when expanded at level of scapula**
(0) Ventral edge of cleithrum tapering gradually to a point
(1) Ventral edge of cleithrum tapering rapidly to a point
(-) Not applicable, posterior expansion absent or only very slightly expanded

**42. Cleithral process of the pelvis: shape of anteroventral edge**
(0) Concave
(1) Convex

**43. Cleithral process of the pelvis: shape of projection anterodorsal to pelvic spine insertion**
(0) Anterolateral flange present
(1) Anterolateral flange absent dark

**44. Anterior flange of the cleithral process of the pelvis: shape**
(0) Oblong, much longer than wide
(1) Roundish, length only slightly greater than width
(2) Rectangular, wider than long
(-) Not applicable, flange absent

**45. Anterior flange of the cleithral process of the pelvis: narrow at base**
(0) Absent, flange not narrowing distinctly at base
(1) Present, flange narrowing distinctly at base
(-) Not applicable, flange absent

**46. Anterior flange of the cleithral process: angle of projection relative to anteroposterior axis**
(0) Less than 30˚ above horizontal

(1) 30˚ or more above horizontal
(-) Not applicable, flange absent

**47. Anterior flange of the cleithral process of the pelvis: size**
(0) Very long, the flanges length approximately ½ the length of the cleithral pelvis
(1) Not very long, the flange less than ½ the length of the cleithral pelvis
(-) Not applicable, flange is absent

***Pelvic fin***

**48. Process of the pelvic bone: ventral surface** (character based on Tyler 1968: 62)
(0) Rounded and only slightly tapering to a point posteriorly
(1) Flattened and tapering distinctly to a point posteriorly

**49. Posterior process of the pelvis: length into width** (data recorded from Tyler 1968: 75, 87, 95, 101, 110, 117, 126, 136, 143, 146, 155, 167, 177, 182, 187, 199, 211, 220, 228; Tyler 1993: 4, 14; Matsuura 1982: 389  Matsuura 1984: 32 Matsuura 2018: 32; Santini 2006: 187)
(0) Pelvis exceedingly narrow, the length at least 6.5x the width
(1) Pelvis narrow, length at least 3x the width or greater
(2) Pelvis broad, the length only 1.5 to 3x greater than the width

**50. Pelvis: region dorsal to the locking mechanism of the pelvic spine when ventral surface of pelvis is flattened.**
(0) Only slightly raised
(1) Raised area is prominent
(-) Not applicable, ventral surface of pelvis is rounded

**51. Pelvic spines: retrorse barbs present on adherent scales**
(0) Retrorse barbs shorter than ½ width of the pelvic spines or absent
(1) Retrorse barbs length approximately ½ or greater than the width of the pelvic spines

**52. Second pelvic ray: ontogenetic tendency** (as noted from Tyler 1968: 75, 96, 135, 142, 151, 152, 153, 167, 185, 195, 217; Tyler 1980: 79; Santini 2006)
(0) Second pelvic ray becoming rudimentary and eventually disappearing beneath the skin
(1) Second pelvic ray becoming small but never buried beneath the skin
(2) Second ray always absent

**Dorsal and anal fins**

***Dorsal fin***

**53. Dorsal spines: nature of decreasing size** (character based on Tyler 1968: 60, 235)

(0) Spines taper in size rapidly, the first spine a robust rod and the second spine considerably more thin
(1) Spines taper gradually, the second spine shorter but similar in profile.

54. **Spiny dorsal fin membrane scales when scales are non-cruciform** (data recorded from Tyler 1968: 80, 89, 97, 102, 121, 127, 137, 144, 147, 168, 179, 182, 187, 202, 212, 222, 229, 102, 111; Tyler 1993: 20; Matsuura 1982: 387; Matsuura 1984: 32; Matsuura 2018: 31)
(0) Scales absent to few, scattered along the basal margin of the membrane
(1) Scales present
(-) Not applicable, scales are non-cruciform

55. **Dorsal spine six: size of** (coded from EtOH preserved specimens)
(0) Rudimentary, only slightly exposed or buried beneath scales
(1) Not rudimentary, clearly extended dorsally above the scales

56. **Dorsal spine five: size** (coded from EtOH preserved specimens)
(0) Rudimentary, only slightly exposed or buried beneath scales
(1) Small, between ¼ and ½ the length of dorsal spine 4.
(2) Large, greater than ½ length of dorsal spine 4.

57. **Dorsal spine four: size** (coded from EtOH preserved specimens)
(0) Rudimentary, only slightly exposed or buried beneath the scales
(1) Small, less than ½ the length of dorsal spine 3
(2) Large, at least ¾ the size of the dorsal spine 3

58. **Dorsal spine three: size** (coded from EtOH preserved specimens)
(0) Rudimentary, only slightly exposed or buried beneath the scales
(1) Spine is well developed in size, only slightly smaller than dorsal spine 2

59. **First spiny dorsal pterygiophore: shape of entire anterior edge**
(0) Smooth, lacking a sharp turn in the midsection
(1) Angular, bearing a sharp turn in the midsection

60. **First spiny dorsal pterygiophore: shape of anterodorsal edge when pterygiophore is angular**
(0) Straight to slightly convex or concave
(1) Edge strongly concave
(-) Not applicable, anterior side of pterygiophore is smooth

61. **First spiny dorsal pterygiophore: depth to width ratio when pterygiophore is angular**
(0) Squarish, width and height approximately equal
(1) Narrow, depth considerably greater than width
(-) Not applicable, anterior side of pterygiophore is smooth

**62. First spiny dorsal pterygiophore: anterodorsal edge when pterygiophore is smooth**

(0) Straight
(1) Convex
(-) Not applicable, anterior pterygiophore bears a strong point

**63. First spiny dorsal pterygiophore: posterior projection**

(0) Greatest pterygiophore depth tapers to a shallow expansion posteriorly
(1) Greatest pterygiophore depth tapers broad plate-like projection posteriorly

**64. Dorsal edge of pterygiophore: with a thick section forming the interlocking mechanism for dorsal spine erection**

(0) Absent, dorsal edge without a thick ridge
(1) Present, dorsal edge with a thick ridge

**65. Dorsal edge of pterygiophore anterior to the first dorsal spine: length of thickened ridge**

(0) Thickened ridge length equal to or only slightly greater than pterygiophore
(1) Thickened ridge length clearly greater than pterygiophore
(-) Not applicable, thickened dorsal ridge is absent

**66. Dorsal edge of pterygiophore anterior to the first dorsal spine: angle of thickened ridge**

(0) Anterior side is not curved ventrally, tapering anteriorly
(1) Anterior side curved, tapering ventrally
(-) Not applicable, thickened dorsal ridge is absent

**67. Dorsal edge of pterygiophore posterior to the first dorsal spine: shape of thickened ridge**

(0) Ridge rounded and raised; raised area approximately as tall as wide
(1) Ridge not rounded and raised; raised area taller than wide
(-) Not applicable, thickened dorsal ridge is absent

**68. Second spiny dorsal pterygiophore: shape**

(0) Pterygiophore highly reduced, a horizontal bone with ‘V’-shaped ventral projection or only a slight ventral projection
(1) Pterygiophore angular, the parallel dorsolateral edges forming the anterior and posterior sides of pterygiophore are approximately equal in length before tapering to a ‘V’-shaped ventral point.
(2) Pterygiophore less angular, the parallel dorsolateral edges forming the anterior and posterior sides of pterygiophore are not equal in length, still forming a ‘V’-shape ventrally
(3) Pterygiophore triangular, lacking parallel dorsolateral edges before tapering to a ‘V’-shape ventrally

**69. Spiny dorsal pterygiophores 2 and 3: connection dorsally**

(0) Not tightly connected on dorsal edge

(1) Tightly connected on dorsal edge

**70. Spiny dorsal pterygiophores 3 and 4: connection of dorsally**

(0) Not tightly connected on dorsal edge; separated or only articulating
(1) Tightly connected on dorsal edge

#### *Anal fin*

**71. First anal pterygiophore: shape of cross section** (character based on Tyler 1980; 56):

(0) Cross sectional area is '+' shaped; shaped by broadening lateral edges and perpendicular to these edges, a laterally compressed crest expanded anteriorly.
(1) Cross section area is 'T' shaped; shaped by broadening lateral edges laterally and the absence of a laterally compressed crest anteriorly

**72. Haemal spines: counts between the anal pterygiophores**

(Note, if the first anal pterygiophore is fused to two haemal spines, the fused spines are included in the count. Any neural spines beyond the further posterior boundary of an anal fin pterygiophore are not counted)

(0) Five
(1) Six
(2) Seven

### Caudal region

#### *Caudal fin*

**73. Hypural plate: number of hypurals that form majority of caudal fin**

(0) Two
(1) Three
(2) Four

### Miscellaneous

**74. Scales** (character based on Tyler 1980; 79)

(0) Cruciform, bearing perpendicular rows of raised spinules
(1) Non-cruciform, lacking perpendicular rows of raised spinules

**75. Upper lip scales when scales are non-cruciform** (data recorded from Tyler 1968: 79, 121, 127, 137, 144, 147, 168, 179, 182, 187, 202, 214, 223, 230; Matsuura 1982: 387; Matsuura 1984: 32; Matsuura 2018: 31)

(0) Present, bearing patches of isolated scales
(1) Absent, lacking patches of isolated scales
(-) Not applicable, scales are cruciform

**76. Lower lip scales when scales are non-cruciform** (data recorded from Tyler 1968: 79, 121, 127, 137, 144, 147, 168, 179, 182, 187, 202, 214, 223, 230; Matsuura 1982: 387; Matsuura 1984: 32; Matsuura 2018: 31)

(0) Present, bearing patches of isolated scales
(1) Absent, lacking patches of isolated scales
(-) Not applicable, scales are cruciform

**77. Lips: shape**

(0) Small
(1) Enlarged and spongy
(2) Reduced

**78. Average body depth in adults** (coded from Tyler 1968: 75, 87, 95, 101, 110, 120, 126, 136, 143, 146, 154, 167, 177, 181, 186, 198, 211, 220, 228, 291; Matsuura 1982: 389; Matsuura 194: 32; Matsuura 2018: 32; Santini 2006: 197; Tyler 1993: 4, 14)

(0) Standard length is at most 2.5x greater than body depth
(1) Standard length is at least 3x greater than body depth

**79. Body coloration in live specimens**

(0) Silver with brown to yellow patches
(1) Red to yellow

**80. Color pattern when body color is red to yellow** (data recorded from Tyler 1968: 80, 91, 97, 104, 112, 121, 128, 137, 144, 148, 169, 187, 202, 214, 223, 230; Matsuura 2018: 31; Bemis et al., 2020: 23; Matsuura & Tyler 1997: pg. 174)

(0) Striped
(1) Irregular/reticulate
(2) Eye spot
(3) Scattered freckle-like spots
(4) Colored red dorsally fading to a pale hue ventrally.

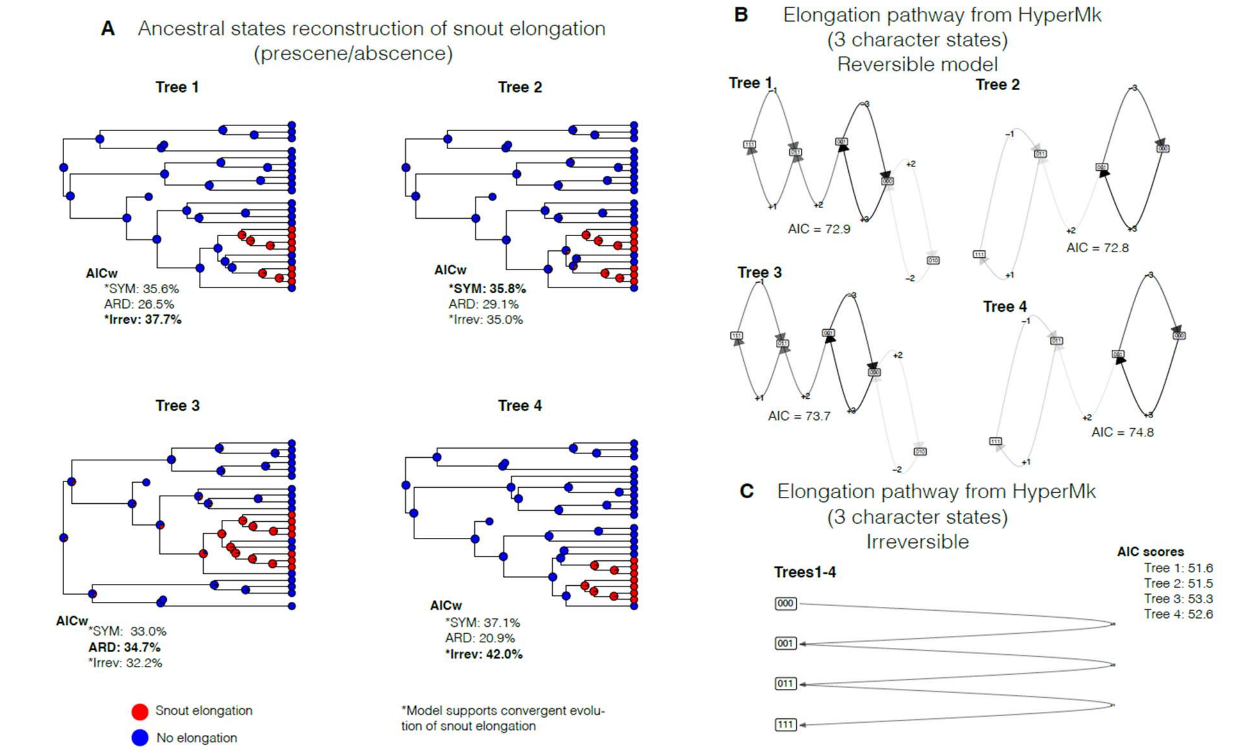


**Figure S1. Character evolution of snout elongation in spikefishes across multiple models.** (A) Ancestral state reconstructions of snout elongation across four phylogenetic trees. AIC weights for each of the models tested (SYM: symmetrical, ARD: all rates different, and Irrev: irreversible model designed with a custom Q matrix), with the best model bolded. Asterisks (*) denote models supporting convergent evolution of snout elongation. Blue circles indicate the absence of snout elongation, and red circles indicate its presence. (B) Character evolution pathways under a reversible accumulation model (HyperMk package) for four phylogenetic trees, with AIC values reported for each. (C) Character evolution pathways under an irreversible accumulation model. Plots are identical across trees, with AIC scores approximately 20 units lower than those in the reversible model.

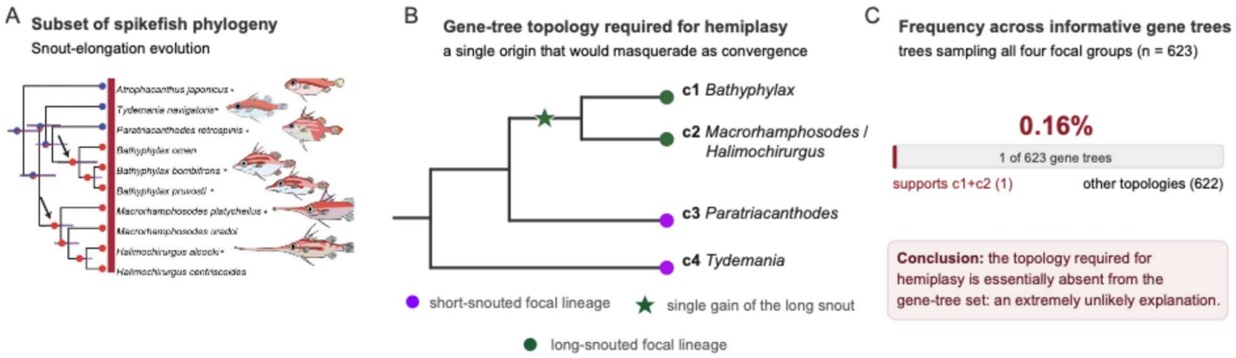

**Figure S2. Hemiplasy is an unlikely explanation for convergent snout elongation in spikefishes. A.** Spikefish subtree showing two independent origins of the elongate snout (arrows). Tip and node symbols indicate snout state (blue: short; red: elongated); the asterisks denote species shown in the illustrations. **B.** The gene-tree topology required for hemiplasy: a single snout-elongation gain (star) uniting the two elongated-snouted focal groups, c1 (*Bathyphylax*, intermediate-snouted), and c2 (*Macrorhamphosodes*/*Halimochirurgus*, extremely long-snouted), to the exclusion of the short-snouted groups c3 (*Paratriacanthodes*) and c4 (*Tydemania*). **C.** This topology occurs in only 1 of the 623 gene trees sampling all four focal groups (0.16%), indicating that hemiplasy is highly unlikely.

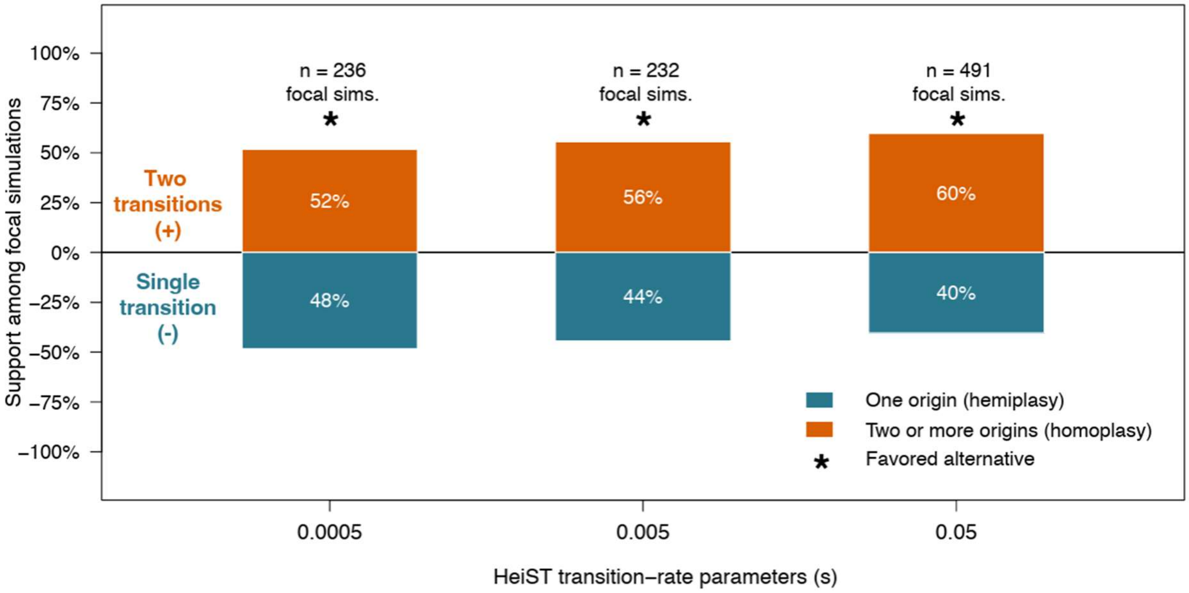


**Figure S3 Hemiplasy test using HeiST:** HeiST simulations evaluating alternative evolutionary histories for the observed distribution of snout elongation. Bars show the proportion of focal simulations supporting hemiplasy (one origin; blue, below zero) or homoplasy (two or more origins; orange, above zero) across three trait-transition-rate parameters (s). Within each rate setting, proportions sum to 100%; n denotes the number of focal simulations. Asterisks indicate the alternative with greater support at that rate.

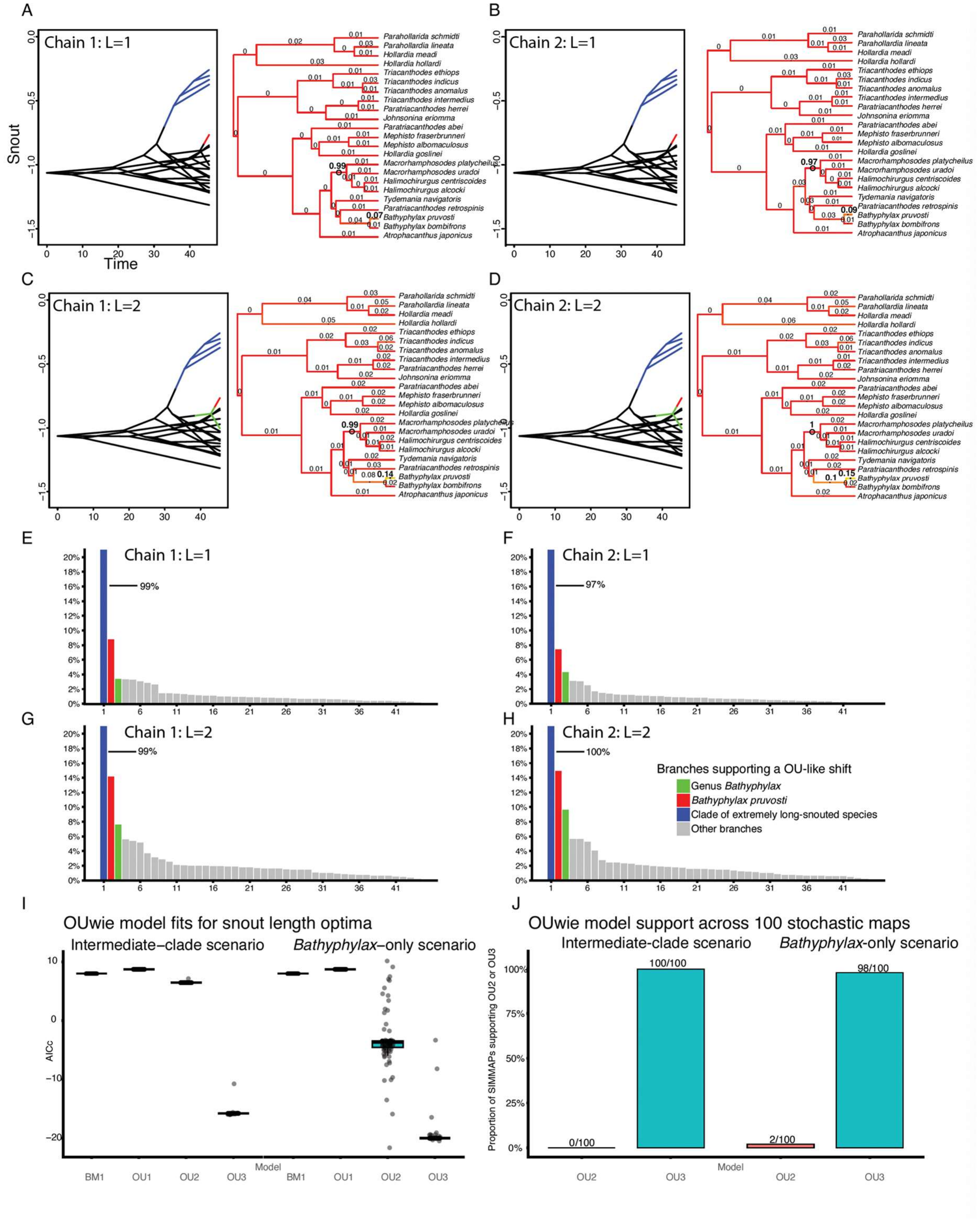


**Figure S4. Snout-elongation shifts inferred from continuous snout data under Bayesian and ML frameworks. A–D.** bayou reversible-jump analyses of relative snout length across two independent MCMC chains and two different prior hypotheses (L-values) regarding the expected number of shifts. Each result includes phenograms (left; colored branches indicating shifts to new optima, with extremely long-snouted spikefishes and the next-most likely transition colored) and per-branch posterior probabilities (pp) (right), indicating the proportion of samples that recovered a shift along a given branch. **E–H.** pp from each run shown in A–D, sorted by magnitude. *Bathyphylax pruvosti* is consistently the second-most-supported transition, followed by a transition at the base of the entire genus. Plots are truncated at 20% pp; lines extending from the blue boxes indicate the pp associated with the extremely long-snouted clade. **I.** AIC values estimated across 100 stochastic character maps (i.e., SIMMAPs) to evaluate the recovery rate of OU3 (i.e., two shifts toward long-snoutedness) for the 'intermediate-clade' (*Paratriacanthodes* + *Tydemania* + *Bathyphylax*, collectively representing a shift) and '*Bathyphylax*-only' (*Bathyphylax*-only representing a shift) hypotheses. **J.** Model sensitivity across 100 SIMMAPs, showing whether Brownian motion, OU1 (no shift, single optimum), OU2 (one shift, two optima), or OU3 (two shifts, three optima) is the best-supported model.

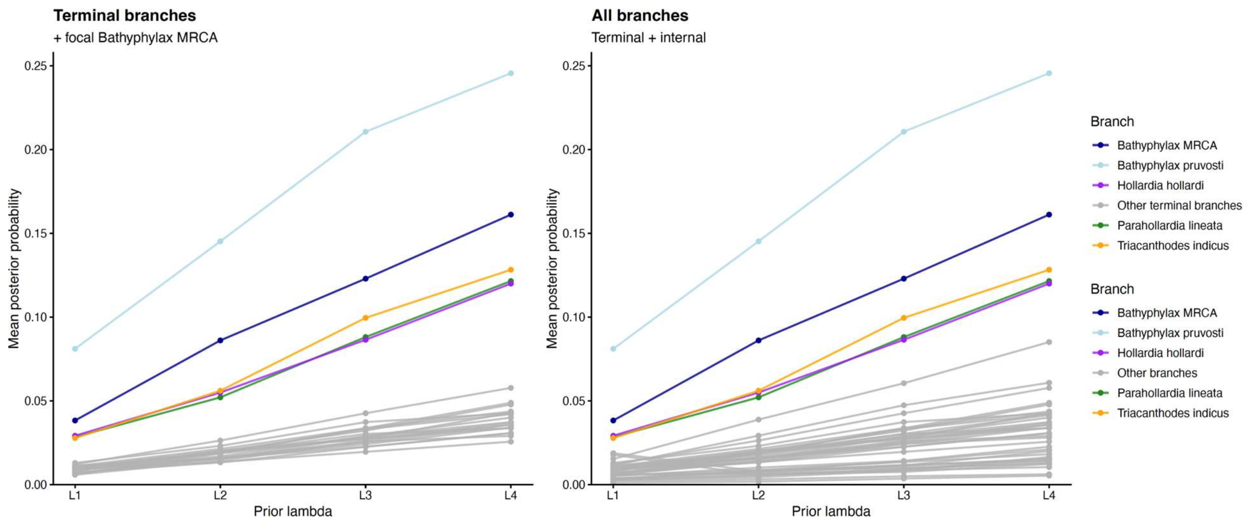


**Figure S5.** Branch-level bayou support for snout-elongation shifts across four shift-count priors. Per-branch posterior probabilities of an adaptive shift in relative snout length from reversible-jump OU analyses run under four shift-count priors (L = 1–4), each with two independent MCMC chains (eight runs total). Across all priors and chains, the extremely long-snouted clade (*Halimochirurgus* and *Macrorhamphosodes*) received the strongest support (branch posterior probabilities ≈ 0.97–1.0), and the branches leading to *Bathyphylax pruvosti* and to the base of the genus *Bathyphylax* consistently carried the next-highest posterior probabilities (7–25% and 4–16% of posterior samples, respectively), with absolute support increasing with the prior mean but rank order remaining stable. No branch elsewhere in the tree exceeded the support of these branches under any prior or chain.

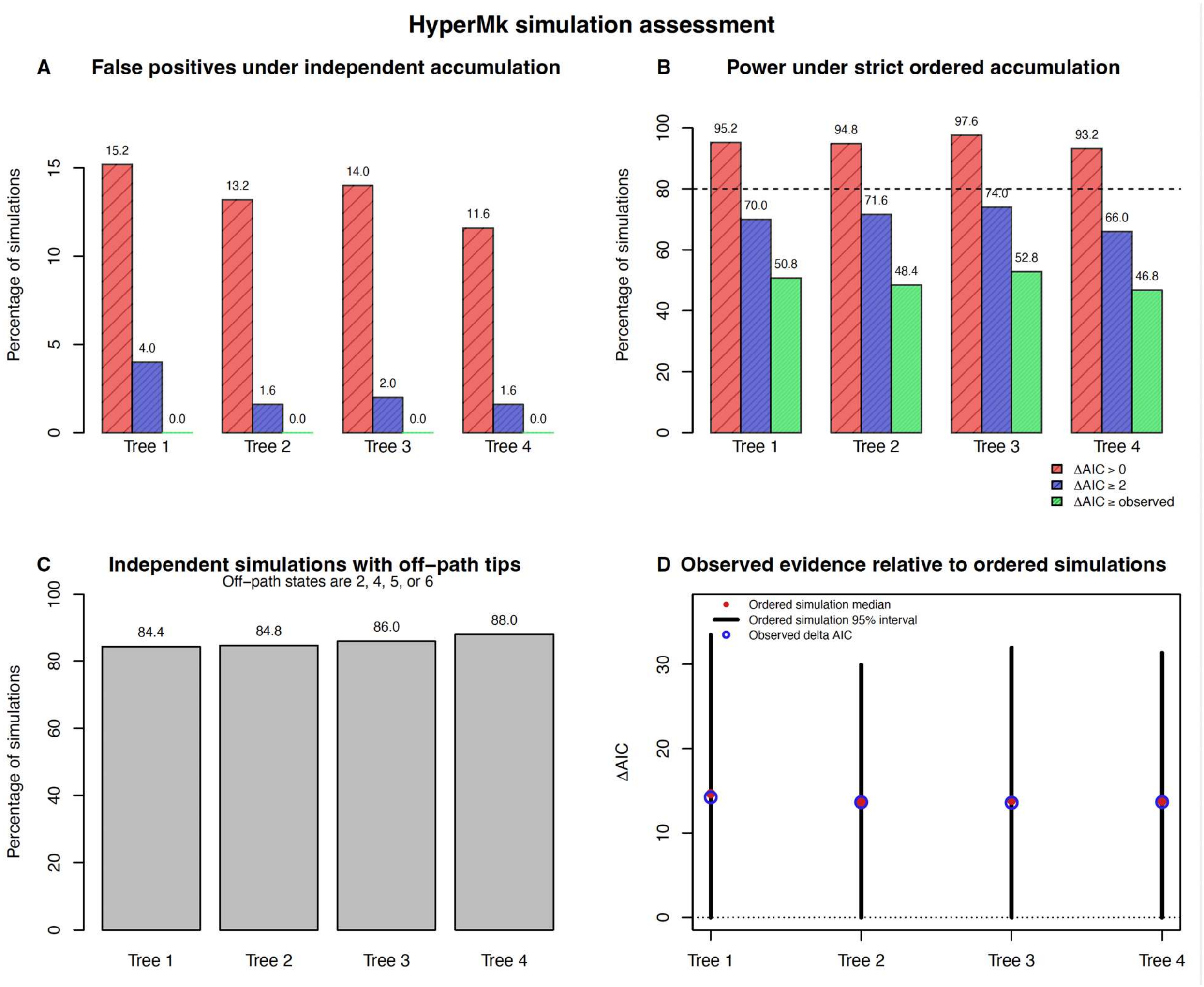


**Figure S6. Simulation-based evaluation of false-positive rates and statistical power in HyperMk.** For each of four empirical trees, transition-rate parameters were estimated under an unconstrained independent-accumulation model (first.order = TRUE and reversible = FALSE; all irreversible single-character gains allowed, with one shared rate per acquired binary character) and a constrained ordered-accumulation model, permitting only the empirically supported pathway (000 → 001 → 011 → 111). The resulting Q matrices were used to simulate 250 replicate datasets under each generating model per tree (1,000 per model), and every dataset was refit with both models. Support was measured as ΔAIC = AIC (unconstrained) − AIC (constrained), with positive values favoring the constrained model. **A**. Under the unconstrained model, the constrained model was favored in 11.6–15.2% of datasets (red) and favored with ΔAIC ≥ 2 in 1.6–4.0% (blue), but none reached the empirically observed ΔAIC (green). **B.** Under the constrained model, the constrained model was favored in 93.2–97.6% of datasets (red), with ΔAIC ≥ 2 in 66.0–74.0% (blue); 46.8–52.8% produced fits at least as strong as the empirical values (green). Dashed line indicates an 80% power benchmark. **C.** Among unconstrained simulations, 84.4–88.0% contained at least one tip state outside the constrained pathway, confirming that alternative accumulation sequences were generated. **D.** Empirical ΔAIC (blue) fell within the 95% interval and near the median of the constrained-model distribution for every tree (the same comparison summarized by the green bars in B), indicating the empirical fit was neither spuriously low nor high. These results indicate high power to detect the constrained pathway and a low probability of obtaining similarly strong support when the unconstrained process generated the data.

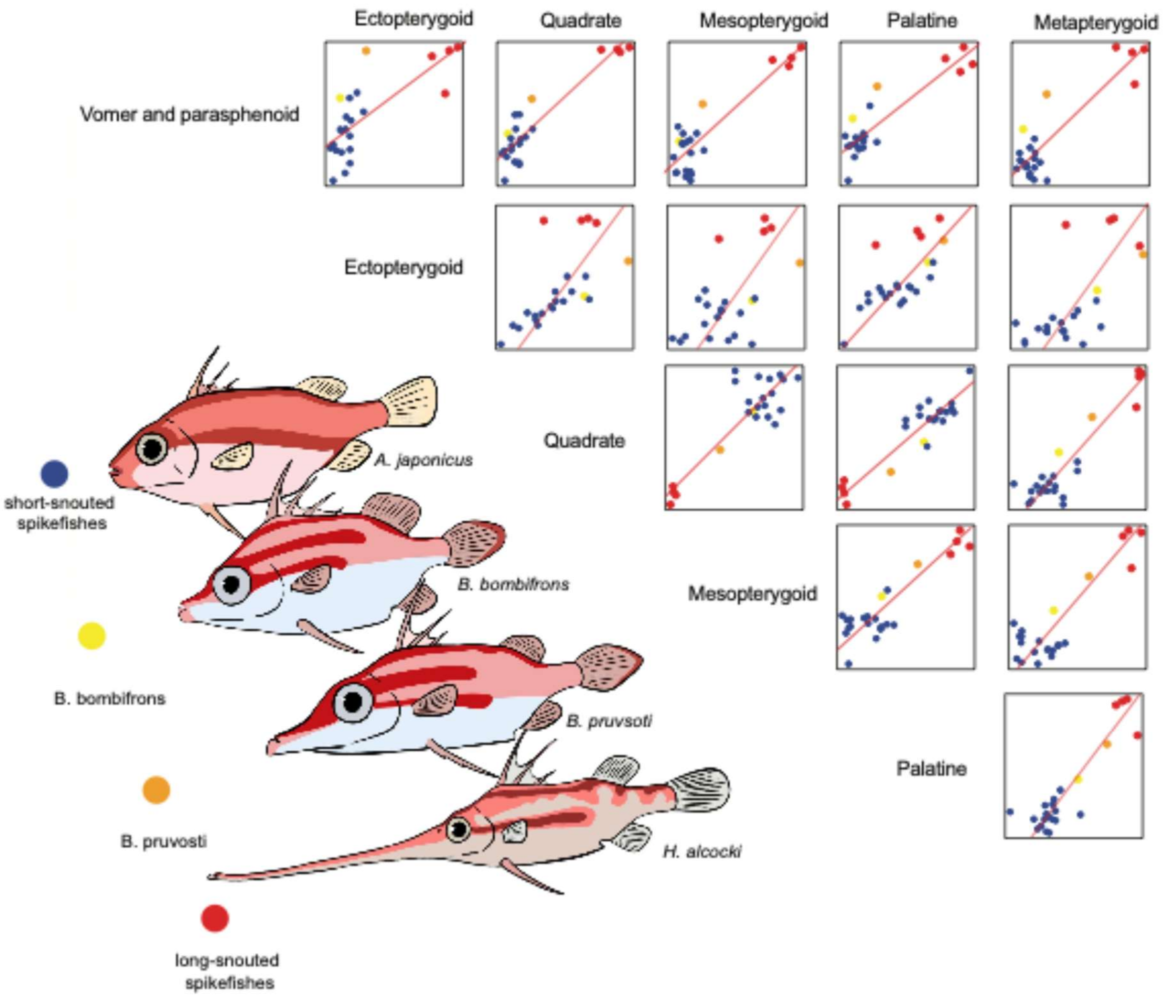


**Figure S7. Pairwise integration of snout bones using two-block partial least squares (2BPLS).** Pairwise integration calculated through 2BPLS regressions, presented in a matrix format. Columns and rows are labeled with the individual bones analyzed in each 2BPLS comparison. When points are organized as blue, yellow, orange, red, the integrated shape pathway of snout elongation is supported.

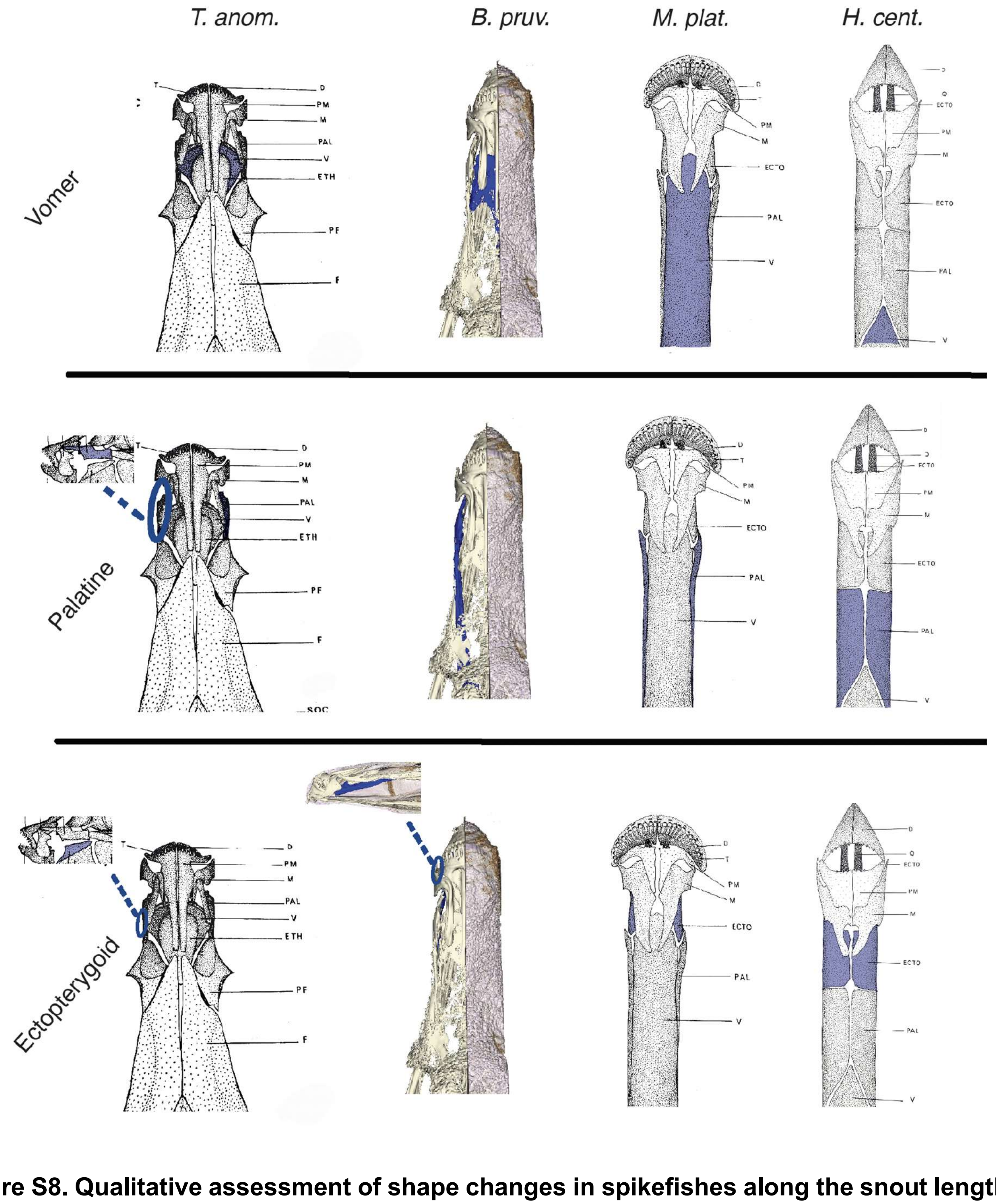


**Figure S8. Qualitative assessment of shape changes in spikefishes along the snout length continuum.** Four species *Triacanthodes anomalus* (*T.anom;* short snout)¸ *Bathyphylax pruvosti* (*B. pruv;* intermediate snout)*, Macrorhamphosodes platycheilus* (*M. plat*; second most elongated)*, Halimochirurgus centriscoides* (*H. cent*; longest snout), and changes to the shapes vomer, palatine, and ectopterygoid. Drawings used from Tyler 1968[27]. Shorthand for bones are as follows: dentary (D), premaxilla (PM), maxilla (M), palatine (PAL), vomer (V), ethmoid (ETH), pre-frontal/lateral ethmoid (PF), frontal (F).

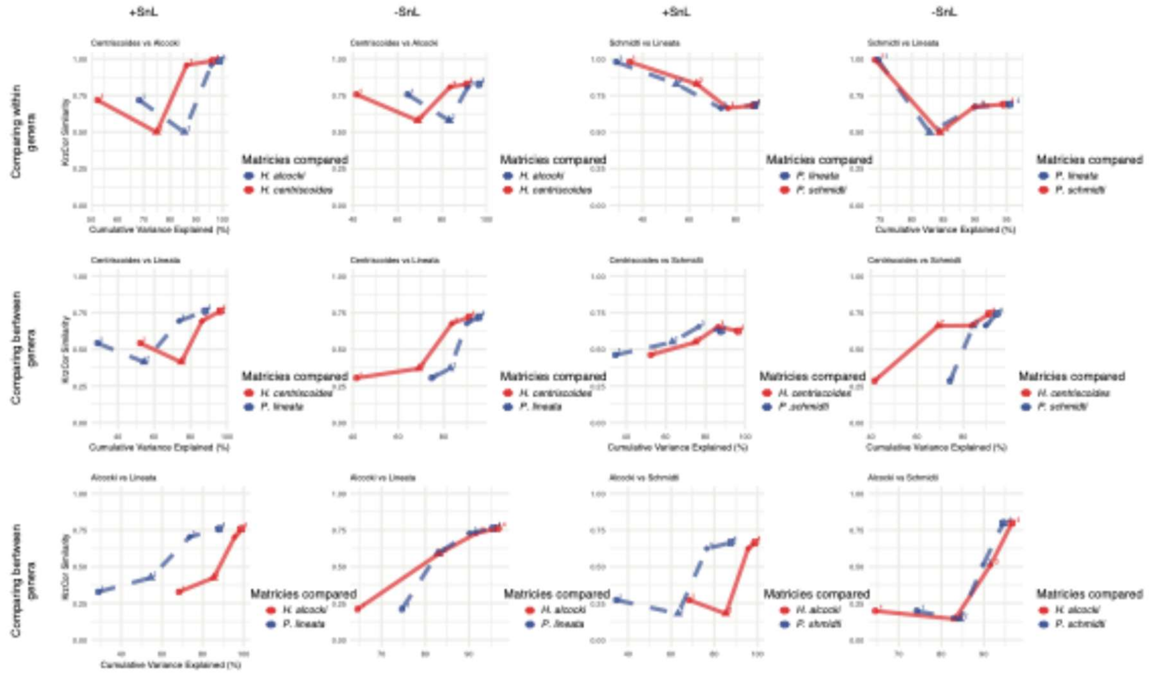


**Figure S9. Matrix similarity is greater among species with shared phenotype.** Krzanowski correlation indices were calculated between the PC1–PC4 subspaces of each species' P matrix and compared (colors distinguish the two matrices within each panel and are not consistent across comparisons). Line plots show the proportion of variance explained by either species's **P**. Higher Krzanowski correlation values indicate greater alignment of the leading PC subspaces between **P** matrices Similarity estimates are shown for shared phenotype (top; i.e., comparing long-snout with long-snout and short with short. species (top), long-snouted species *H. centriscoides* compared with both short-snouted (middle), and long-snouted *H. alcocki* with short-snouted (bottom). Columns indicate size correction performed with head length included (+SnL, including snout) or excluded (-SnL, excluding snout).

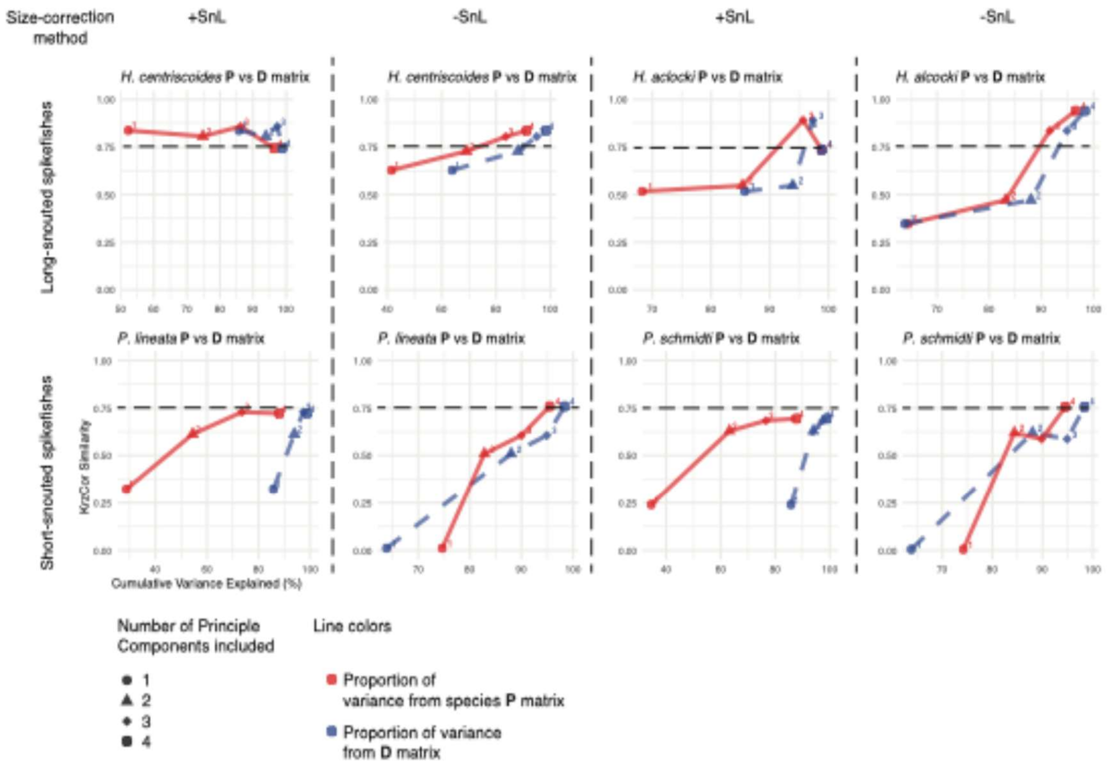


**Figure S10. Long-snouted species P matrices are more similar to clade-wide D matrix than short-snouted.** Krzanowski correlation indices were calculated between the PC1–PC4 subspaces of each species' P matrix (red) and the clade-wide **D** matrix (blue). Line plots show the proportion of variance explained by each PC for **P** and **D**. Higher Krzanowski correlation values indicate greater alignment of the leading PC subspaces between **P** and **D**. Similarity estimates are shown for long-snouted species (top) and short-snouted species (bottom). Columns indicate size correction performed with head length included (+SnL, including snout) or excluded (-SnL, excluding snout). The dashed line at 0.75 is included as a visual reference (arbitrary threshold); most long-snouted spikefishes exceed this value, whereas most short-snouted species do not.

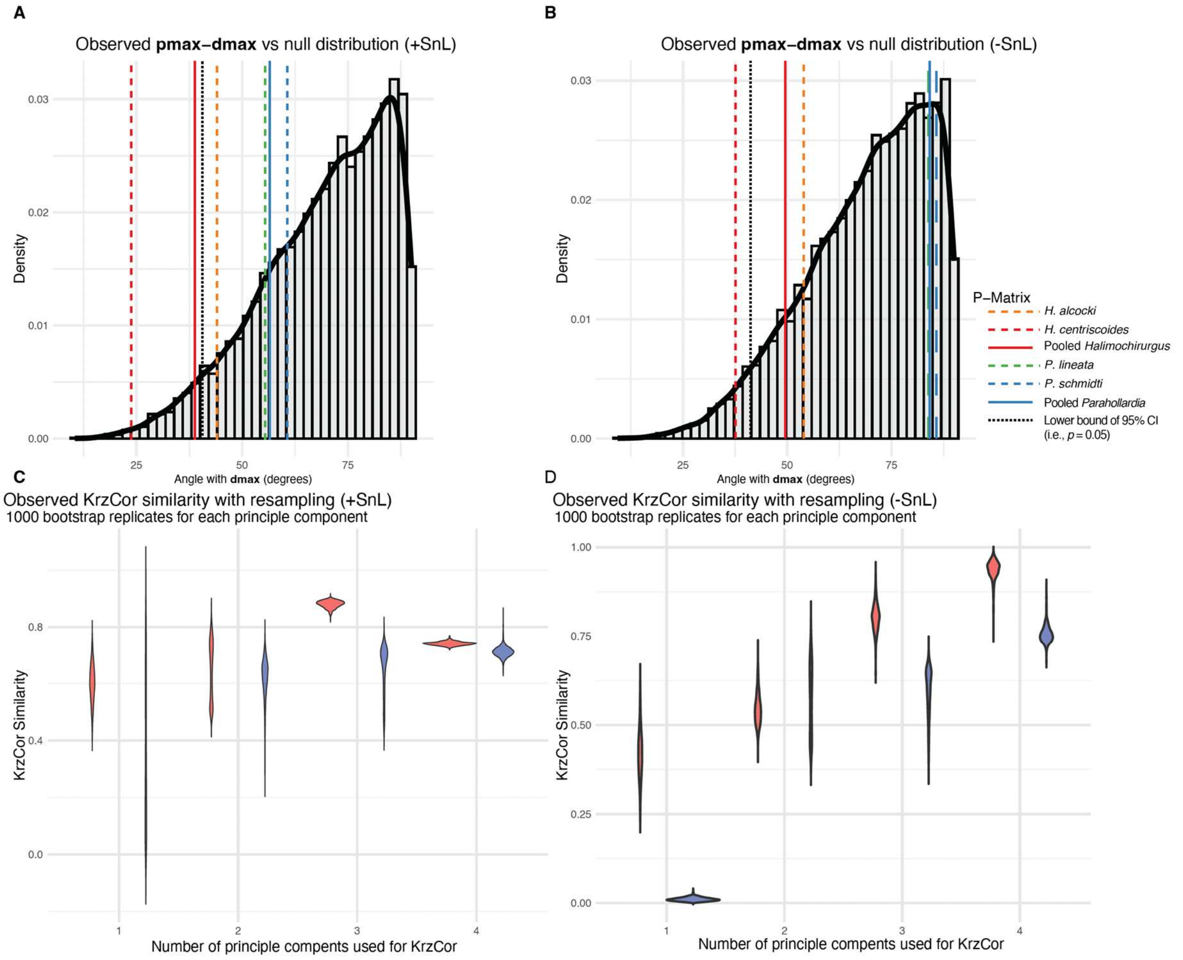


**Figure S11. Alignment of within-species P matrices with the clade-wide divergence matrix (D) under a phylogenetic null. A–B.** Observed vector alignment (degrees) between dmax and pooled- and species-specific pmax, compared to a null distribution of 10,000 alignments generated by simulating six-trait datasets under multivariate Brownian motion on the phylogeny with the empirically estimated rate matrix (following Simon et al.[39]), recomputing dmax for each simulated dataset. Solid lines show pooled **P** matrices (red = long-snouted *Halimochirurgus*; blue = short-snouted *Parahollardia*). Dashed colored lines show species-specific **P** matrices (orange = *H. alcocki*; red = *H. centriscoides*; green = *P. lineata*; blue = *P. schmidti*). The dotted black line marks the 5th percentile of the null distribution (one-tailed critical value, p = 0.05); observed angles below this value indicate stronger-than-expected alignment. **C–D.** Krzanowski correlation between pooled genus-level P matrices and D, estimated 1,000 times via species resampling to quantify uncertainty. Krzanowski correlations were calculated for PC1–PC4 subspaces of each resampled pooled P matrix relative to **D** (red = *Halimochirurgus*; blue = *Parahollardia*). Note, on average, long-snouted matrices and vectors are always more similar to **D** than short-snouted counterparts.

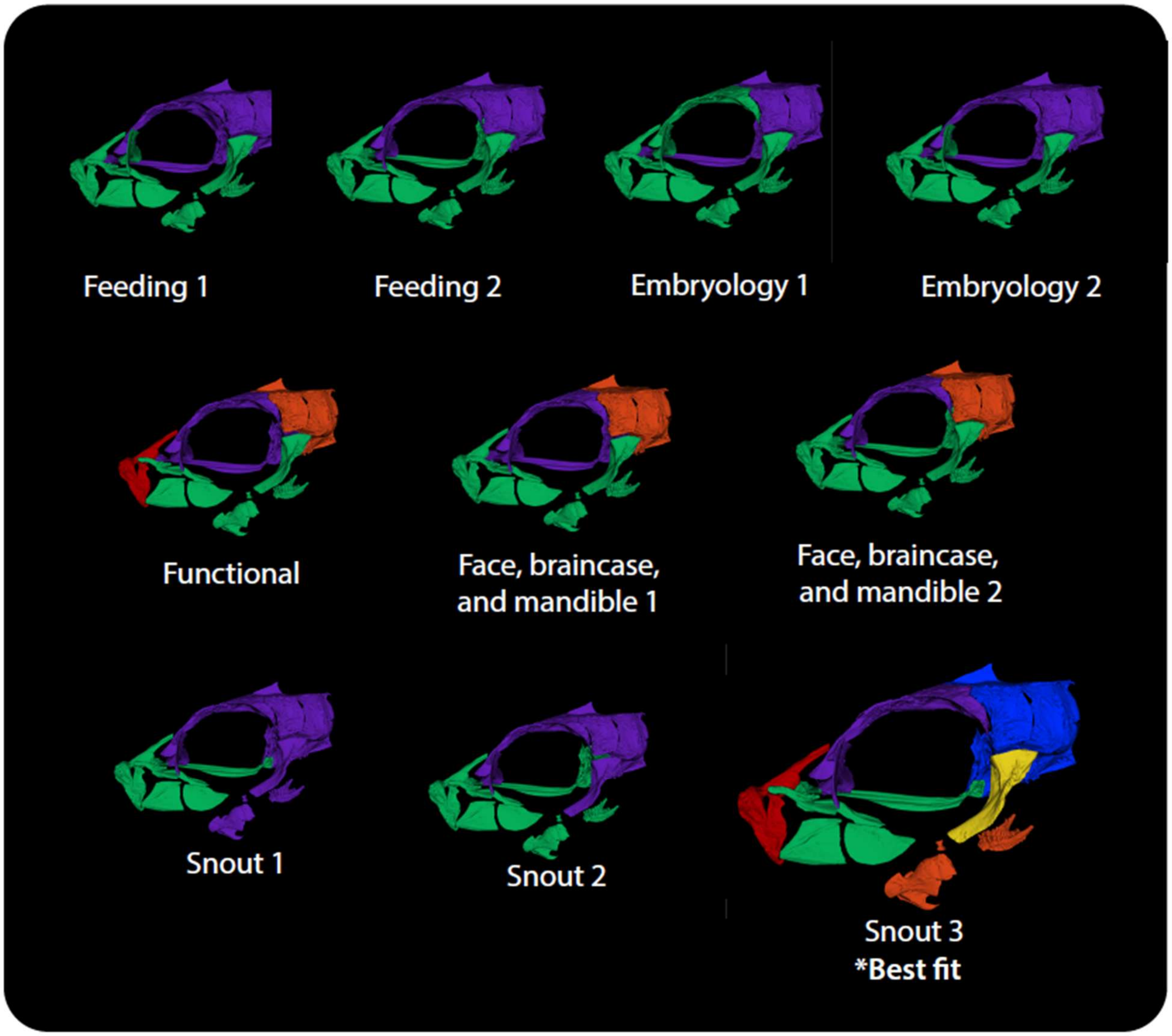


**Figure S12. Testing alternative modularity hypotheses in spikefishes cranium**. Alternative modularity hypotheses evaluated to identify the strongest modularity signature in the spikefish cranium. Modular hypotheses 1–7 are adapted from Troyer et al.[22], while hypotheses 8–10 (Snout 1–3) are newly proposed in this study. Skull shown is *Paratriacanthodes retrospinis*.

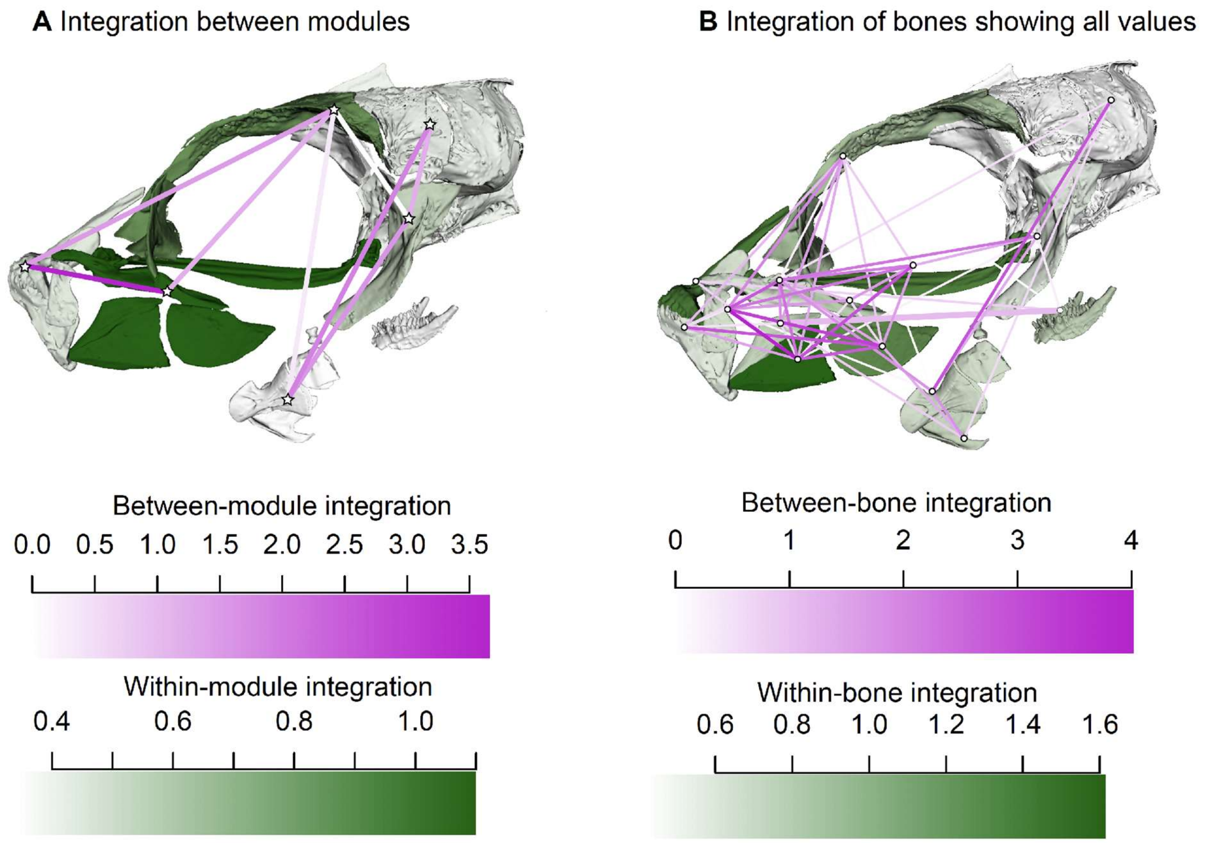


**Figure S13. Extended integration networks of spikefishes cranial modules and bones.** (A) Network plots of significantly integrated modules (post-FDR correction). Darker pink lines indicate stronger integration, with stars marking distinct modules. Bone color reflects the strength of within-module integration (Vrel), as shown in the color gradient. (B) Network plots of all significantly integrated bones using the same color scheme as in (A). Unlike Figure 2G, which shows only the top third of strongest integrations, this panel includes all significant integrations. Skull shown is *Paratriacanthodes retrospinis*.

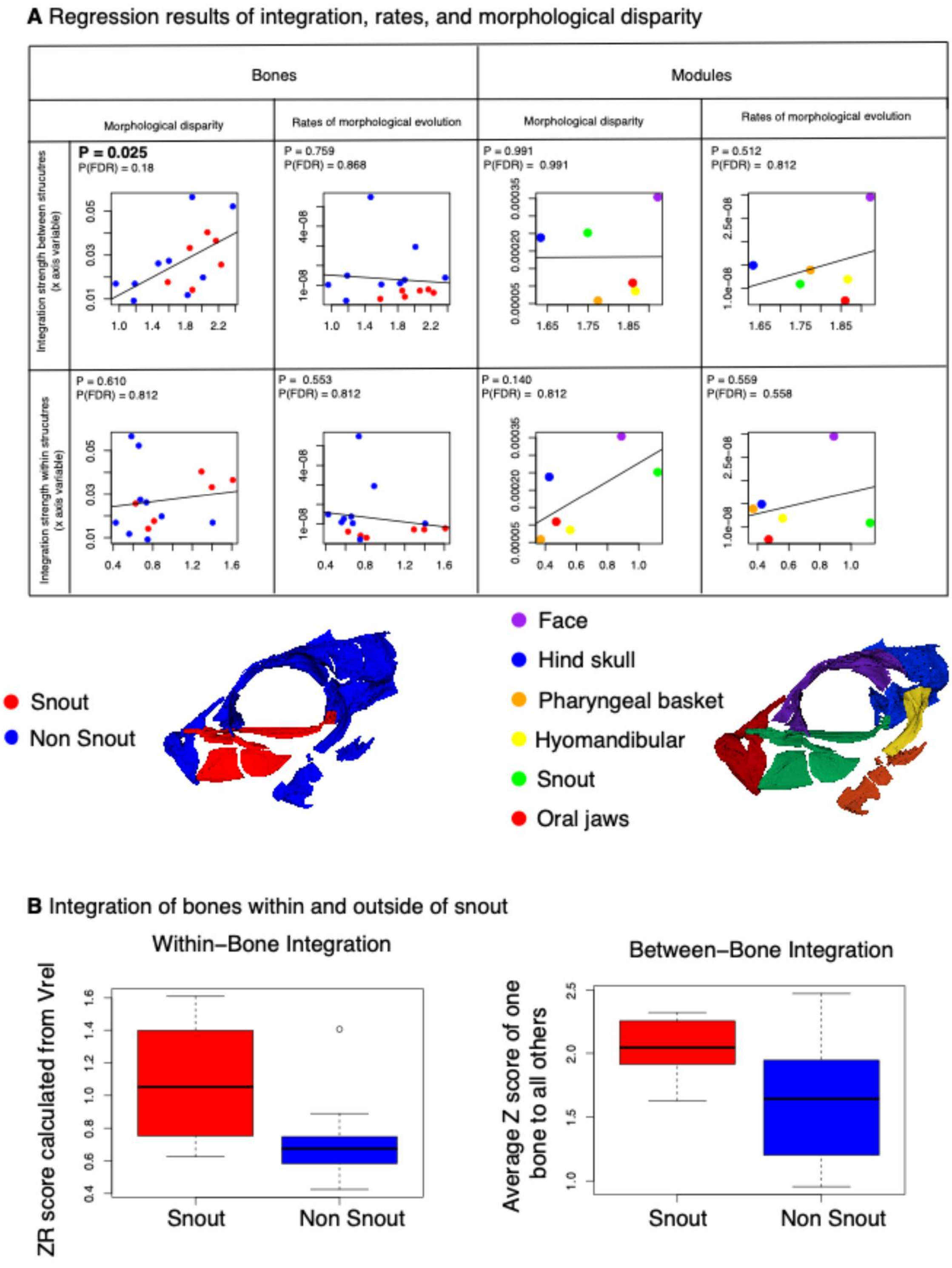


**Figure S14. Integration, evolutionary rates, and morphological disparity in spikefishes cranial structures.** (A) Regression matrix comparing integration strength (within and between bones), rates of evolution, and morphological disparity. Bones are colored snout and non-snout or by module affiliation: snout (red), non-snout (blue). Modules are color-coded: face (purple), hind skull (blue), pharyngeal basket (orange), hyomandibular (yellow), snout (green), oral jaws (red). P-values reported pre- and post-FDR correction, with significant correlations bolded.(B) Box plots of within-bone integration partitioned by snout and non-snout bones. (C) Same as B but using the average integration strength from one bone to all others. Any comparison including a snout bone was categorized as 'Snout'. Skull shown are *Paratriacanthodes retrospinis*.

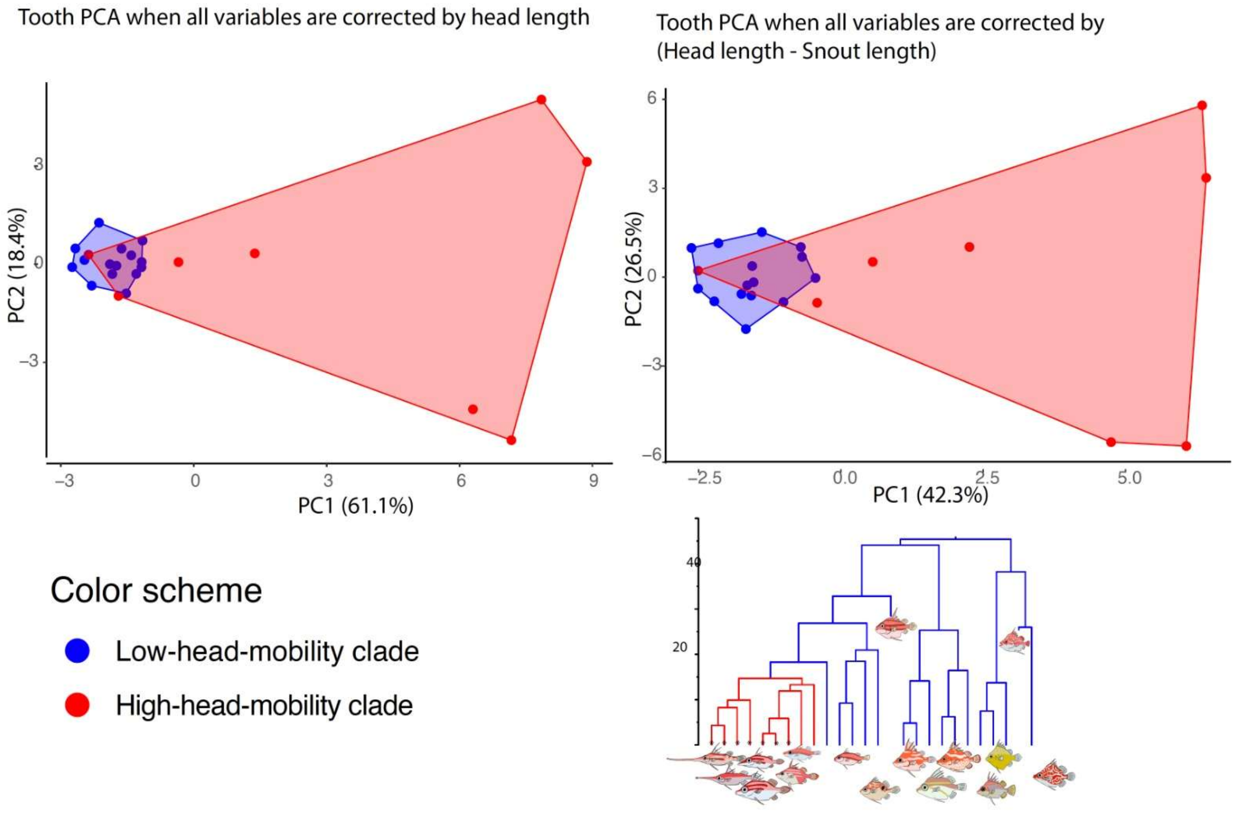


**Figure S15. Principle component analyses of spikefish dental diversity under two alternative frameworks.** (A) Conventional PCA of the tooth dataset, with all variables corrected by head length. (B) Alternative PCA correcting for the effects of snout length, using variables corrected by head length - snout length, and excluding the snout length variable.

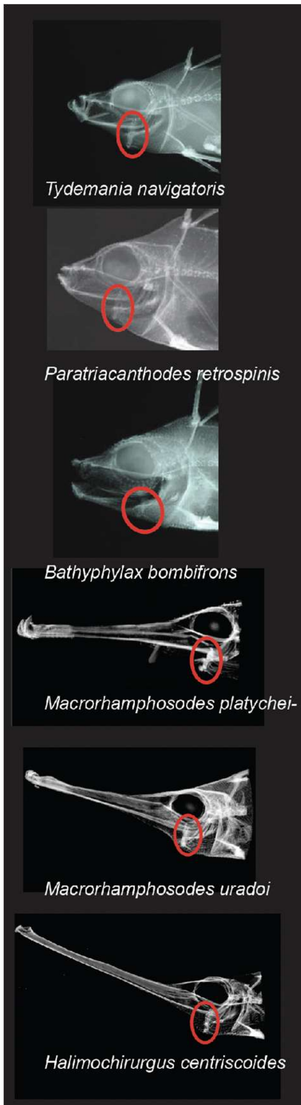


| Species | With hyoid rotation | Total specimen with skeletal prep* |
|---|---|---|
| Tripodfish | 0 | 2 |
| *P. schmidti* | 0 | 12* |
| *P. lineata* | 0 | 13* |
| *H. meadi* | 0 | 2 |
| *H. hollardi* | 0 | 4* |
| *T. ethiops* | 0 | 4 |
| *T. indicus* | 0 | 1 |
| *T. anomalus* | 0 | 3 |
| *T. intermedius* | 0 | 1 |
| *P. herrei* | 0 | 3* |
| *J. eriomma* | 0 | 12* |
| *P. abei* | 0 | 1 |
| *M. fraserbrunneri* | 0 | 2* |
| *M. albomaculosus* | 0 | 1 |
| *H. goslinei* | 0 | 3* |
| *A. japonicus* | 0 | 20* |
| *M. platycheilus* | 5 | 8* |
| *M. uradoi* | 3 | 4 |
| *H. alcocki* | 2 | 12* |
| *H. centriscoides* | 4 | 10* |
| *T. navigatoris* | 1 | 9* |
| *P. retrospinis* | 1 | 9* |
| *B. omen* | 0 | 1 |
| *B. pruvosti* | 1 | 4* |
| *B. bombifrons* | 1 | 4* |

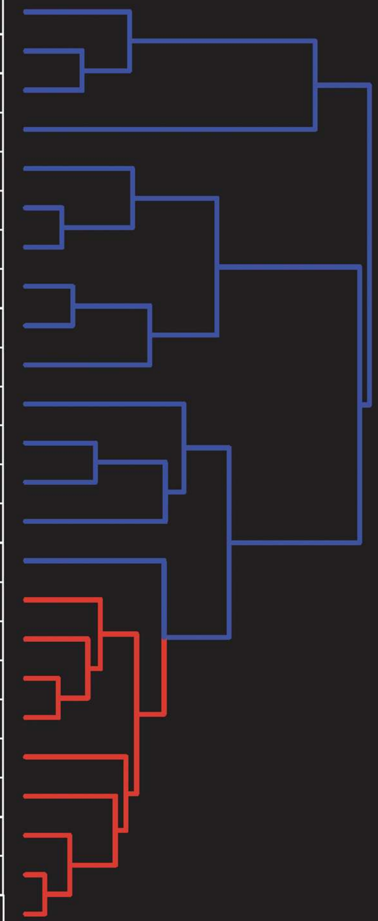

*Includes X-ray photographs from either literature, USNM database, or ASIZP X-ray

**Figure S16. Species, specimens, and phylogenetic distribution of cranial elevation coupled buccal expansion adaptations in spikefishes.** (A) X-ray and CT-scan evidence of specimens preserved with the hyoid rotated ventrally and the head tilted dorsally. Red circles depict the ventrally rotated hyoid. (B) Tabulated counts of specimens preserved with hyoid rotation by individual species. Values in each cell are calculated from a data set of CT scans and expanded by images and X-ray data from the USNM, SIO, and ASIZP (expanded counts marked with asterisks). Species ordered by phylogeny (right). Red values indicate the clade where hyoid rotation is observed.

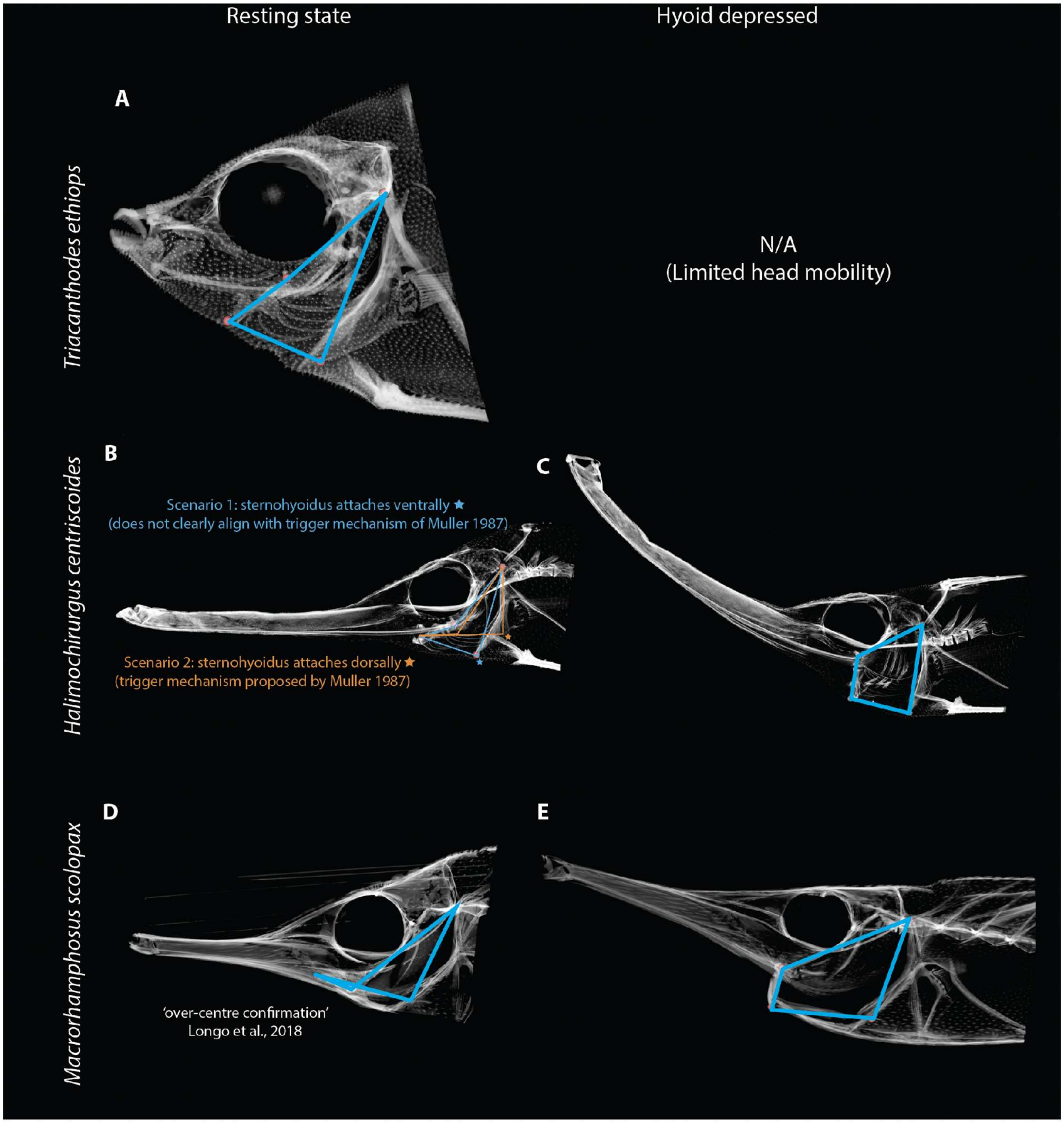


**Figure S17. Comparisons of four bar linkage systems in snipe- and spikefishes.** The predicted four-bar linkage (blue) system for hyoid-coupled head-elevation in closed and opened positions. (A) Short-snouted spikefish, *Triacanthodes ethiops,* (B) long-snouted spikefish, *Halimochirurgus centriscoides* in the closed position. Orange four-bar linkage depicted in *H. centriscoides* aligns with Muller[69] 'trigger mechanism' if the sternohyoidus muscle attaches at the broadest, striated portion of the cleithrum. (C) The four bar linkage shape when the head is elevated in *H. centriscoides*. (D) The snipefish, *Macrorhamphosus scolopax*, (Syngnathiformes) in the 'over-centre' documented in Longo et al.[61] and (E) the head elevated position.

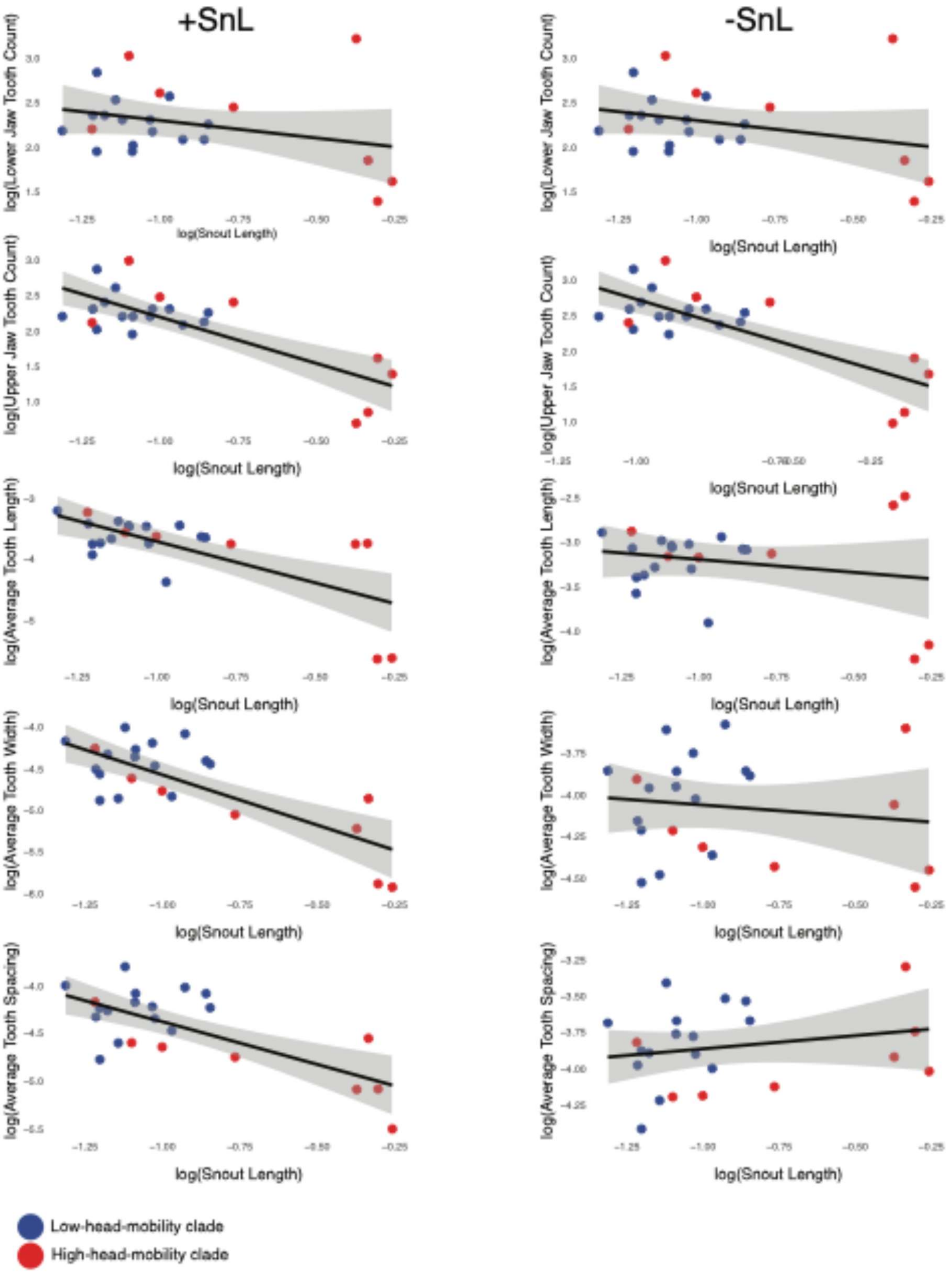


**Figure S18. Linear regressions of tooth traits with snout length.** Regression of tooth traits with snout length. Blue dots correspond to the low-head-mobility clade; red corresponds to high-head-mobility species. Columns indicate size-correction performed with head length included (+SnL, including snout) or excluded (-SnL, excluding snout).

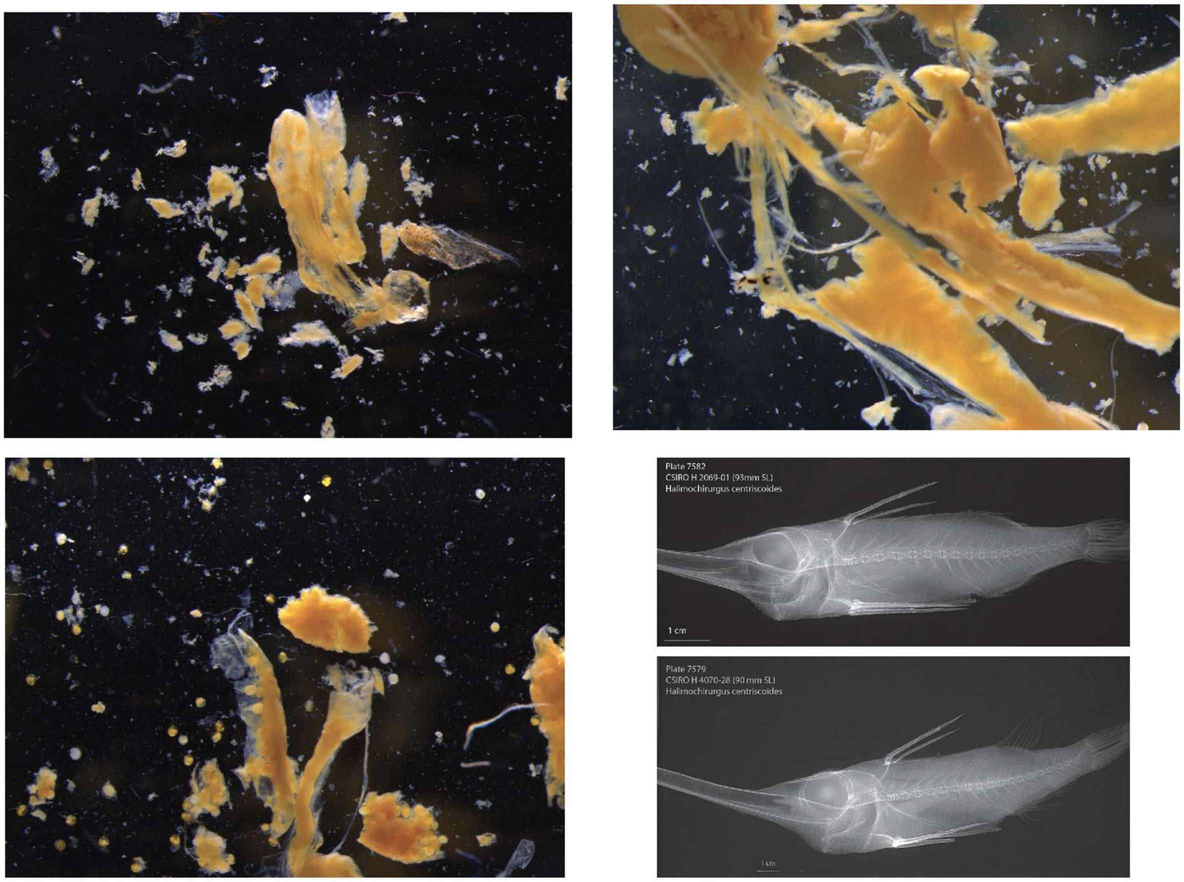


**Figure S19. Representative stomach content and x-rays of *Halimochirurgus*.** Images of stomach contents from a *Halimochirurgus* stomach showing indistinguishable food items. Small spheres are suspected eggs of a gravid spikefish–not gut contents. X-ray images show empty stomachs.

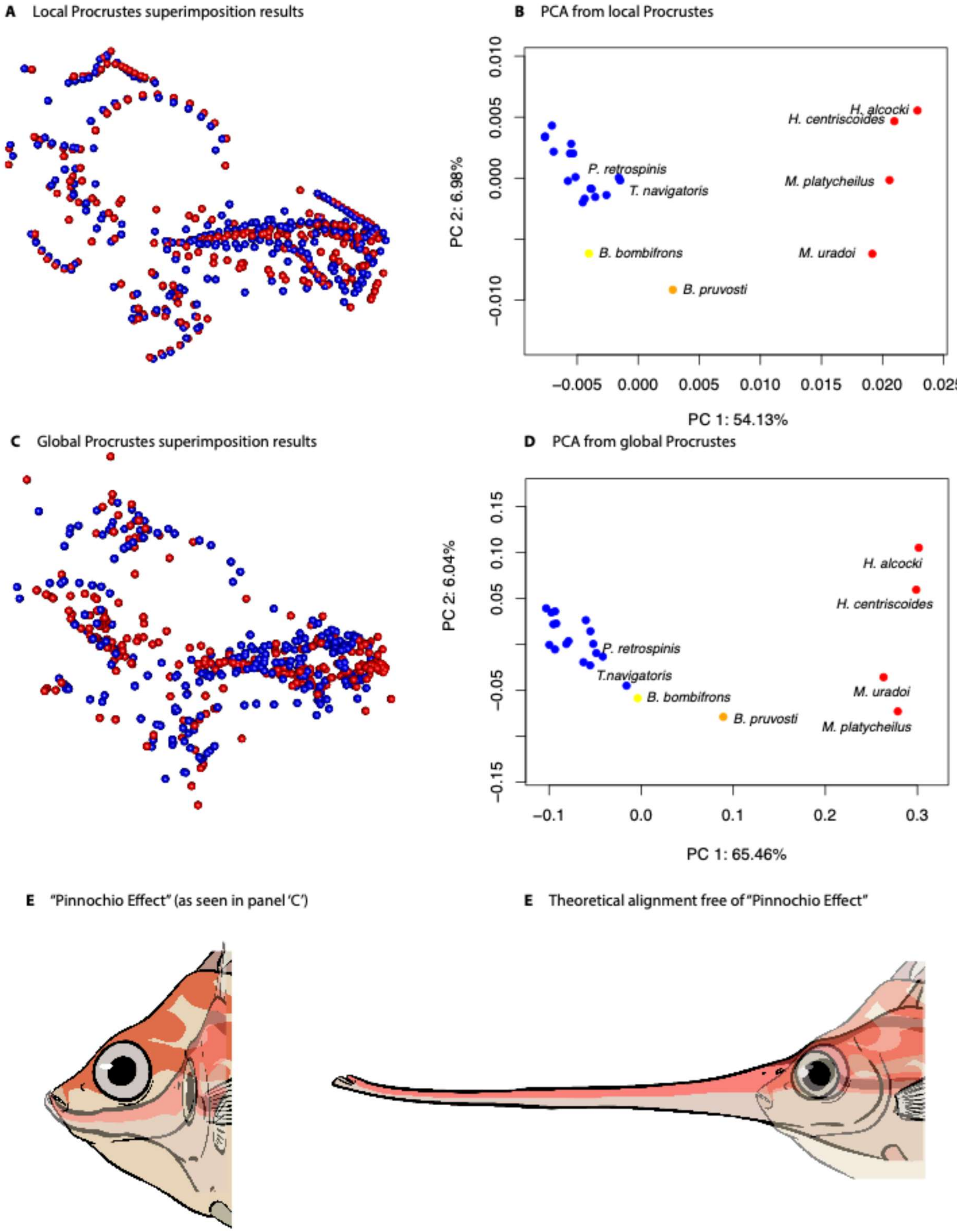


**Figure S20. Effects of local and global superimposition on shape analysis and the 'Pinocchio effect' in spikefishes.** (A) Coordinate alignments of landmarks for each bone, superimposed separately and mapped to a reference (left). Blue and red points represent a short-

and long-snout species, respectively. This local alignment scheme isolates shape by removing position and size information. (B) Principal coordinate analysis of shape data derived from the local alignment in A. (C) Coordinate alignments using a global superimposition scheme (all landmarks aligned simultaneously) illustrating the ‘Pinocchio effect’ where landmark distances are minimized to maximize shaped distortion. Alignment scheme maintains some positional information but compromises biological accuracy; see panel E. (D) Principal coordinate analysis estimated from shape alignment in C.  (E) Qualitative example illustrating misalignment of outlier taxa during global superimposition; as shown in C. (F) Qualitative example of a theoretical shape alignment unaffected by the ‘Pinocchio effect,’ demonstrating an unbiased alignment approach.

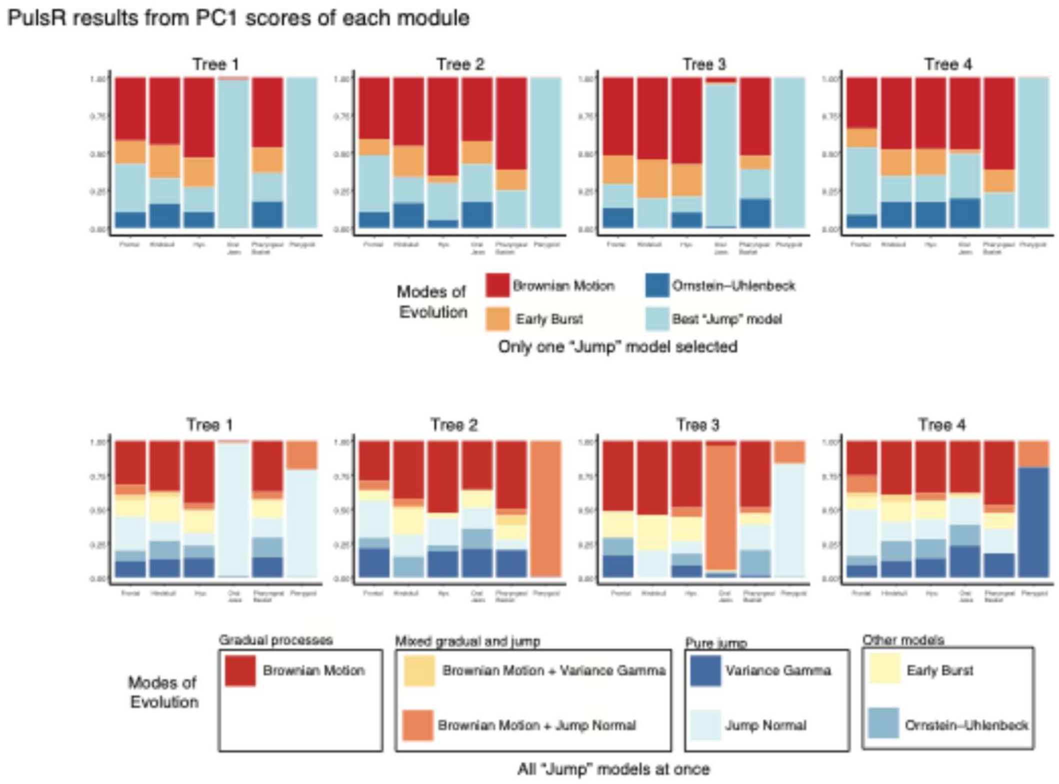


**Figure S21. Robustness of PulsR model fitting across tree topologies conducted across six cranial module PC1 scores:** PulsR model fitting results for the primary axis of variation from PCAs of shape data across six cranial modules. Top panels compare Brownian Motion, Early Burst, and Ornstein-Uhlenbeck models with the best of four ‘jump’ models tested. Bottom panels show initial model fitting with all jump models compared simultaneously. Jump models included Jump Normal, Jump Normal + Brownian Motion, Variance Gamma, and Variance Gamma + Brownian motion.

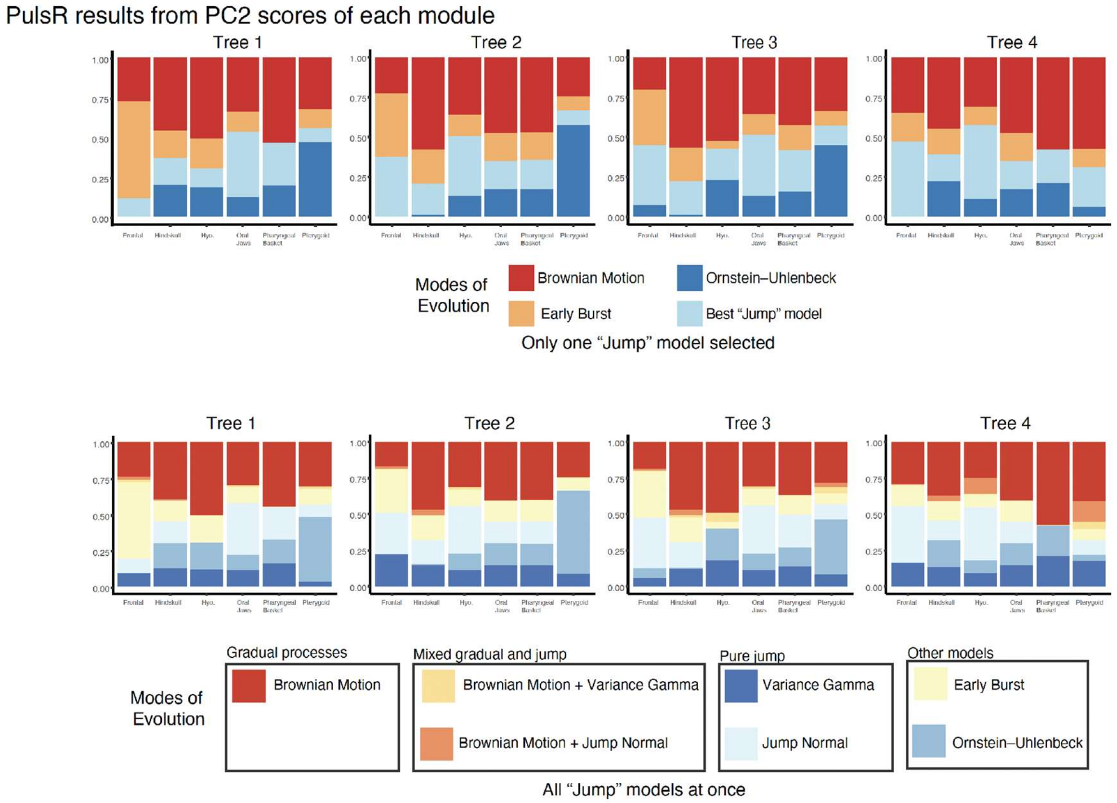


**Figure S22. Robustness of PulsR model fitting across tree topologies conducted across six cranial module PC2 scores:** PulsR model fitting results for the second axis of variation from PCAs of shape data across six cranial modules. Top panels compare Brownian Motion, Early Burst, and Ornstein-Uhlenbeck models with the best of four 'jump' models tested. Bottom panels show initial model fitting with all jump models compared simultaneously. Jump models included Jump Normal, Jump Normal + Brownian Motion, Variance Gamma, and Variance Gamma + Brownian motion.

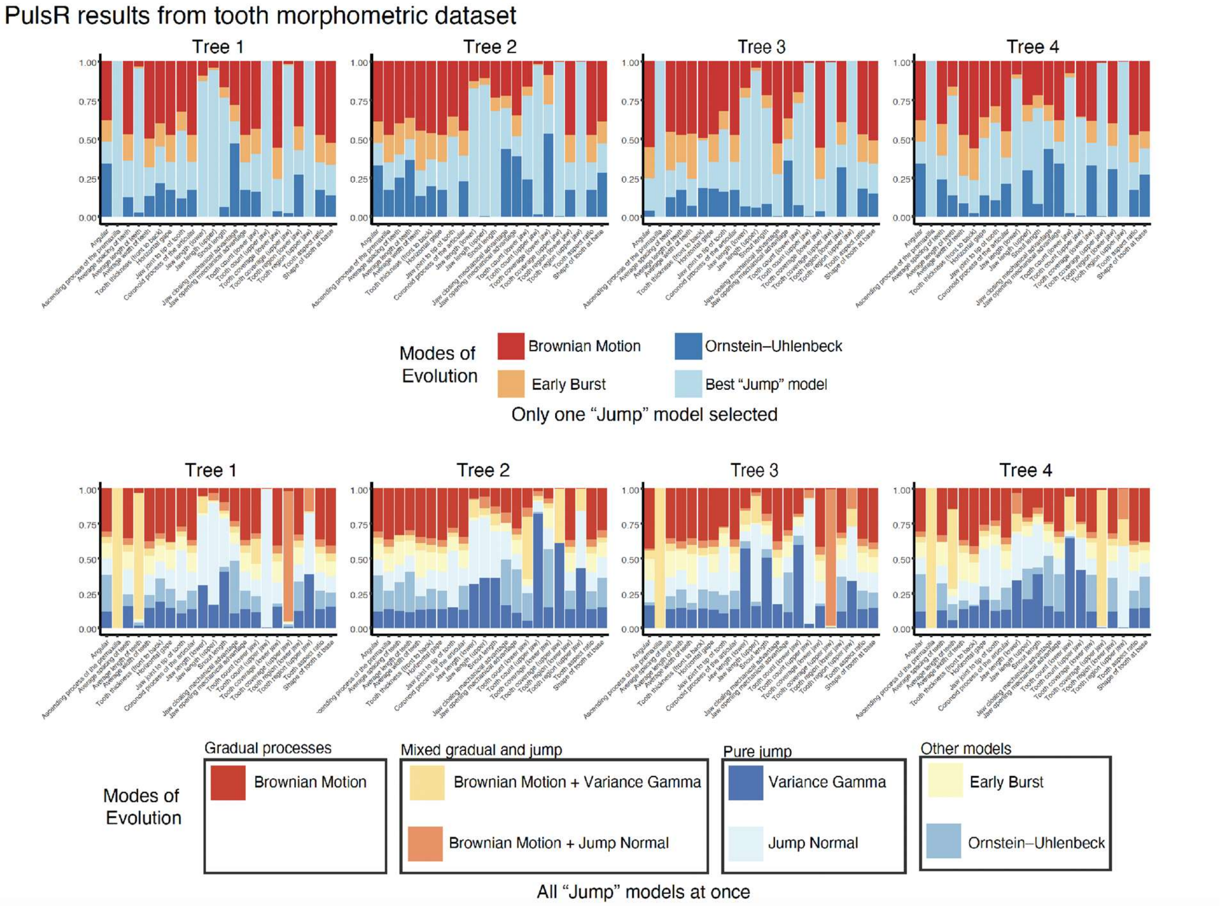


**Figure S23. Robustness of PulsR model fitting across tree topologies conducted across tooth morphometric measurements:** PulsR model fitting results from size-corrected linear measurements of teeth. Top panels compare Brownian Motion, Early Burst, and Ornstein-Uhlenbeck models with the best of four 'jump' models tested. Bottom panels show initial model fitting with all jump models compared simultaneously. Jump models included Jump Normal, Jump Normal + Brownian Motion, Variance Gamma, and Variance Gamma + Brownian motion.

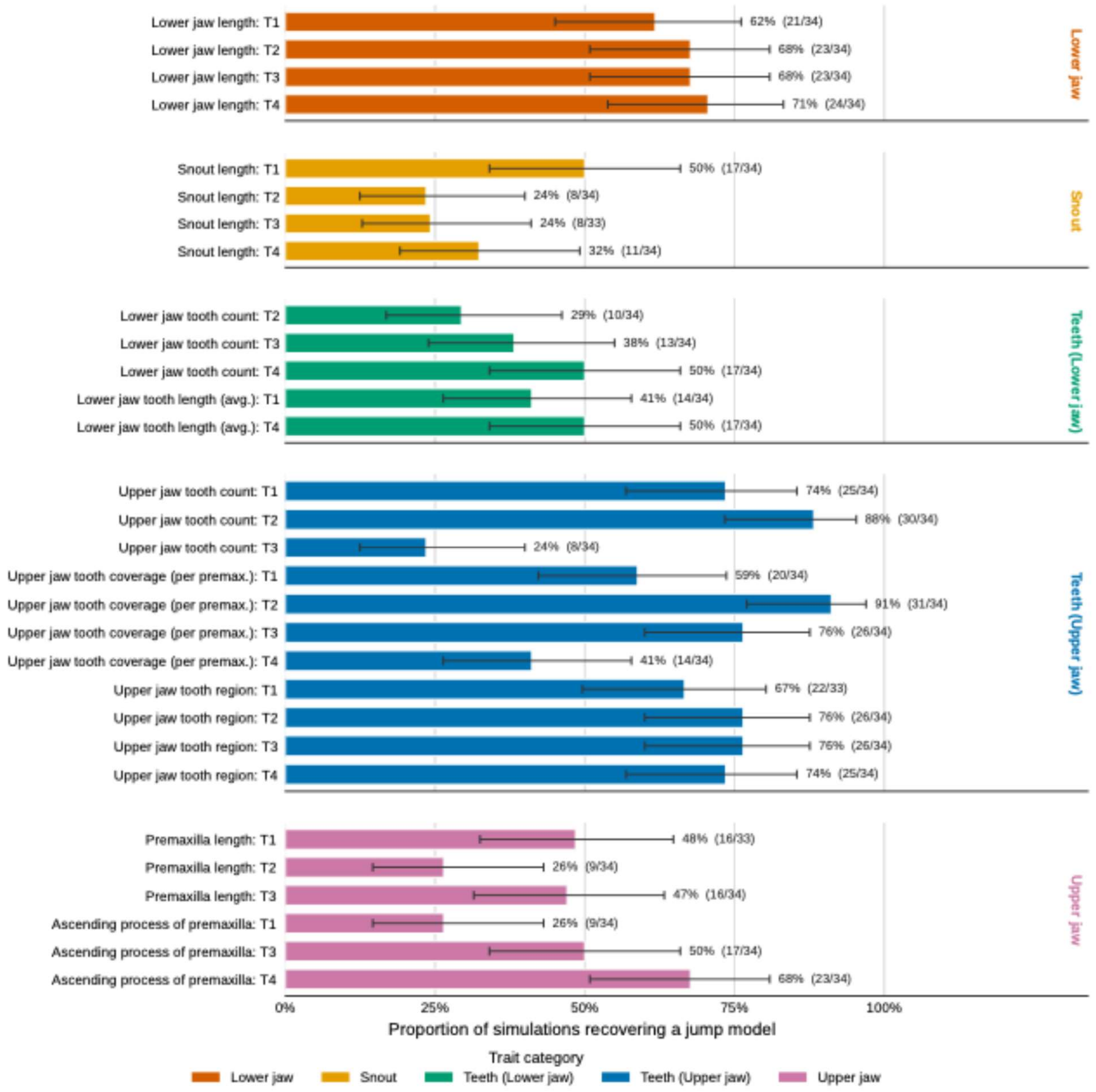


**Figure S24. PulsR sensitivity analyses using empirical parameter estimates.** For each trait supported by a jump model in the empirical PulsR runs, data were simulated under the fitted model on the corresponding tree and re-analyzed to assess how often a jump model was recovered (recovered/total in parentheses). Bars show the proportion of simulations recovering any jump model; error bars are 95% confidence intervals. Three of 1,020 simulations failed to converge and were excluded (n = 33 for the affected trait–tree combinations; n = 34 otherwise). Each bar is an individual trait on one tree, grouped into five categories (color) and, within each category, by tree (T1–T4, estimated with ASTRAL and IQ-TREE); traits not supported on all four trees have fewer than four bars. Trait categories are color-coded as lower jaw (dark orange), snout (gold), lower jaw teeth (green), upper jaw teeth (blue), and upper jaw (reddish purple). Recovery was trait- and tree-dependent: lower-jaw length (62–71%) and upper-jaw tooth region (67–76%) were recovered consistently, other tooth and premaxilla traits varied across trees (24–

91%), and snout length showed the lowest recovery (24–50% as any jump model; 9–21% as the exact generating model), indicating limited power for these parameter estimates at n = 24.

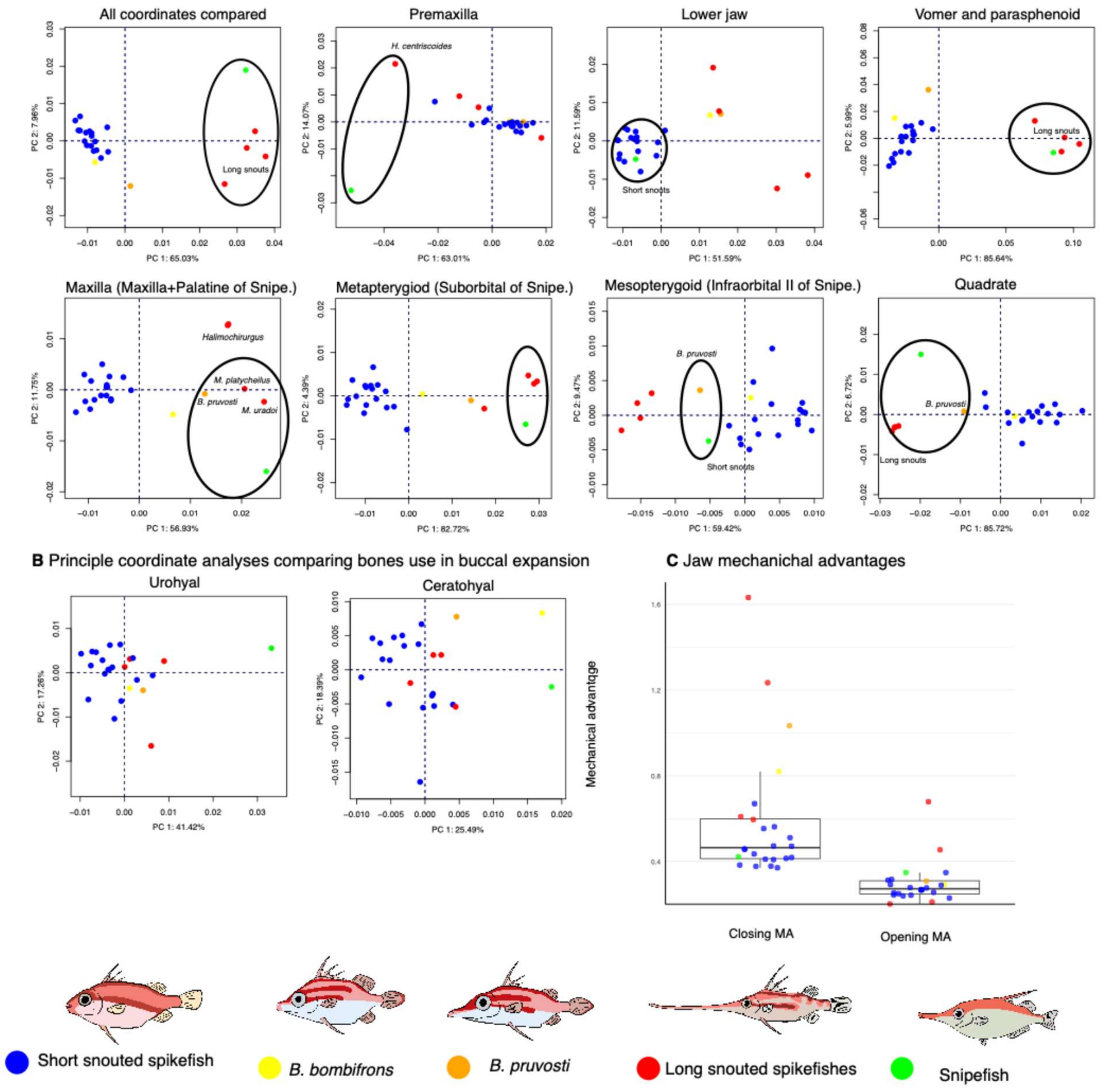


**Figure S25. Morphological comparisons of spikefishes and snipefish bones.** (A) Principal coordinate analyses (PCoA) of snout bones in spikefishes compared to *Macroramphosus scolopax.* Parenthetical PCoA titles indicate comparisons of distinct bones between spikefish and spikefish species based on positional homology. Black circles illustrate clustering of the snipefish to spikefishes (B) As above but comparing bones involved in buccal expansion. (C) Average jaw opening and closing mechanical advantages—calculated as ratio of jaw lever (the coronoid process) to the width-corrected lower jaw length (see Heiple et al. 2021) for 23 spikefish species and *M. scolopax.* (A–C) Colors of data points are as follows: short snout spikefishes=blue, *B. bombifrons*=yellow, *B. pruvosti*=orange, long snout spikefishes=red, snipefish=green.

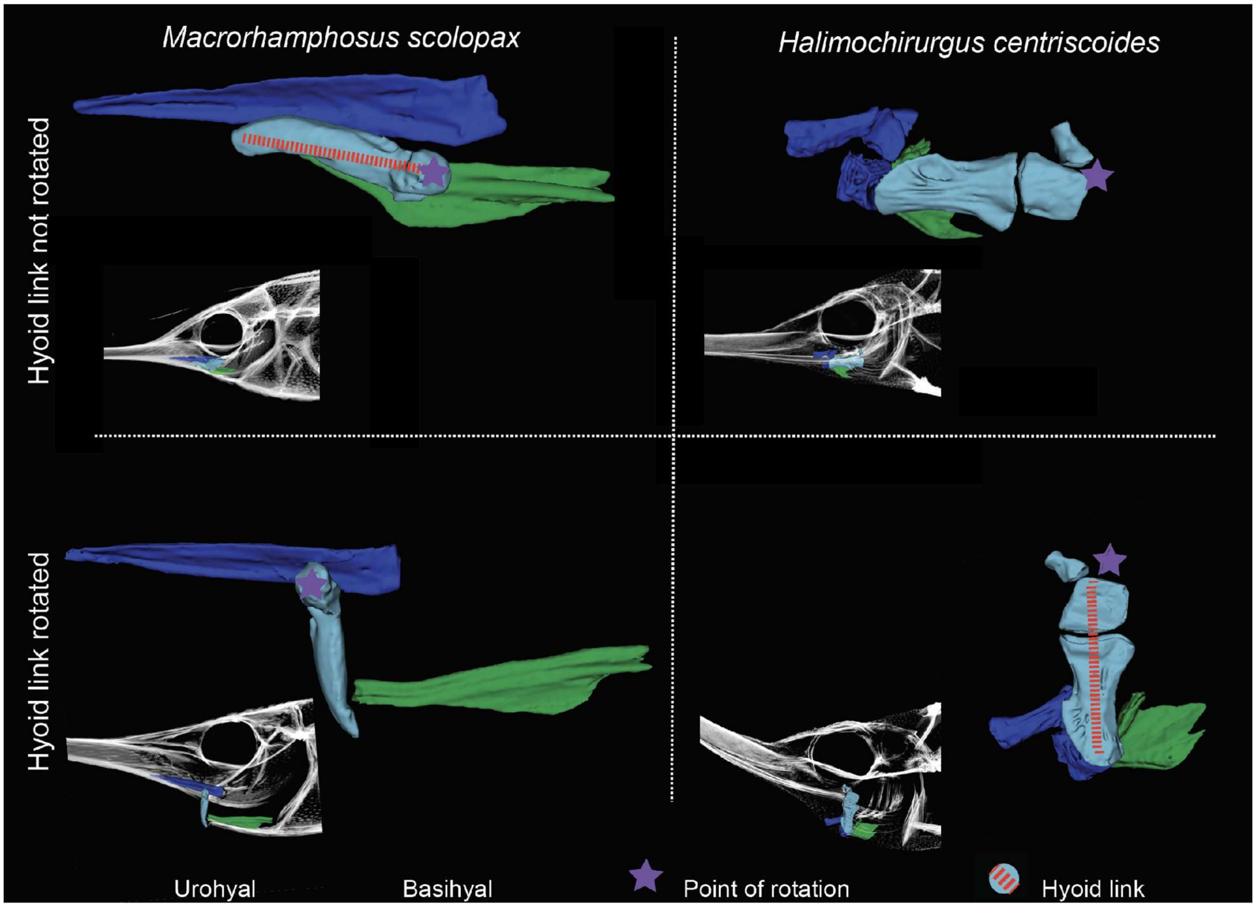


**Figure S26. Cranial elevation coupled buccal expansion mechanism of snipe- and spikefishes is functionally similar but morphologically distinct.** Comparative anatomy of the pharyngeal basket in the snipefish *Macrorhamphosus scolopax*[61] (left) and the spikefish *Halimochirurgus centriscoides* (right). The urohyal is shown in green, the basiohyal in blue, and the hyoid link in teal with red dashed outlines, indicating its composition from multiple bones in spikefishes. Note the differing points of rotation (purple stars) and the relative position of the basiohyal in relation to the hyoid depression.

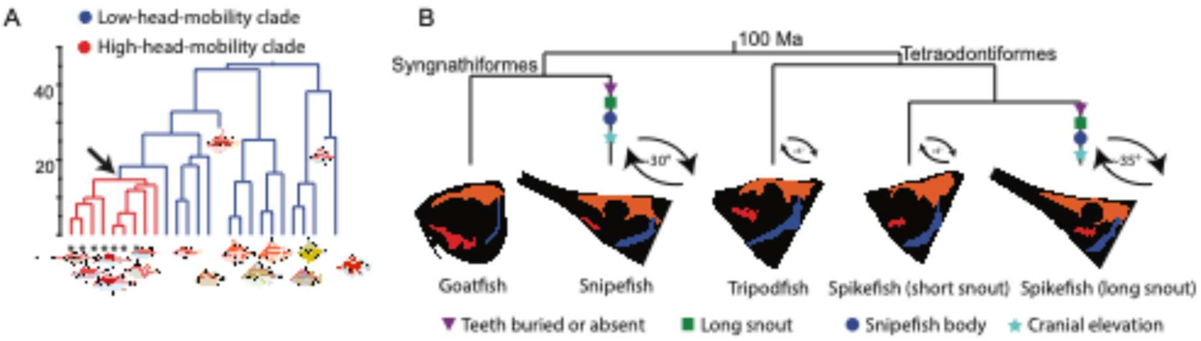


**Figure S27. Spikefish head mobility adaptations and convergent evolution. (**A) Phylogeny of spikefishes highlighting a subclade with adaptations for buccal expansion and dorsoventral head rotation (red branches); taxa lacking these features are shown in blue. Terminals marked with an asterisk indicate species with elongated snouts. **(**B) phylogeny depicting relationships among spikefishes and snipefishes, and their close relatives (goatfish and tripodfish). Traits mapped onto the tree illustrate convergent features shared between spikefishes and snipefishes, with shading corresponding to the neurocranium (orange), its articulation with the cleithrum/post temporal bones (blue), and the hyoid bar (red).

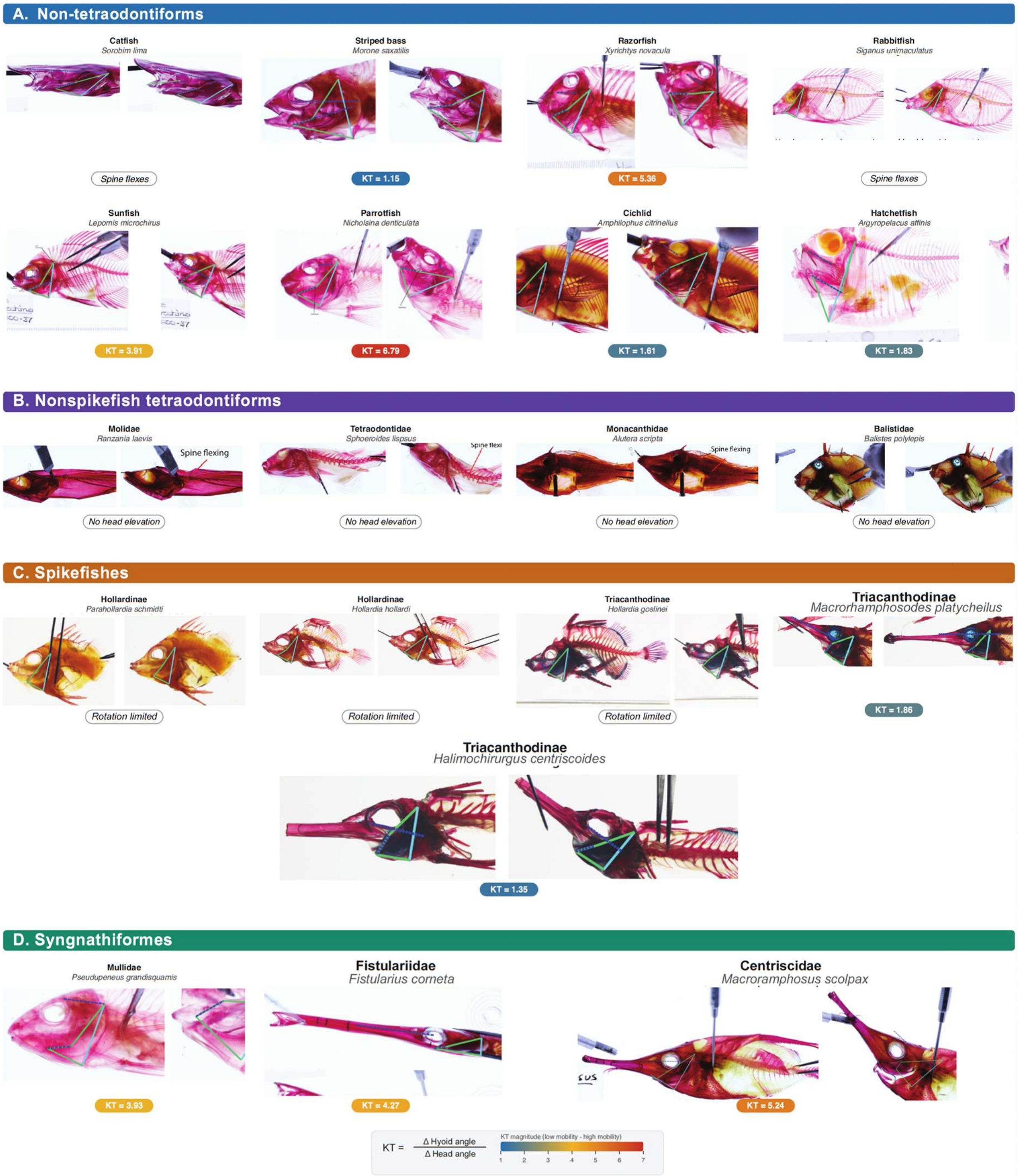


**Figure S28. Head elevation and kinematic transmission of the four-bar linkage coupling head elevation and buccal expansion in tetraodontiform fishes, compared with syngnathiforms and other teleosts**. Kinematic transmission (KT), the ratio of hyoid rotation to head rotation (KT = Δhyoid angle / Δhead angle), was estimated from cleared-and-stained specimens. For each taxon, the two images show head position before head elevation (left) and during head elevation (right). The four-bar linkage comprises a fixed link (solid light-blue line) between the neurocranial head joint and the ventral cleithrum, and three mobile links (green lines) connecting the neurocranial joint to the interhyal, the interhyal to the symphysis of the ceratohyals, and that symphysis to the ventral cleithrum. Blue dashed lines mark the hyoid bar and head position in each state. Linkages are not drawn, and KT is not reported, where head elevation is absent or highly limited; "spine flexing" denotes taxa in which head movement is achieved through dorsal-spine flexion rather than the four-bar mechanism. KT values are reported below each specimen and colored by magnitude, from low (blue) to high (red) mobility, following the scale at bottom. Comparisons are grouped as (A) non-tetraodontiform teleosts, (B) nonspikefish tetraodontiforms, (C) spikefishes, and (D) syngnathiforms

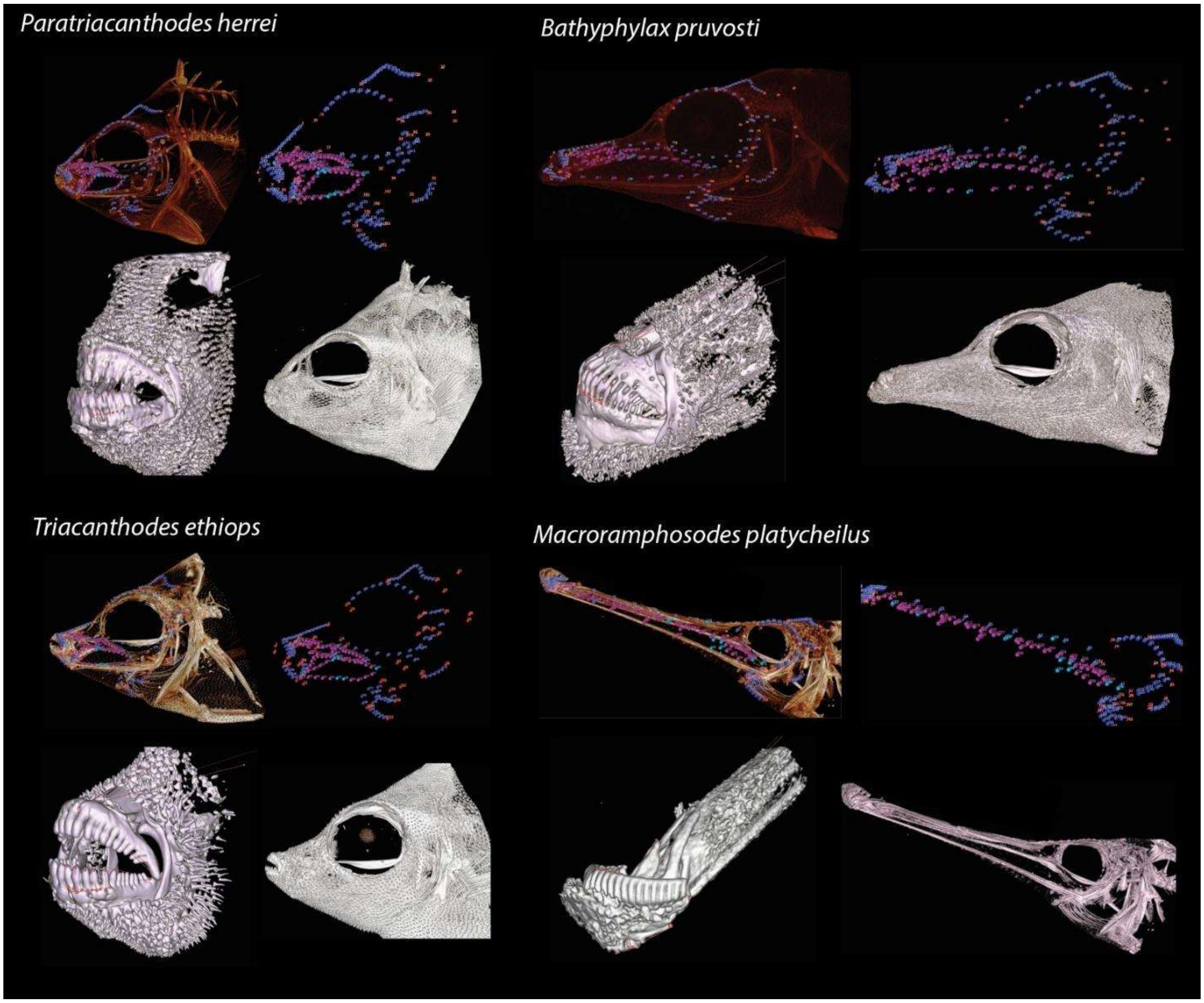


**Figure S29. Morphological data collected for this study.** Landmark and tooth morphometric data collected shown in four species: *Paratriacanthodes herrei, Bathyphylax pruvosti, Triacanthodes ethiops,* and *Macrorhamphosodes platycheilus*. Top panels for each species show the landmarking scheme. The bottom left panels show tooth morphometric measurements. Bottom right panels show the fish with only raw CT data.

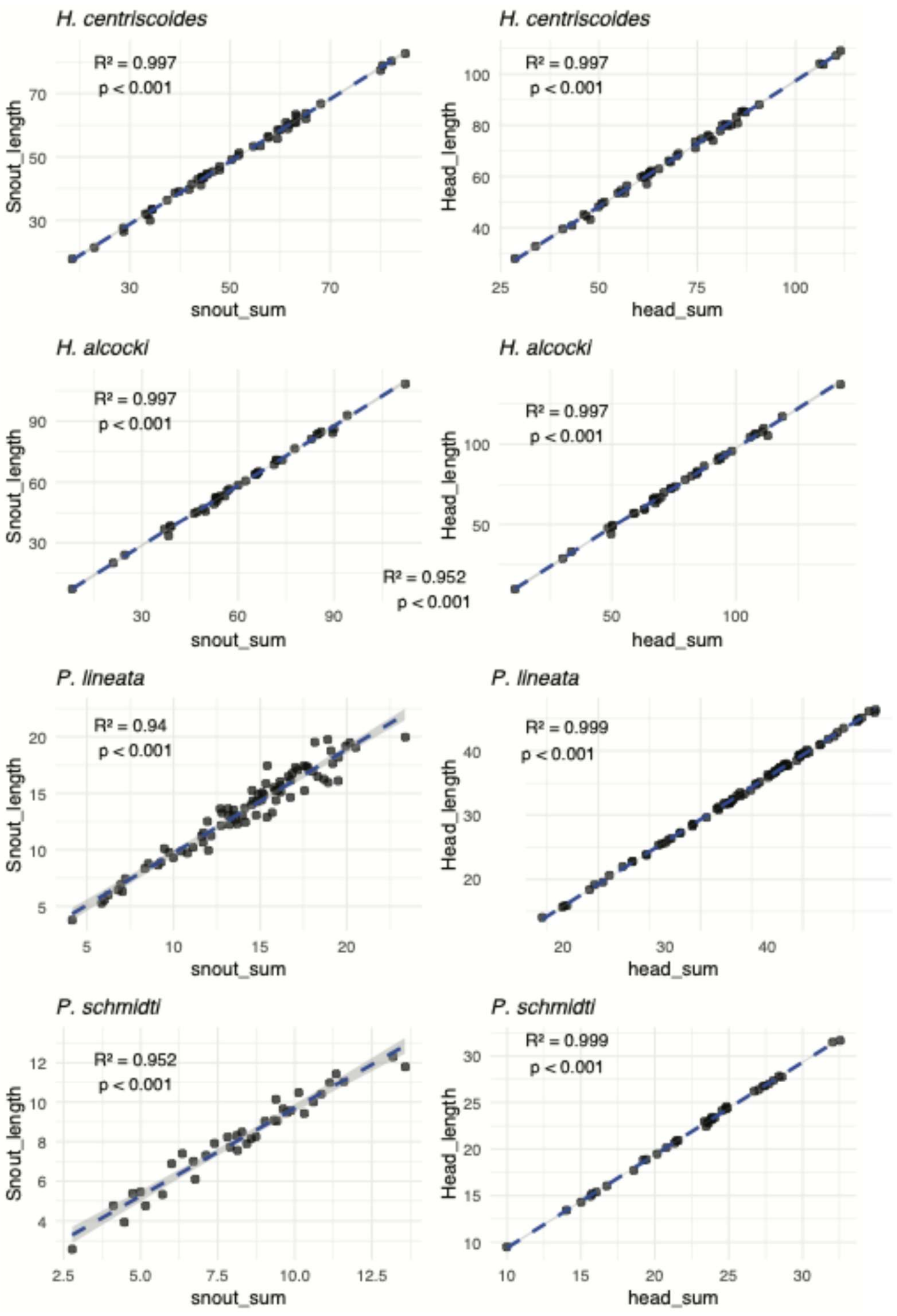


**Figure S30. Modified snout and head length protocol are effective proxies.** Linear regressions of our modified head length (head_sum) and modified snout length (snout_sum) with unmodified measurements.

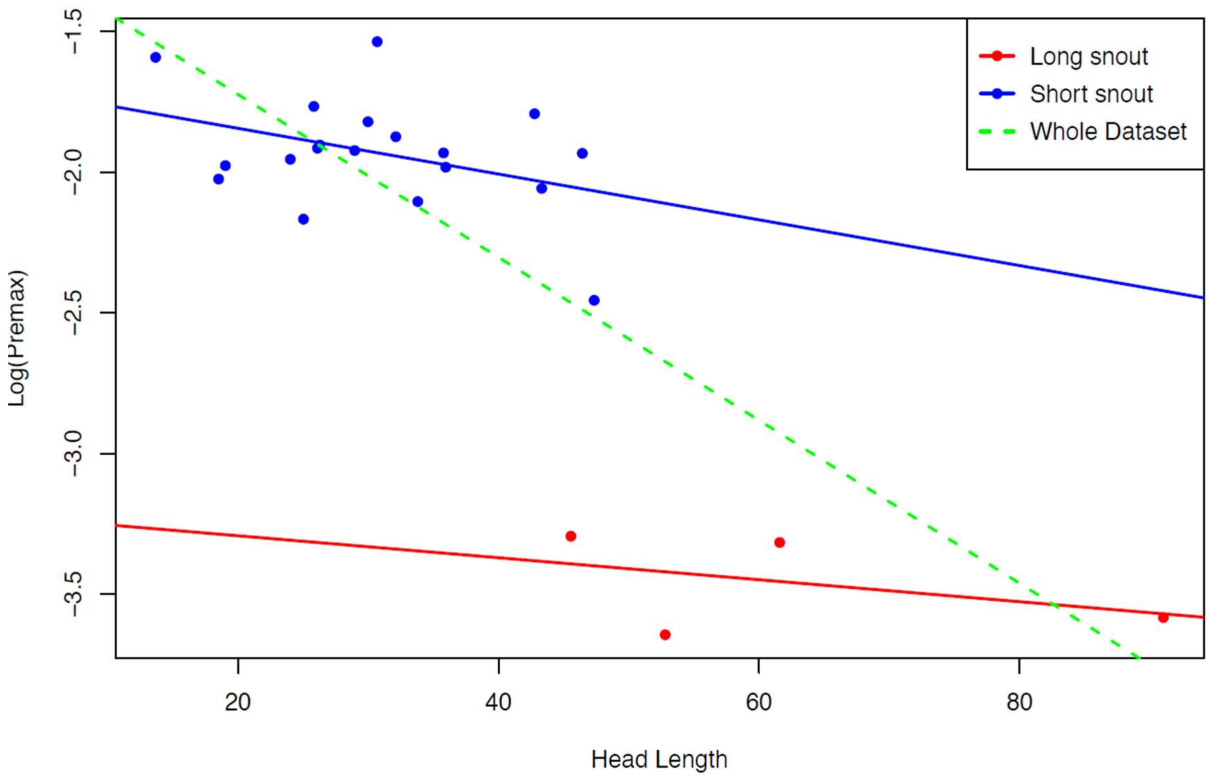


**Figure S31. Allometric regression of premaxilla ascending process and head length in spikefishes.** Regression analysis testing for allometry between the ascending process of the premaxilla (log-transformed) and head length. Data points are colored by snout length: long-snout spikefish (red), short snout spikefish (blue), with trend lines for each group (red and blue, respectively) and for all species combined (green). Long-snout species often appear as outliers, impacting the combined trend line, and making subsequent residuals challenging to interpret.

### Supplementary table captions

**Table S1. Specimens and metadata for phylogenomic analysis.** List of specimens used for phylogenomic tree estimation, including museums, collection voucher numbers, and GGBN (Global Genome Biodiversity Network) IDs. Museum acronyms follow Sabaj (2020)[70] for standardized institutional codes. Sequence names match the input file names, and species names reflect the final nomenclature adopted for this study.

**Table S2. Species with only morphological data in Bayesian tree estimation.** A list of species included in total-evidence tree estimation with either 'Morphological data only' or 'Genomic + Morphological data'. Each species corresponds to the set of individuals used to code the morphological matrix.

**Table S3. Alignments quality control.** A list of the markers used for phylogenomic tree estimation. Reporting names of genes mined using the bioinformatic pipeline detailed in Hughes et al. (2021)[1], the number of individuals captured by a gene. Genes with less than 10 individual per species were excluded, noted ‘Removed because less than 10 species.’

**Table S4. Alignments used in downstream analyses.** Specimen metadata, including species name, file names, number of genes captured per individual, and inclusion in various phylogenetic tree estimation analyses.

**Table S5. Materials examined, morphological matrix.** Specimen metadata for all materials used in curation of the morphological matrix, including species name, voucher numbers, and preservation media. Preservation media include cleared and stained (C&S), ethanol-preserved (EtOH), CT-scanned (CT), or illustrations from Tyler (1968)[27] used to aid character scoring. Museum acronyms follow Sabaj (2020)[70] for standardized institutional codes.

**Table S6. Scored morphological matrix.** Morphological character matrix (80 characters) scored by species. Bracketed values indicate species polymorphism. Dashes denote “not applicable,” and question marks denote uncertainty in scoring. The list of corresponding characters can be found near the top of this document.

**Table S7. Divergence times per scheme per subset.** Divergence-time estimates derived from the best-fit clock model (Table S8) for the most recent common ancestors of Triacanthodidae (Outgroup absent) alone and of Triacanthodidae + Triacanthidae (Outgroup present). For each of the four non-overlapping genomic subsets, the mean age, 95% confidence interval, and standard deviation are reported.

**Table S8. The results from clock fitting using the Stepping-Stone approach implemented in MrBayes.** Log-likelihood scores for the outgroup-present (Triacanthodidae+Triacanthidae) and outgroup-absent (Triacanthodidae only) schemes under the autocorrelated TK02 and uncorrelated IGR clock models, respectively.

**Table S9: Morphological characters consistent with independent snout elongation in *Bathyphylax*.** Subsampled morphological character matrix highlighting the characters that distinguish Bathyphylax and are consistent with an independent, convergently evolved event of snout elongation.

**Table S10. The results of two-block partial least squares tests for all bones in the snout module.** Results of two-block partial least squares (2BPLS) regression, reporting for each comparison the r-PLS statistic (analogous to $R^2$; values close to 1 indicate high integration), the effect size (Z-score; larger values denote stronger integration), and the associated *p*-value. Coordinates used to estimate values were either locally (left) or globally (right) superimposed coordinates.

**Table S11. *Macrorhamphosodes platycheilus* intermediate status.** A reduced morphological character matrix including only species with snout elongation (genera *Bathyphylax, Macrorhamphosodes, and Halimochirurgus*), showing characters that indicate whether *M. platycheilus* more closely resembles long-snouted spikefishes or *Bathyphylax* (the genus with intermediate snout elongation). The condition typical of long-snouted species is highlighted in grey; note that *M. platycheilus* alternates between grey and white cells. Bracketed values indicate species polymorphism. Dashes denote "not applicable," and question marks denote uncertainty in scoring. The list of corresponding characters can be found near the top of this document.

**Table S12. Within and between genera P matrix comparisons.** Within- and between-genus comparisons of **P** matrices. Matrix similarity indices were calculated for pairs of species **P** matrices across size-correction methods. Indices include Random Skewers (RS), Krzanowski correlations estimated using enough PC axes to explain ≥80% of the variance (KrzCor@80), and vector alignments (in degrees), defined as the angle between PC1 of the two matrices.

**Table S13. Alignment of species- and genus-level PC1 with divergence PC1**. Comparisons of PC1 eigenvectors from species or pooled-genus **P** matrices with the divergence matrix, **D**. **pmax** = PC1 of the **P** matrix; **dmax** = PC1 of the divergence matrix. Significance was assessed by comparing the observed alignment to a null distribution of 10,000 six-trait datasets simulated under multivariate Brownian motion on the phylogeny with the empirically estimated rate matrix, following Simon et al.[39].

**Table S14. Modularity fitting results.** Results of module fitting in geomorph, showing for each tested module the covariance ratio (CR; 0–1, with lower values indicating stronger modularity), the p-value for the significance of modular structure, and the effect size (Z-score; more negative values denote a stronger modular signal). Analyses were performed under two Procrustes superimposition schemes: global (all coordinates aligned simultaneously; 'Global') and local (coordinates of each bone aligned separately; 'Local').

**Table S15. The results of integration *compare.ZVrel* for bones.** Integration calculated using eigen dispersion values for landmarks within each bone. Reporting integration for each bone as ZR ratio (top), effect sizes estimated from pairwise comparisons of integration between bones (middle) and the corresponding statistical significance values (bottom) for each comparison. Low ZR values indicate low integration. Large pairwise effect-sizes between bones in the matrix show the magnitude of difference in integration.

**Table S16. The results of integration *compare.ZVrel* for modules.** Integration calculated using eigen dispersion values for landmarks within each module. Reporting integration for each module (top), effect sizes estimated from pairwise comparisons of integration between bones (middle) and the corresponding statistical significance values (bottom) for each comparison. Low ZR values indicate low integration. Large pairwise effect-sizes between modules in the matrix show the magnitude of difference in integration.

**Table S17. Integration network values used to create network graph for bones.** Comparison of between-bone integration across. Reporting average effect sizes of between-bone integration for each group (top), effect sizes estimated from pairwise comparisons of individual bones, and the corresponding statistical significance values (bottom) for each comparison.

**Table S18. Integration network values used to create network graph for modules.** Comparison of between-bone integration across. Reporting average effect sizes of between-module integration for each group (top), effect sizes estimated from pairwise comparisons of individual modules, and the corresponding statistical significance values (bottom) for each comparison.

**Table S19. Dataset for testing relationship of integration, rates of evolution and morphological disparity for each bone.** A dataset including the average integration of one module to all others, the integration within each module reported as a ZR-ratio (a converted Z-score; 0= no integration and positive values indicate integration), morphological disparity, and rates of evolution for each individual bone.

**Table S20. Dataset for testing relationship of integration, rates of evolution and morphological disparity for each module.** A dataset including the average integration of one module to all others, the integration within each module reported as a ZR-ratio (a converted Z-score; 0= no integration and positive values indicate integration), morphological disparity, and rates of evolution for each individual module.

**Table S21. Description of 22 linear measurements for tooth dataset.** Descriptions of the 22 linear measurements collected for the dentition dataset and the names of each variable as they appear in the code.

**Table S22.Tooth morphometric dataset PC1&2 loadings, size corrected by head length.** The eigenvectors of each trait and their respective magnitudes when head length was used to size correct variables.

**Table S23. Tooth morphometric dataset PC1&2 loadings, size corrected by snout-subtracted head length.** The eigenvectors of each trait and their respective magnitudes when snout-subtracted head length was used to size correct variables.

**Table S24. Summary diet table.** Summary of diet data reported in the literature and from stomach-content observations. Species shown in bold have diet information documented from stomach contents and/or published sources. "Hard prey items" are reported as 'yes' if any individual possessed stomach contents that could be visualized in CT or X-rays

**Table S25. PulsR simulations: Simulation results for PulsR**, results are based on data simulated from parameters associated with each “Significant fits from empirical PulsR runs” column and the proportion of times a jump model was recovered.

**Table S26. Raw FishShapes data for Ct analyses:** Metadata and raw body-shape data pre-transformation for Ct.

**Table S27. Ct results across size-correction methods.** Results of Ct analyses and statistical significance reported for three conceptually distinct size-correction approaches: Approach 1, phylogenetically informed linear regression of trait ~ standard length → extract residuals → generate non-phylogenetically corrected morphospace coordinates → estimate convergence; Approach 2, volume-based size correction → generate non-phylogenetically corrected morphospace coordinates → estimate convergence; and Approach 3, volume-based size correction → generate phylogenetically corrected morphospace coordinates → estimate convergence.

**Table S28. Head mobility estimates from cleared and stained specimens:** Head elevation estimates collected for 23 specimens representing 23 species, including specimen metadata.

**Table S29. Materials examined, CT scans.** Specimens CT-scanned for this project, including associated collections metadata, scan resolution, voltage, amperage, scanning facility and location, Morphosource ID (when applicable), and whether each scan was included in the dentition or skull dataset. Museum acronyms follow Sabaj[70] for standardized institutional codes.

**Table S30. Landmark metadata, spikefish only.** Metadata for all landmarks, including non-applicable entries (NULL), the ordering used in both global and local Procrustes superimpositions, modular-hypothesis assignments grouping landmarks into categories for modularity analyses, and other pertinent details. Note that the datasheet can be filtered to reflect modularity test inputs for global and local superimposition schemes.

**Table S31. Materials examined for estimation of P and D matrices**. List of X-ray image data collected to quantify **P** and **D** matrices. Identified by ZWH indicates instances where specimens were re-identified. "Focal taxa" indicates that the species was either *P. lineata, P. schmidti, H. centriscoides*, or *H. alcocki.*

**Table S32. Description of linear measurements for cranial dataset.** Descriptions of eight linear measurements collected from cranial linear measurements dataset.

**Table S33. Materials examined, head rotation.** List of specimens and associated X-ray image data collected to quantify hyoid and head mobility in spikefishes. Images taken from CT scans, USNM X-ray database, or ASIZ-P database.

**Table S34. Landmark metadata, spikefish and snipefish.** Metadata as in Table S30, restricted to the subset of bones used for the snipefish–spikefish comparison.

(27) Tyler, James C. *A Monograph on the Plectognath Fishes of the Superfamily Triacanthoidea*; Monographs of the Acad.of Nat. Sci. Philadelphia; Academy of Natural Sciences of Philadelphia: Philadelphia, PA, 1968.

(28) Tyler, James C. *Osteology, Phylogeny, and Higher Classification of the Fishes of the Order Plectognathi (Tetraodontiformes)*; NOAA Technical Report NMFS Circular; U.S. Department of Commerce: Washington, DC, 1980.

(48) Rolfe, S.; Pieper, S.; Porto, A.; Diamond, K.; Winchester, J.; Shan, S.; Kirveslahti, H.; Boyer, D.; Summers, A.; Maga, A. M. SlicerMorph: An Open and Extensible Platform to

Retrieve, Visualize and Analyse 3D Morphology. *Methods Ecol. Evol.* **2021**, *12* (10), 1816–1825. https://doi.org/10.1111/2041-210X.13669.

(49) Heiple, Z.; Huie, J. M.; Medeiros, A. P. M.; Hart, P. B.; Goatley, C. H. R.; Arcila, D.; Miller, E. C. Many Ways to Build an Angler: Diversity of Feeding Morphologies in a Deep-Sea Evolutionary Radiation. *Biol. Lett.* **2023**, *19* (6), 20230049. https://doi.org/10.1098/rsbl.2023.0049.

(50) Mihalitsis, M.; Bellwood, D. Functional Implications of Dentition-Based Morphotypes in Piscivorous Fishes. *R. Soc. Open Sci.* **2019**, *6* (9), 190040. https://doi.org/10.1098/rsos.190040.

(51) Paradis, E. *Analysis of Phylogenetics and Evolution with R*; Use R!; Springer: New York, 2006.
